# Bioresorbable permanent-magnet beacons for tracking internal biomechanics during free movement

**DOI:** 10.64898/2026.11.07.760898

**Authors:** Zhongyi Nie, Xiang Lin, Ji Wan, Jie Xu, Kedong Wu, Yan Wang, Chong Zhang, Chendong Yang, Ruyue Zhong, Ning Sun, Weitao Man, Yingda Huang, Shaotong Zhang, Denghua Wu, Yiran Wang, Yaozheng Wang, Minki Hong, Yujie Sun, Hongjie Hu, Jahyun Koo, Mengdi Han

## Abstract

Perioperative complications represent a global health burden that can be reduced through timely interventions guided by continuous physiological monitoring. Bioresorbable sensors excel through natural degradation without subsequent surgical extraction, but motion artefacts disrupt their accurate measurement. We synthesize bioresorbable permanent-magnet nanocomposites from aligned iron nanowires as implantable beacons. These beacons intrinsically generate magnetic fields responsive to biomechanical changes, thereby enabling wireless and chipless monitoring. An artificial-intelligence-enhanced wearable magnetic-field camera decodes these spatial maps into calibrated values that tolerate vigorous movement. The system achieves a displacement-equivalent mean absolute error of 2.74 μm. Experiments in rat models demonstrate intracranial pressure tracking with clinical-grade accuracy during vigorous activities, and longitudinal monitoring of intracranial hypertension associated with ischaemic stroke during natural behaviours.

## Main

An estimated 4.2 million people die within 30 days of surgery each year globally^1^. Early detection of perioperative complications can reduce this mortality burden, as pathological changes evolve silently over hours to days even when patients appear stable^2–4^. Implantable sensors can provide continuous monitoring to capture transient, gradual and cumulative information^2,4–7^, as well as activity-dependent abnormalities near the surgical sites^8,9^. Conventional implants^10,11^, however, outlast their clinical utility, as they require secondary retrieval after the perioperative monitoring window^2,12–14^ or impose chronic foreign-body burdens if left in situ^15–18^. Therefore, an ideal perioperative implantable sensor should combine continuous monitoring with complete bioresorption after a predefined monitoring window.

Existing implantable bioresorbable systems function in vivo over predefined operational windows, ranging from hours to weeks, to match the temporary needs of perioperative monitoring^19–25^. These bioresorbable systems often rely on external acoustic^26–28^ or electromagnetic excitations^29^ to achieve wireless sensing without non-resorbable electronics or power sources. However, signal readout depends on stable coupling between the implant and external interrogator. Motion-induced changes in relative position and orientation disrupt this coupling and produce artefacts indistinguishable from genuine physiological changes. These artefacts can cause false alarms or obscure deterioration in freely moving subjects.

Here, we introduce bioresorbable permanent-magnet beacons (BioBeacons) for wireless, chipless, and motion-tolerant tracking of biomechanical signals during natural behaviour. BioBeacons exploit aligned iron nanowires (FeNWs) to confine the rotation of magnetic moment^30^, thereby generating sufficient magnetic fields (MFs) without compromising bioresorbability^31^. The generated quasi-static MFs require no external excitation and penetrate biological tissues without attenuation to enable readout from internal tissues without physical windows or energy transfer losses. A customized wearable MF camera maps the biomechanically altered field, and deep learning decodes these data into motion-tolerant biomechanical values corresponding to a mean absolute error of 2.74 μm, a precision exceeding conventional imaging approaches^32^. In vivo demonstrations across healthy and diseased freely behaving rat models validate the practical utility of the bioresorbable sensing system.

### Bioresorbable hard-magnetic implants paired with wearable MF imaging

The wireless sensing system combines a millimetre-scale, fully bioresorbable BioBeacon with a wearable MF camera positioned on or near the skin surface (Fig. 1a). The compact form factor of BioBeacon facilitates deployment within confined anatomical spaces, including the cranial, thoracic, and abdominal cavities, and BioBeacon undergoes complete degradation and resorption after a predefined monitoring period. Unlike acoustic and high-frequency electromagnetic fields that undergo tissue-dependent attenuation, quasi-static MFs propagate through biological tissues, fluids, bone, and air with negligible loss (Fig. 1b). The wearable camera can therefore capture the spatial MF distribution without requiring direct physical contact or a specific transmission window. The tissue transparency of MF also avoids attenuation-mediated coupling between the transmission path and the measured signal, providing a physical basis for motion-tolerant readout (Supplementary Note 1).

**Fig. 1.**
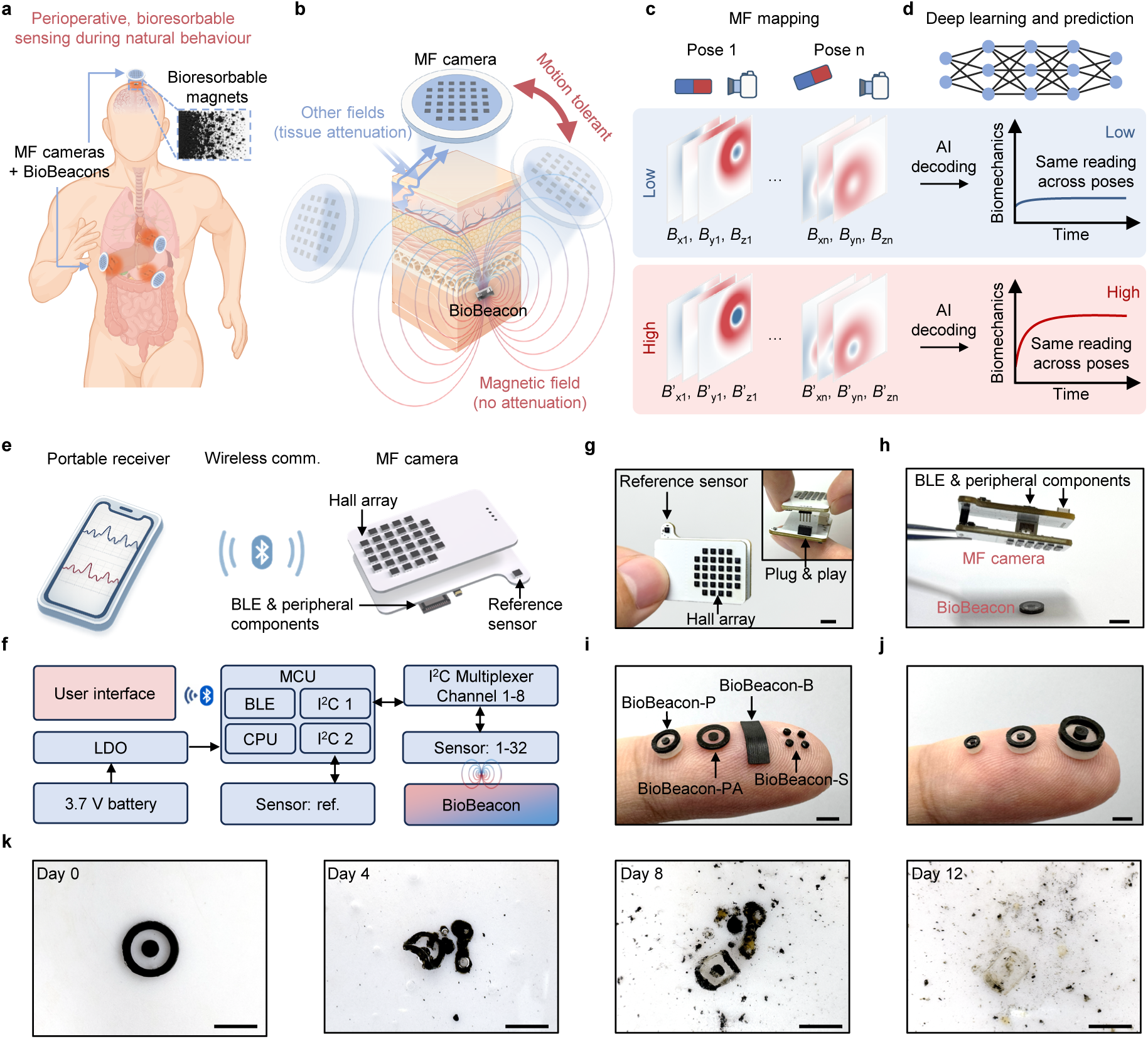
Bioresorbable, motion-tolerant wireless sensing system. **a**, Concept of BioBeacons paired with wearable MF cameras for monitoring biomechanics at different anatomical sites during free motion; inset, bioresorption of BioBeacon. **b**, Illustration of the reflection and absorption of ultrasonic and high-frequency electromagnetic fields from tissues, and the negligible attenuation of quasi-static MFs when passing through biological tissues. This tissue-transparent propagation provides a physical basis for motion-tolerant readout. **c**, Tri-axial MF maps under distinct biomechanical states (rows) and implant–camera poses (columns). **d**, Deep-learning model for decoding the biomechanical signal. **e**, Schematic of the wearable MF camera that wirelessly transmits signals to a portable receiver via BLE; the camera comprises a Hall-sensor array board and a BLE board with a spatially separated reference sensor. **f**, System-level signal flow. **g**, Photograph of the camera; inset, plug-and-play connection between boards. **h**, Representative arrangement of the camera above a BioBeacon without strict alignment. **i**, Configurations for sensing pressure (BioBeacon-P), atmosphere-referenced pressure (BioBeacon-PA), bending/strain (BioBeacon-B), and strain/shear (BioBeacon-S). **j**, Size tunability of BioBeacon-P, with diameters from 2.4 to 6.5 mm. **k**, Disintegration of a BioBeacon during accelerated degradation in phosphate-buffered saline (pH 7.4) at 85 °C over 12 days. Scale bars, 5 mm (**g**,**h**,**k**) and 3 mm (**i**,**j**).

Biomechanical alterations deform BioBeacon and modify the map of magnetic flux density (*B*; Fig. 1c, rows), whereas changes in the relative position and orientation between BioBeacon and the MF camera produce distinct mappings even for the same biomechanical state (Fig. 1c, columns). An end-to-end deep-learning algorithm analyses the measured tri-axial MF distribution and reconstructs the corresponding motion-robust, quantitatively calibrated biomechanical values (Fig. 1d).

The wearable MF camera integrates 32 Hall sensors (corner-truncated 6 × 6 grid; pitch: 2.5 mm) and a Bluetooth Low Energy (BLE) microcontroller that transmits the mapping to a portable receiver (Fig. 1e and Supplementary Figs. 1–4). A reference sensor compensates for common-mode variations in the geomagnetic field (Fig. 1f,g and Supplementary Fig. 5). The external MF camera sits above BioBeacon without the requirement of strict alignment (Fig. 1h) and can scale up for mapping over a larger area (Supplementary Figs. 6 and 7).

BioBeacon can exhibit varied shapes, sizes, and spatial arrangements to accommodate different biomechanical quantities and anatomical constraints (Fig. 1i and Supplementary Table 1). For pressure sensing, the concentric dual-magnet BioBeacon comprises a fixed annular magnet and a cylindrical magnet bonded to a compliant membrane (Supplementary Fig. 8). Applied pressure displaces the cylindrical magnet axially relative to the annular magnet, which changes the spatial MF distribution detected by the camera (Supplementary Fig. 9). A sealed cavity provides an internal pressure reference for fully implanted measurements (BioBeacon-P), whereas a cavity-free configuration uses the atmosphere as an external reference (BioBeacon-PA). For bending and strain sensing (BioBeacon-B), a bioresorbable dynamic covalent polyurethane (b-DCPU) film containing aligned FeNWs undergoes continuous deformation that alters its MF distribution. For strain and shear sensing (BioBeacon-S), a planar b-DCPU membrane carrying four discrete magnets converts tensile strain into rectangular expansion and shear into parallelogram distortion (Supplementary Fig. 10). Dimensions of BioBeacon are also broadly tunable, as exemplified by BioBeacon-P devices ranging from 2.4 to 6.5 mm in diameter (Fig. 1j and Supplementary Fig. 11), and a simplified membrane model guides their design for different applications (Supplementary Figs. 12 and 13 and Supplementary Note 2).

BioBeacon consists entirely of bioresorbable materials: FeNW/poly(ε-caprolactone) (PCL) composites as the magnets, b-DCPU (modulus: ∼2.5 MPa; Supplementary Figs. 14 and 15) as the major deformable components, and pure PCL (modulus: ∼350 MPa) as the rigid supports. An assembled BioBeacon undergoes gradual hydrolysis and structural disintegration over 12 days in phosphate-buffered saline (PBS; pH 7.4) at 85 °C (Fig. 1k).

### Synthesis and characterization of bioresorbable hard magnetic nanocomposites

The bioresorbable magnet exploits FeNWs as its hard magnetic component. Unlike traditional permanent magnets (e.g., NdFeB and SmCo), which are inherently toxic, Fe is a biologically essential element and can degrade and enter physiological metabolic pathways^31^. Bulk Fe consists of multiple magnetic domains, and its magnetization changes readily through domain-wall motion and domain rotation, resulting in relatively low coercivity (∼10 Oe) and remanence (<1 emu/g; Fig. 2a,b)^33,34^. In contrast, confining Fe into nanowires below the critical threshold for domain-wall formation (e.g., ∼140 nm with aspect ratio of 10)^35^ yields single-domain magnets that inherently restrict magnetization reversal^30^ (Supplementary Note 3). The high aspect ratio introduces strong shape anisotropy and establishes the nanowire long axis as the easy magnetization axis, thereby producing higher coercivity and remanence along this direction (Fig. 2c,d and Supplementary Note 3).

**Fig. 2.**
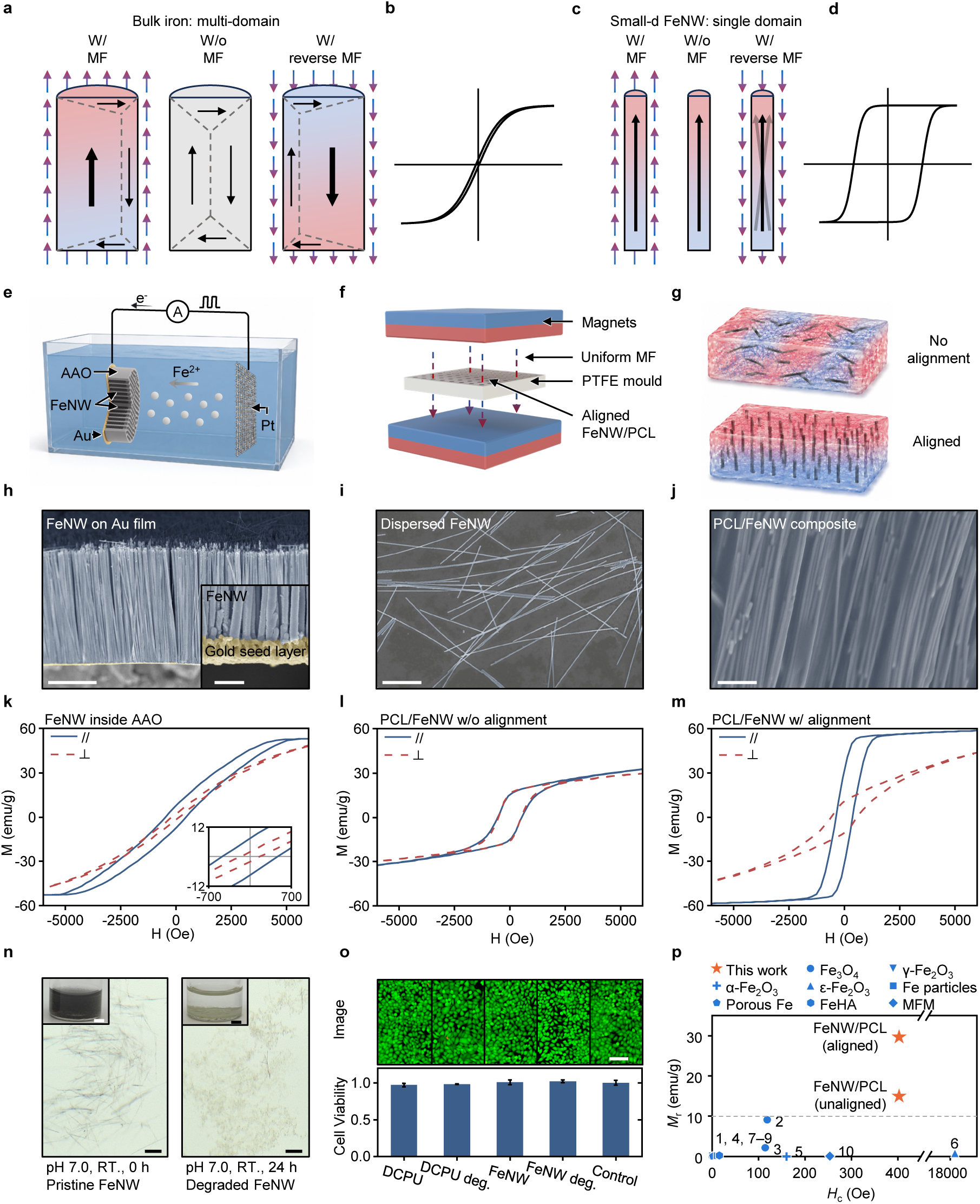
Synthesis and characterization of bioresorbable hard-magnetic nanocomposites. **a**–**d**, Schematics of magnetization reversal and hysteresis in multidomain bulk iron (**a**,**b**) and high-aspect-ratio FeNWs (**c**,**d**). **e**, Pulsed electrodeposition of FeNWs in an Au-coated anodic aluminium oxide (AAO) template. **f**, Field-assisted alignment of FeNWs in PCL solution during casting. **g**, Schematics of FeNW orientations in composites prepared without and with the alignment field. **h**–**j**, Scanning electron micrographs of FeNWs on the Au seed layer after AAO dissolution (**h**; inset, FeNW–Au interface), dispersed FeNWs (**i**), and aligned FeNWs in PCL (**j**). **k**–**m**, Hysteresis loops measured parallel and perpendicular to the nanowire or alignment direction for 60-nm FeNWs in AAO (**k**), unaligned FeNW/PCL (**l**), and aligned FeNW/PCL (**m**). **n**, Optical images of pristine FeNWs (left) and products after 24 hours (right) in phosphate-buffered saline (pH 7.0); insets, macroscopic views. **o**, Live/dead images and viability of L929 cells exposed to b-DCPU, FeNWs, and their degradation products; untreated cells serve as controls. Bars, mean; error bars, s.d. of three independent measurements. **p**, Remanence and coercivity of bioresorbable magnetic composites; numbers correspond to entries in Supplementary Table 2. Scale bars, 10 μm (**h**,**n**), 400 nm (**h**, inset), 4 μm (**i**), 1 μm (**j**), 5 mm (**n**, insets), and 100 μm (**o**).

Synthesis of FeNWs includes sputtering a layer of Au (thickness: ∼350 nm) onto one side of a nanoporous anodic aluminium oxide (AAO) template to act as the working electrode, and subsequent pulsed electrodeposition of Fe within the channels to produce FeNWs (Fig. 2e and Supplementary Figs. 16–18). After selective dissolution of the AAO template and ultrasonic dispersion, scanning electron microscopy (SEM) images reveal the high uniformity and an aspect ratio of 383 for the electrodeposited FeNWs, and elemental mapping confirms the presence of Fe (Fig. 2h and Supplementary Fig. 19).

Transforming FeNWs into a functional, macroscopic component involves their integration within a bioresorbable PCL matrix. The fabrication exploits an MF-assisted alignment strategy to ensure uniform alignment of FeNWs within the polymer, thereby maximizing the macroscopic remanence (*M*_r_). Specifically, the use of a low-viscosity precursor (FeNWs dispersed in a PCL/cyclohexanone solution; PCL concentration: 10% w/v; PCL-to-FeNW mass ratio: ∼2 to 1) facilitates the rotation of nanowires in the MF, while simultaneously enabling facile casting into target geometries with a mould. A uniform MF (∼120 mT) from a pair of parallel NdFeB plate magnets (10 cm × 10 cm × 3 cm; Fig. 2f) exerts torque on the randomly suspended FeNWs (Fig. 2i) during solvent evaporation, and aligns their easy magnetization axes parallel to the field direction before the polymer matrix solidifies (Fig. 2g). Supplementary Fig. 20 shows the detailed fabrication process, alignment setup, and resulting FeNW/PCL nanocomposite magnets with different geometries. Optical microscopy shows that FeNWs dispersed in polyvinylpyrrolidone (PVP)/ethanol adopt random orientations without an external MF but preferentially align upon application of the field, demonstrating their field-responsive rotational behaviour (Supplementary Fig. 21a,b and Supplementary Video 1). After incorporation into the PCL matrix, SEM images reveal a highly ordered FeNW microstructure in the MF-aligned composite, in sharp contrast to the disordered arrangement observed in the unaligned control (Fig. 2j and Supplementary Fig. 21c,d).

Magnetic hysteresis loops of FeNWs (diameter: 60 nm) retained in the AAO template show high remanence and coercivity (*M*_r_: ∼7 emu/g; *H*_c_: ∼500 Oe) along the nanowire axis, together with pronounced axial magnetic anisotropy (Fig. 2k). Consistent with the theoretical size dependence (Supplementary Note 3), thinner FeNWs (diameter: 30 nm) exhibit enhanced coercivity and remanence (*M*_r_: ∼12 emu/g; *H*_c_: ∼750 Oe) compared with their 60-nm counterparts (Supplementary Fig. 22). Both remanence values are normalized to the total sample mass, including the AAO template. Because the 60-nm FeNWs experience larger magnetic torque, which facilitates alignment in PCL/cyclohexanone, they serve as the magnetic material for the FeNW/PCL nanocomposite. When the FeNWs are incorporated into PCL without MF-assisted alignment, their random orientations partially average out the directional magnetic responses, resulting in lower macroscopic remanence (*M*_r_: ∼15 emu/g; *H*_c_: ∼400 Oe) and weaker directional dependence (Fig. 2l). By contrast, MF-assisted alignment increases the macroscopic remanence and loop squareness along the alignment direction (*M*_r_: ∼30 emu/g; *H*_c_: ∼400 Oe; Fig. 2m), and produces pronounced magnetic anisotropy in the composite (*M*_r_: ∼10 emu/g; *H*_c_: ∼500 Oe perpendicular to the alignment direction). These results validate the effectiveness of the alignment strategy for constructing bioresorbable permanent magnet with pronounced anisotropy.

Further degradation and cytocompatibility tests confirm the suitability of using FeNW-based materials as bioresorbable implants. When incubated in PBS (pH: 7.0), FeNWs (diameter: 60 nm) undergo rapid morphological disintegration within 24 h, transforming from nanowires into fine light-yellow precipitates (Fig. 2n and Supplementary Figs. 23 and 24). This visible structural transition confirms the degradability of FeNWs under a physiologically relevant buffered environment. X-ray diffraction (XRD) further confirms the degradability. In contrast to the distinct body-centred cubic α-Fe peak (2θ: ∼44.7°) observed in the pristine FeNWs, the light-yellow precipitates obtained after degradation show no distinct diffraction peaks, indicating the disappearance of the crystalline α-Fe phase and the formation of poorly crystalline or amorphous degradation products (Supplementary Fig. 23j). CCK-8 assays using both FeNW dispersions (100 μg/mL) and FeNW degradation solutions (5 μg/mL) show high cell viability over the tested concentrations, indicating limited cytotoxicity of the pristine and degraded FeNW products. Tests of b-DCPU (3 cm^2^/mL) and its degradation products (0.3 cm^2^/mL) yield similar results (Fig. 2o and Supplementary Fig. 25). Live/dead staining further confirms predominantly viable cells after exposure to these materials and their degradation products, consistent with the CCK-8 results (Fig. 2o and Supplementary Fig. 26). Compared with other macroscopic magnetic structures made from bioresorbable magnetic materials, the FeNW/PCL nanocomposite exhibits more than threefold higher remanence while retaining substantial coercivity (Fig. 2p, Supplementary Table 2 and Supplementary Note 4). These results demonstrate that FeNW-based composites provide the required combination of hard magnetic performance, degradability, and biological compatibility for bioresorbable implants.

### Motion-tolerant wireless sensing via MF mapping and deep learning

During freely moving operation, the spatial MF distributions measured by the wearable camera (Δ*B*_all_) vary with three factors: pose-dependent variations in the geomagnetic background (Δ*B*_geo_), changes in the relative position/orientation between BioBeacon and the MF camera (Δ*B*_rel_), and target biomechanical signals that alter the internal configuration of BioBeacon (Δ*B*_bio_; Fig. 3a). Motion-tolerant sensing therefore requires compensation for geomagnetic variations and separation of biomechanical information from changes induced by relative motion.

**Fig. 3.**
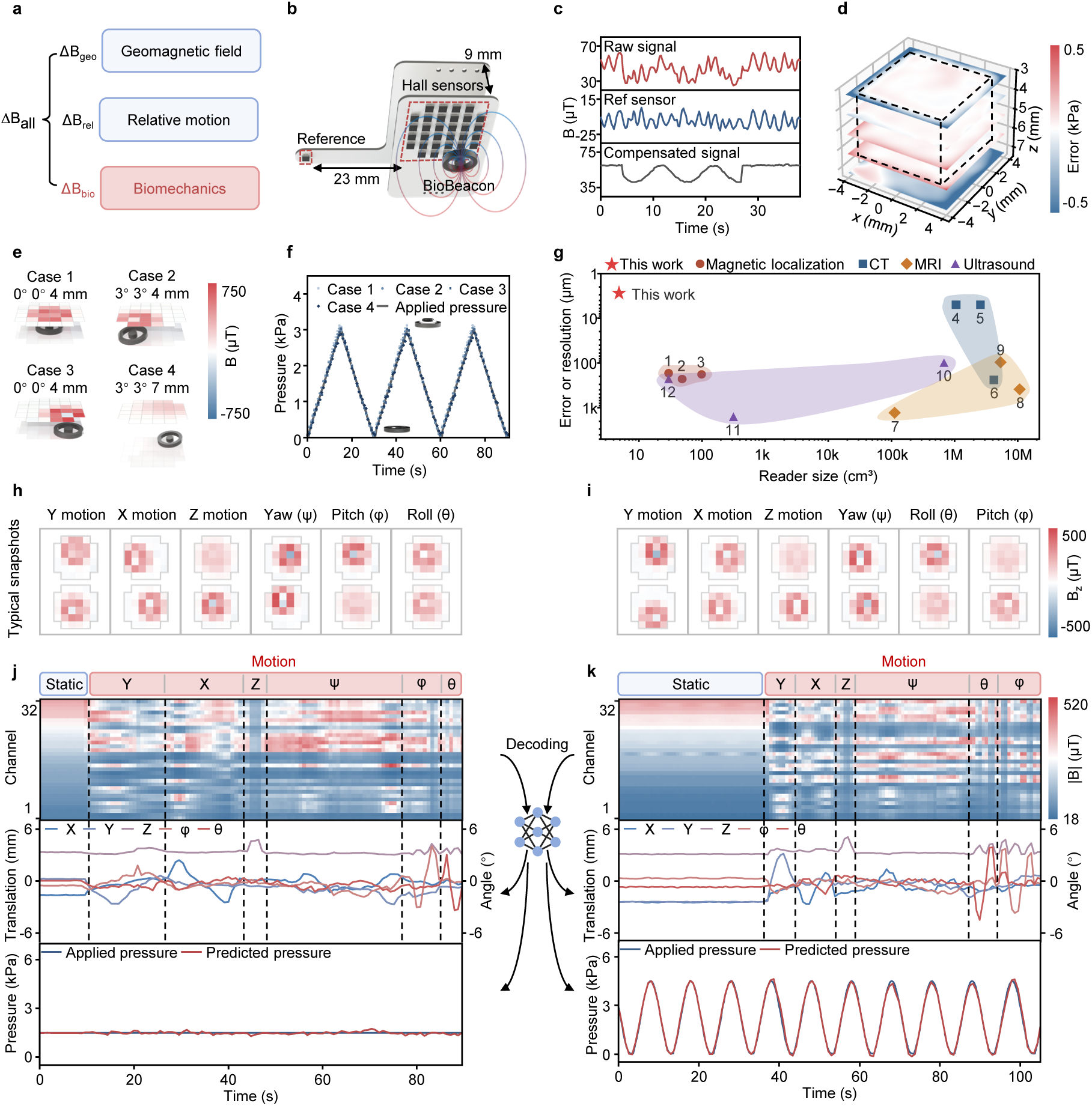
Motion-tolerant MF mapping and quantitative decoding of biomechanical signals. **a**, Schematic of the sources of measured MF changes: geomagnetic background, implant–camera motion, and target biomechanics. **b**, Camera layout with the reference sensor. **c**, Raw, reference, and compensated signals during simulated physiological variation with common-mode magnetic disturbance. **d**, Spatial map of pressure-decoding error across an extended workspace; dashed cuboid, trained region. **e**, MF maps for four camera–implant configurations. **f**, Ground-truth and decoded pressure for these configurations during triangular inputs. **g**, Comparison of the MF camera with magnetic-localization and clinical imaging systems in reader size and spatial precision. Numbers correspond to entries in Supplementary Table 3. **h**,**i**, MF snapshots during translations and rotations about three axes under constant (**h**) and dynamic (**i**) pressure. **j**,**k**, Multichannel MF signals (top), reconstructed five-dimensional poses (middle), and decoded pressure (bottom) during sequential multi-axis motion at 1.5 kPa (**j**) and under sinusoidal pressure (**k**); dashed lines separate motion stages.

To compensate for geomagnetic variations during untethered operation, the reference sensor is positioned laterally 23 mm and vertically 9 mm from the Hall sensor array, where the MF generated by BioBeacon decreases to <2 μT (comparable to the sensor noise), to track geomagnetic variations (Fig. 3b and Supplementary Fig. 27). Reference subtraction removes the common-mode variations and reveals the simulated physiological signal (Fig. 3c and Supplementary Note 5).

The multi-magnet architecture of BioBeacon provides a physical basis for separating biomechanical signals from pose-induced variations. In BioBeacon-PA, pressure changes the internal relative configuration of its two magnets, whereas rigid-body motion changes their collective pose relative to the MF camera. Because the MF camera captures the spatial superposition of the fields generated by both magnets, these two processes produce distinguishable spatial MF signatures (Supplementary Fig. 28 and Supplementary Note 6). Compared with analytical models that rely on idealized material properties and device geometries, a data-driven approach directly learns from measured MF distributions and thereby accounts for material inhomogeneities and shape imperfections (Supplementary Note 7). A 6-axis robotic arm moves the MF camera through predefined positions and orientations while a pressure controller applies different pressure levels to BioBeacon-PA, which generates a pose–pressure dataset for model training (Supplementary Fig. 29 and Supplementary Note 8).

On an independent validation set comprising previously unseen poses within the calibrated workspace, the decoded pressures of the trained model (Supplementary Fig. 30) closely match the ground-truth values, with an *R*^2^ of 0.9994 and a mean absolute error (MAE) of 0.031 kPa (Supplementary Fig. 31). The spatially resolved decoding errors remain low within the calibrated workspace (MAE: 0.034 kPa) and increase beyond its boundaries (MAE: 0.35 kPa; Fig. 3d). Expanding the calibration range enlarges this reliable workspace (Supplementary Fig. 32 and Supplementary Note 9). The sensor spacing and number also govern the decoding error (Supplementary Fig. 33). Across lateral displacement, increased vertical separation, and angular tilting between the MF camera and BioBeacon-PA (Fig. 3e), the decoded pressure traces closely follow the ground-truth signals under triangular, sinusoidal, and square pressure inputs (Fig. 3f and Supplementary Figs. 34 and 35). Pointwise conversion of the validation-set pressures through the calibrated pressure–displacement curve of BioBeacon-PA yields an equivalent displacement MAE of 2.74 μm (Supplementary Fig. 36 and Supplementary Note 10). Compared with conventional magnetic-localization and medical-imaging systems, the MF camera exhibits a more compact form factor and higher spatial precision in resolving subtle implant deformations (Fig. 3g and Supplementary Table 3).

Additionally, the system preserves accuracy during continuous dynamic motions that emulate real-world freely behaving scenarios (Supplementary Videos 2 and 3). In the first protocol, the MF camera sequentially undergoes multi-axis translations and rotations under a static pressure of 1.5 kPa. These motions disrupt the spatial maps (Fig. 3h) and channel-wise temporal output (top of Fig. 3j). Nevertheless, the decoded pressure remains close to the ground-truth value (MAE: 0.065 kPa; bottom of Fig. 3j), and the model also reconstructs the imposed translations and tilts (middle of Fig. 3j). In the second protocol, under sinusoidal pressure with similar motions (Fig. 3i), the decoded pressure continues to closely track the ground-truth pressure (*R*^2^: 0.9948; MAE: 0.086 kPa; Fig. 3k). These results demonstrate the ability to recover true dynamic pressure signals when pressure variations and motion artefacts occur simultaneously, which enables continuous physiological monitoring without immobilization or anaesthesia.

Reconfigurable magnetic layouts extend MF mapping to other biomechanical deformations. Four discrete magnets in the BioBeacon-S layout bonded directly to an ex vivo rabbit stomach use the tissue itself as a deformable substrate, which minimizes constraints on its native deformation. Gastric expansion increases the inter-magnet distances and shear changes the angles of the initially square layout, which appears as an expanded and tilted quadrilateral in the MF map (Supplementary Fig. 37a). For BioBeacon-B imaged by a 256-sensor MF camera, bending under through-thickness magnetization shifts the smoothed boundary between positive and negative *B*_z_ regions in accordance with the bent profile (Supplementary Fig. 37b), and tensile deformation under longitudinal magnetization alters the positions and shapes of these field regions (Supplementary Fig. 37c and Supplementary Note 11).

### In vivo validation of wireless ICP monitoring in healthy rats

Intracranial pressure (ICP) is a key physiological parameter for detecting evolving intracranial hypertension in traumatic brain injury^6,36^, stroke-associated cerebral oedema^37,38^ and hydrocephalus^39^. Continuous monitoring can capture both rapid pressure elevations and gradual changes to guide timely intervention^5–7,36,37^. In vivo measurements in healthy rats demonstrate the clinical-grade accuracy and motion tolerance of BioBeacon. After craniotomy, BioBeacon is placed epidurally over the burr hole and sealed with dental cement, and the MF camera is mounted on the head surface (Fig. 4a and Supplementary Fig. 38). The sealed space mechanically couples to ICP without compressing the underlying brain tissue. Subcutaneously injected FeNWs show a marked reduction in visible FeNWs from day 1 to day 279, which provides evidence of gradual bioresorption in vivo (Fig. 4b). Thirty days after implantation, haematoxylin and eosin (H&E) staining of tissues near the implantation site and major visceral organs reveals preserved tissue architecture and normal cellular morphology, and Masson’s trichrome staining shows no obvious excess collagen deposition or fibrosis compared with the control group (Fig. 4c and Supplementary Figs. 39 and 40). These results support the bioresorbability as well as the local and systemic biocompatibility of BioBeacon.

**Fig. 4.**
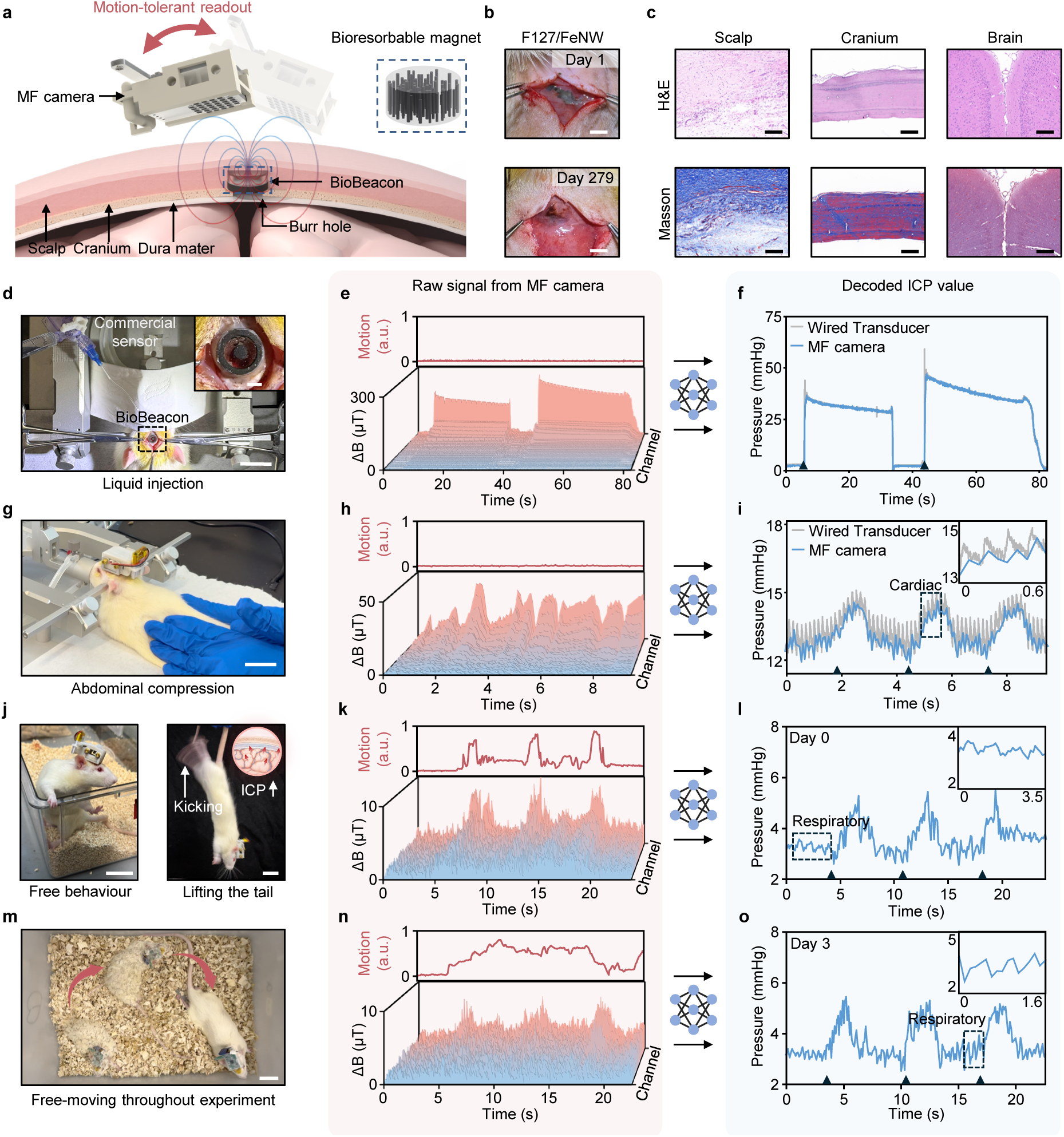
In vivo validation of motion-tolerant wireless ICP monitoring. **a**, Schematic of BioBeacon over a cranial burr hole, with the head-mounted MF camera maintaining readout during relative motion; inset, FeNW/PCL magnet. **b**, Images of subcutaneous FeNWs in Pluronic F127 on days 1 and 279. **c**, H&E (top) and Masson’s trichrome (bottom) staining of scalp, cranium, and brain 30 days after implantation. **d**, Photograph of wireless ICP monitoring during liquid injection in an anaesthetized rat; inset, implanted BioBeacon. **e**,**f**, Normalized motion intensity (upper trace in **e**), geomagnetically compensated multichannel MF signals (lower plot in **e**), and decoded ICP with a wired commercial transducer (**f**) during liquid injection. **g**, Photograph of abdominal compression. **h**,**i**, Same as in **e** and **f**, but during abdominal compression; inset in **i**, cardiac pulsations. **j**, Photographs of free movement (left) and tail lifting (right) in an awake rat on postoperative day 0. **k**,**l**, Motion intensity and compensated multichannel MF signals (**k**) and decoded ICP (**l**); inset in **l**, respiratory fluctuations. **m**, Superimposed photographs of free movement. **n**,**o**, Same as in **k** and **l**, but on postoperative day 3. Triangles in **f**, **i**, **l** and **o** mark perturbations. Scale bars, 5 mm (**b**), 200 μm (**c**), 2 cm (**d**), 2 mm (**d**, inset), and 3 cm (**g**,**j**,**m**).

Comparison with a tethered commercial pressure transducer under anaesthesia validates the accuracy of BioBeacon in ICP monitoring. Controlled liquid injection generates ICP elevations up to 50 mmHg (Fig. 4d–f), whereas manual abdominal compression induces rapid transient pressure changes (Fig. 4g–i). In both cases, the normalized motion intensity derived from the reference sensor remains low, but the multichannel signals exhibit pronounced, channel-dependent responses during the ICP perturbations (Fig. 4e,h). The decoded pressure traces exhibit close agreement and precise temporal synchronization with the commercial reference (Fig. 4f,i), with a mean absolute error below 0.5 mmHg that satisfies the ANSI/AAMI clinical requirement for ICP monitoring (<2 mmHg at 0–20 mmHg and <10% at 20–100 mmHg)^40^. BioBeacon also detects subtle cardiac-induced pressure pulses with the same dominant frequency as the commercial reference (6.56 Hz, ∼394 beats per minute; Fig. 4i, inset, and Supplementary Fig. 41).

Tail-lifting tests in awake, freely moving rats verify the capability of the system for continuous monitoring (Fig. 4j,m and Supplementary Fig. 42). The inverted body posture (head-down) induces an immediate elevation in ICP through the gravity-induced redistribution of blood toward the cranium. These manoeuvres and spontaneous animal movement introduce substantial geomagnetic variations, as indicated by the motion intensity (Fig. 4k,n and Supplementary Fig. 43). After reference compensation, the multichannel signals still exhibit pronounced variations in response to both pressure changes and residual relative motion. In contrast, the decoded ICP traces consistently capture the pressure elevations due to tail lifting from day 0 to day 3 after implantation and return to a stable baseline of ∼3.5 mmHg despite free movement (Fig. 4l,o and Supplementary Video 4). Respiratory-related pulsations also remain detectable in the decoded ICP throughout this period (Fig. 4l, inset), which confirms the multi-day functional stability of the system under freely moving conditions.

### Continuous monitoring of ICP dynamics in a freely behaving rodent stroke model

Ischaemic stroke provides a clinically important example in which evolving cerebral oedema can cause delayed ICP elevation. Focal ischaemia and reperfusion induce cerebral oedema formation^41^, and severe space-occupying oedema after middle cerebral artery infarction can lead to neurological deterioration and life-threatening intracranial hypertension^37,38^. Patients in such cases may require decompressive hemicraniectomy to reduce mortality risk^42^. Therefore, early detection and longitudinal tracking of oedema-related ICP elevation over multiple days are important for guiding timely clinical intervention^38,43^.

This clinical scenario demands the MF camera to maintain high measurement precision and stable energy supply. A resonant wireless power-transfer system is constructed by winding a transmitter coil (15 cm × 28 cm, 2 turns) around the housing cage to envelop the behavioural area with a continuous alternating MF (Fig. 5a). A compact receiver coil integrated into the wearable circuitry harvests this energy by inductive coupling. Importantly, wireless power is delivered exclusively to the external MF camera, whereas the bioresorbable BioBeacon remains entirely passive and excitation-free and is detected solely through its MF (Fig. 5b). Impedance matching of the receiver coil maximizes the harvesting efficiency (matched resonance at 13.56 MHz; impedance, 7.09 kΩ; phase angle, 0.06°; Fig. 5c). The wirelessly powered MF camera supports sensing modes with different precision and energy requirements. Increasing the number of sampling repetitions suppresses measurement noise but increases the energy consumption per sample, providing a tunable trade-off between measurement precision and power consumption (Fig. 5d). Real-time monitoring of the battery voltage shows rapid depletion in the battery-powered mode, whereas the wirelessly powered system maintains a stable voltage during continuous high-precision monitoring (Fig. 5e). This strategy ensures long-term operation on the external side and preserves the passive, battery-free, and fully bioresorbable nature of the internal implant.

**Fig. 5.**
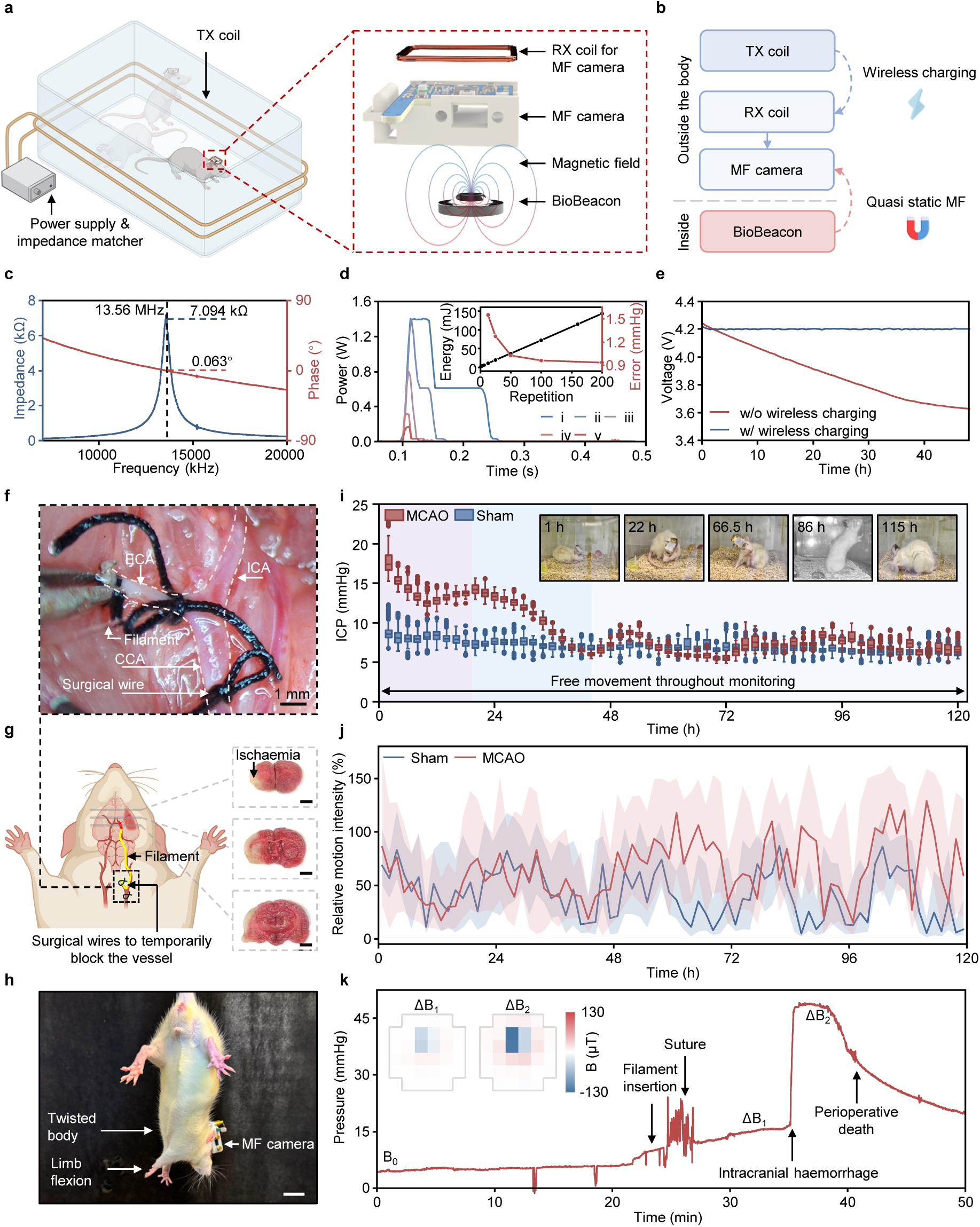
Wireless powering of the MF camera and longitudinal ICP monitoring after stroke. **a**, Schematic of the transmitter coil around the home cage and receiver coil on the camera. **b**, Illustration of separate power (inductive) and sensing (quasi-static MF) pathways; BioBeacon remains passive. **c**, Receiver-coil impedance and phase, showing resonance at 13.56 MHz. **d**, Power–time profiles for repetition settings i to v (X/Y- and Z-axis repetitions of 160/200, 50/63, 13/16, 3/4, and 1/1); inset, energy per measurement and decoding error. **e**, Battery voltage over ∼48 hours with and without wireless charging. **f**, Surgical view of filament placement for t-MCAO. **g**, Schematic of transient occlusion and TTC-stained sections showing infarction. **h**, Neurological deficits in a rat carrying the camera after t-MCAO. **i**, ICP box plots in 2-hour bins over ∼120 hours in a t-MCAO (red) and a sham-operated (blue) rat; boxes, interquartile range (IQR); centre lines, medians; whiskers, 1.5 × IQR; diamonds, outliers (n = 295–300 and 292–300 per box, respectively). Insets, representative video frames of the t-MCAO rat at the indicated time points during unrestrained monitoring. **j**, Relative motion intensity of the same rats; lines, medians in 1.5-hour bins; shading, IQR (n = 44–45 per bin). **k**, Perioperative ICP trace from an unsuccessful t-MCAO procedure; insets, MF maps at two stages. Scale bars, 1 mm (**f**), 2.5 mm (**g**), and 2 cm (**h**).

A transient middle cerebral artery occlusion (t-MCAO) rat model simulates severe ischaemic stroke and reperfusion injury^44^: a filament obstructs the blood flow for 1 h, followed by filament removal to restore perfusion (Fig. 5f,g). Behavioural assessments indicate successful model establishment, as postoperative rats exhibit characteristic neurological motor deficits, including asymmetric body twisting and forelimb flexion during tail suspension (Fig. 5h) and spontaneous circling during free locomotion (Supplementary Video 5)^44–46^. 2,3,5-Triphenyltetrazolium chloride (TTC) staining^47^ of brain sections harvested after the monitoring period (120 h) further confirms cerebral infarction, which is absent in the sham-operated rat (Fig. 5g and Supplementary Fig. 44).

Continuous ICP recordings from the t-MCAO rat and the sham-operated rat under freely moving conditions reveal distinct postoperative temporal profiles (Fig. 5i). During the early postoperative phase (0–24 h), ICP in the t-MCAO rat remains elevated relative to the sham-operated rat, consistent with evolving post-ischaemic cerebral oedema and other intracranial volume changes. From 24 to 48 h, ICP gradually declines, which potentially reflects partial oedema resolution and intracranial compensation^48–50^. From 48 to 120 h, ICP remains comparatively stable near the sham-operated range. Mean ICP during 0–48 h in the t-MCAO rat (mean ± s.d.: 11.54 ± 3.20 mmHg) is ∼1.5 times higher than that in the sham-operated rat (7.57 ± 0.81 mmHg), and this difference narrows during the recovery period (48–120 h; 7.06 ± 0.98 versus 6.69 ± 0.64 mmHg). These distinct ICP profiles demonstrate the ability of the system to resolve the temporal evolution of intracranial hypertension associated with ischaemic stroke.

Throughout the monitoring period, both rats remain unrestrained, and the relative motion intensity, calculated from changes in the reference-MF vector, provides a continuous measure of their spontaneous activity (Fig. 5j). The sham-operated rat exhibits a pronounced day–night activity pattern (mean motion intensity: 43.49% during the daytime versus 63.04% at night), whereas the t-MCAO rat shows a less pronounced pattern (73.00% versus 78.35%) following ischaemic stroke (Supplementary Note 12), consistent with disrupted activity patterns after stroke^51^.

Beyond this multi-day pathological trajectory, the system also captures ICP dynamics at finer temporal scales. In the t-MCAO rat, breathing-related ICP fluctuations are distinguishable across different respiratory states, with amplitudes increasing from ∼0.3 mmHg under deep anaesthesia to ∼2 mmHg during laboured breathing (Supplementary Fig. 45 and Supplementary Video 6). Data from two aborted t-MCAO surgeries show abrupt ICP elevations following filament manipulation and before perioperative death (Fig. 5k and Supplementary Fig. 46). Such high ICPs reflect acute intracranial complications, and post-mortem identification of an intracranial haematoma in one subject indicates surgery-induced cerebral haemorrhage. These findings demonstrate the value of continuous perioperative monitoring for capturing acute intraoperative complications with high temporal resolution and tracking evolving postoperative abnormalities over extended periods.

## Discussion

This work establishes a fully bioresorbable and motion-tolerant system for continuous perioperative biomechanical monitoring during natural behaviour. A bioresorbable nanocomposite of FeNWs and polymers form the BioBeacon that require no power, electronics, wiring, or excitation. Reference-based compensation and deep-learning-based decoding enable robust sensing despite dynamic changes in the relative position and orientation of the BioBeacon and MF camera. These materials, device architecture and data-analysis strategies together address key challenges in bioresorbable implantable monitoring by distinguishing physiological signals from motion artefacts, and enable continuous sensing in freely behaving conditions even when the implants and readers move relative to each other. In vivo validations in healthy and diseased rodent models demonstrate the capabilities to capture transient events and gradual disease progression without prolonged anaesthesia or wired instrumentation.

Existing bioresorbable sensing systems based on LC resonant, ultrasonic, or optical interrogation remain susceptible to motion-dependent changes in implant–reader distance and orientation. In contrast, the fully bioresorbable BioBeacon, paired with the wearable MF camera, preserves measurement accuracy during free behaviour even when body movements shift or rotate the reader with respect to the implant (Supplementary Fig. 47, Supplementary Table 4, Supplementary Note 13). Future developments could enhance the spatial resolution of the external MF camera to improve sensing precision, extend the calibrated operating range to withstand motions with larger amplitudes, develop adaptive decoding methods to accelerate the calibration, and incorporate responsive materials to expand the sensing modality. These advances offer a solution for temporary, untethered, motion-tolerant monitoring during natural behaviour throughout critical perioperative periods, without long-term burden and additional removal procedures associated with conventional implants.

## Methods

### Synthesis and purification of FeNWs

#### Substrate Preparation

A magnetron sputtering system deposited a Au film onto one side of a commercial porous AAO template (25 mm diameter, 60 nm pore diameter, 125 nm interpore distance; DP125-060-50000, Topmembranes) to create the conductive substrate. For this process, the system first evacuated the sputtering chamber to a base pressure of 1.5 × 10^-4^ Pa. An argon (Ar) gas flow of 50 sccm maintained a working pressure of 0.5 Pa. The power supply applied 100 W of DC power to the Au target. Following a 60 s pre-sputtering step to clean the target, the sputtering process continued for 600 s on the AAO template. This procedure yielded an estimated Au film thickness of ∼350 nm.

#### Electrolyte Preparation

Preparation of the electroplating bath involved dissolving 90 g of iron(II) sulfate heptahydrate (FeSO₄·7H₂O), 33.75 g of boric acid (H₃BO₃), 12.75 g of ammonium sulfate ((NH₄)₂SO₄), and 0.75 g of L-ascorbic acid in deionized water. The addition of deionized water brought the solution to a final total volume of 800 mL, followed by stirring until all components fully dissolved.

#### Electrochemical Deposition of FeNWs

The electrochemical setup consisted of a two-electrode system. The Au-sputtered AAO template served as the working electrode (cathode), and a 40 mm × 40 mm platinum (Pt) mesh acted as the counter electrode (anode). A programmable current source (6221, Keithley) applied a pulsed current for a total of 3 hours at room temperature. The pulse, a 10 Hz square wave with a 10% duty cycle, alternated between a cathodic current of 100 mA for 10 ms (ON-time) and 0 mA for 90 ms (OFF-time).

#### Release and Dispersion of FeNWs

The nanowire release process began with immersing the entire AAO template in a 20% (w/v) sodium hydroxide (NaOH) solution at room temperature to selectively etch the alumina matrix. This step resulted in a flexible composite film comprising the FeNW array still attached to the gold substrate. Repeated washing of this composite film with deionized water continued until the supernatant reached a neutral pH (pH = ∼7). Two additional ethanol washes followed the water rinse. The next step involved transferring the washed film to a beaker containing 30 mL of a 2% (w/v) PVP (Mw 40000; S30268, Yuanye) ethanol solution. A 650 W probe sonicator (JC-Y92-IIN, Lichen) operating in an ice bath then detached and dispersed the FeNWs from the gold film. The sonicator operated at 15% power (97.5 W) in a pulse mode (6 s ON, 3 s OFF) for a total sonication time of 45 s. This process yielded a dispersion containing both the iron nanowires and the fragmented gold substrate, suspended in the PVP/ethanol solution. A centrifuge then spun the dispersion at 4000 rpm for 5 s. This brief centrifugation sedimented the dense Au fragments at the bottom of the centrifuge tube, while the well-dispersed FeNWs remained in the supernatant. A pipette carefully collected the FeNW-containing supernatant and transferred it into a clean vial, yielding a purified FeNW dispersion in PVP/ethanol. The fabrication and purification of 30-nm-diameter FeNWs followed the same protocol, except that an AAO template with a pore diameter of 30 nm and an interpore spacing of 65 nm replaced the original template.

### Synthesis of the bioresorbable dynamic polyurethane (b-DCPU) membrane

The synthesis began with adding 10 mL of butyl acetate (287725, Sigma-Aldrich) and 1.35 g of Polycaprolactone (PCL) triol (PCL-triol; average Mn ∼830 g/mol; DZ00002, Xushuo) in a glass vial heated at 60 °C, followed by mixing them until full dissolution. Adding 400 μL of hexamethylene diisocyanate (HDI; 52649, Sigma-Aldrich) as the crosslinking agent and 3.35 μL of tin(II) 2-ethylhexanoate (Sn(Oct)_2_; S3252, Sigma-Aldrich) as the biocompatible catalyst and mixing the solution again until full dissolution (we call the solution here the liquid precursor of the b-DCPU), followed by drop-casting on a silicon wafer yielded evenly-distributed liquid films. The wafers were placed for 12 h on a hotplate at 65 °C for solvent evaporation and curing, resulting in a thin film of around 40 μm. Preparing a 1 mol/L NaCl solution in a petri dish and setting the voltage of a voltage source to 24 V, followed by clamping the silicon wafer to the positive electrode of the voltage source and immersing it in the sodium chloride solution along with the negative electrode for electrolysis enabled the peeling off of b-DCPU film from the silicon wafer after ∼30 s. For b-DCPU structures with other thicknesses or geometries, the liquid precursor filled moulds with the desired shapes and dimensions. Heating the moulds at 65 °C for 12 h promoted solvent evaporation and polymer curing. When curing shrinkage reduced the b-DCPU volume below the target volume, additional precursor filled the remaining space, followed by another curing cycle at 65 °C. Repeating the filling and curing steps allowed the final b-DCPU structures to reach the desired shapes and dimensions.

### Fabrication of bioresorbable FeNW/PCL magnet

#### Preparation of magnetically aligned FeNW/PCL sheets

First, mixing PCL pellets (MB2453-1, Meilunbio), cyclohexanone (M0113451, Energy Chemical), and a pre-dispersed FeNW suspension (PCL concentration in cyclohexanone, 10% w/v; PCL-to-FeNW mass ratio, ∼2:1) in a centrifuge tube produced a homogeneous liquid precursor slurry. The slurry then filled a shallow rectangular polytetrafluoroethylene (PTFE) mould, whose dimensions matched the target device geometry. Before complete solvent evaporation, the mould entered a magnetic alignment setup while the slurry retained sufficient fluidity. Two opposing NdFeB permanent magnets (10 cm × 10 cm × 3 cm) generated an approximately uniform magnetic field of 120 mT around the mould. The external field exerted magnetic torque on the FeNWs and gradually rotated their long axes toward the prescribed field direction. Continued solvent evaporation solidified the PCL matrix and locked the aligned FeNWs within the polymer network, yielding a rectangular FeNW/PCL sheet with a uniform internal nanowire orientation.

#### Fabrication of annular bioresorbable magnets

For the fabrication of annular magnets (representative example: an annular magnet with an outer diameter of 6.5 mm, an inner diameter of 4.9 mm, and a height of 1.0 mm), the FeNW/PCL slurry first filled a rectangular PTFE groove measuring 18 mm × 4 mm × 200 μm. A magnetic field of ∼120 mT aligned the FeNWs across the 4 mm width of the sheet. After complete solidification of the PCL matrix, removal from the mould yielded an aligned rectangular sheet. A blade cut the sheet parallel to its 18 mm length into four strips, each measuring ∼18 mm × 1 mm × 200 μm. Stacking the four strips produced a total thickness of 0.8 mm. A rectangular PTFE groove measuring 18 mm × 1 mm × 0.8 mm held and compressed the stacked strips. Heating at 85 °C thermally fused the four layers into an integrated strip measuring ∼18 mm × 1 mm × 0.8 mm. This procedure preserved the FeNW orientation across the 1 mm width of the strip. Gentle heating to ∼55 °C increased the flexibility of the laminated strip and facilitated its insertion into a flexible PDMS annular-groove mould with an outer diameter of 6.5 mm, an inner diameter of 4.9 mm, and a height of 1.0 mm. Heating the mould at 85 °C for 10 min thermally joined the two ends of the strip and formed a continuous ring. Cooling and demoulding yielded the final annular FeNW/PCL magnet. After demoulding, an external magnetic field magnetized each FeNW/PCL magnet along its height direction, corresponding to the axial direction of the ring. The same procedure produced rings with other dimensions after corresponding adjustments to the sheet, strip, and mould dimensions.

#### Fabrication of cylindrical bioresorbable magnets

For the fabrication of cylindrical magnets (representative example: a cylindrical magnet with a diameter of 2.0 mm and a height of 1.0 mm), the FeNW/PCL slurry first filled a rectangular PTFE groove measuring 16 mm × 5 mm × 200 μm. A magnetic field of ∼120 mT aligned the FeNWs across the 5 mm width of the sheet. After complete solidification of the PCL matrix, removal from the mould yielded an aligned rectangular sheet. Tightly rolling the sheet along its 16 mm length formed a coiled structure with a height of 5 mm. A flexible PDMS mould containing a cylindrical cavity with a diameter of 2.0 mm and a depth of 5.0 mm held and compressed the coiled sheet. Heating at 85 °C for 10 min thermally fused the adjacent layers and produced an integrated cylinder with a diameter of 2.0 mm and a height of 5.0 mm. This rolling geometry maintained the FeNW alignment along the cylinder axis. After cooling and demoulding, a blade cut the cylinder perpendicular to its axis at 1.0 mm intervals, yielding cylindrical FeNW/PCL magnets with a diameter of 2.0 mm and a height of 1.0 mm. After demoulding, an external magnetic field magnetized each FeNW/PCL magnet along its height direction, corresponding to the axial direction of the cylinder. The same procedure produced cylinders with other dimensions after corresponding adjustments to the sheet and mould dimensions.

#### PCL/NdFeB magnets for in vitro validation

To facilitate device preparation and benchtop validation, the *in vitro* experiments in Fig. 3h–k and Supplementary Fig. 37a–c used geometrically identical PCL/NdFeB (MQFP-B, Magnequench) composite magnets with a slightly lower surface magnetic flux density (∼5 mT for the Φ2 mm cylinder, ∼3 mT for the annular, ∼3 mT for the BioBeacon-B). The fabrication followed the procedures described above but used a PCL/NdFeB mixture (weight ratio, 1:1) and a commercial magnetizer (MA3030, Jiujuok; magnetizing voltage, ∼0.8 kV) for magnetization. All *in vivo* experiments in Figs. 4 and 5 used bioresorbable FeNW/PCL magnets.

### Interfacial bonding and assembly of BioBeacons

#### Bonding between PCL-containing components

Solvent-assisted bonding joined two neat-PCL components or an FeNW/PCL magnet and a neat-PCL frame. A small amount of an organic solvent, such as cyclohexanone, chloroform, or dichloromethane, wetted both mating surfaces. The solvent partially dissolved the interfacial PCL and increased the mobility of the surface polymer chains. Immediate alignment and compression brought the two surfaces into close contact. Holding the assembly at room temperature for 1 h allowed solvent evaporation and polymer-chain interdiffusion, thereby promoting physical fusion and forming a stable interface.

#### Bonding between PCL-containing components and b-DCPU films

An in-situ precursor-curing strategy joined an FeNW/PCL magnet or a neat-PCL component to a b-DCPU film. Application of a small amount of the uncrosslinked b-DCPU liquid precursor created a uniform interfacial layer on the PCL-containing component. Precise placement then brought the coated component into close contact with a preformed b-DCPU film. Heating at 55 °C until complete curing crosslinked the precursor in situ and formed a stable rigid–soft interface between the PCL-containing component and the b-DCPU film.

#### Bonding between b-DCPU components

The b-DCPU liquid precursor served as an interfacial adhesive between two cured b-DCPU films. A thin layer of the uncrosslinked precursor covered the mating surface of one film, and precise alignment brought the second film into close contact with the coated surface. Heating at 55 °C until complete curing promoted in situ polymerization and enabled the precursor to form interfacial crosslinks with the existing dynamic covalent polyurethane networks on both sides. This process created a continuous and seamless bond between the two b-DCPU films.

### Scaling analysis of pressure-induced membrane displacement

#### Design and fabrication of test structures for scaling analysis

Spin coating PDMS (Sylgard 184, Dow Corning) on acrylic substrates produced the membranes with different thicknesses. Moulding produced PDMS support rings with circular apertures of defined radii. Corona-discharge treatment activated the membrane and support-ring surfaces before membrane transfer and bonding. The resulting test structures consisted of a PDMS membrane spanning the aperture of a PDMS support ring. Separate experimental series varied the aperture size, initial membrane thickness and membrane modulus. Water-soluble wax sealed each test structure to a pressure chamber while leaving the pressure inlet unobstructed. A pressure controller regulated the applied pressure. A digital microscope (VHX-7000, Keyence) recorded membrane deformation. Analysis of the microscope images determined the central membrane displacement (*x*) relative to the undeformed configuration.

#### Scaling analysis of pressure-induced membrane displacement

The mean of three measurements defined the initial membrane thickness (*t*_0_) for each membrane condition. Tensile measurements determined the membrane modulus according to *E* = *kL*_0_/*wt*_0_, where *k* was the force–extension slope, *L*_0_ was the initial gauge length, *w* was the specimen width and *t*_0_ was the initial thickness. Producing the different modulus conditions required the preparation of separate membrane samples. Supplementary Note 2 describes the empirical scaling analysis that relates the measured displacement to the applied pressure, aperture radius, initial membrane thickness and membrane modulus.

### Design of the cage for wireless power transmission

A double loop litz coil was wound around a rat cage with a length of 28 cm and a width of 15 cm. It was tuned to operate around 13.56 MHz by capacitors. A customized signal generator and amplifier drove the coil to generate alternating magnetic field. An auto antenna tuner was connected between the coil and signal generator to achieve optimized power efficiency by impedance matching.

### Design of the wearable MF camera

#### Overall architecture

The wearable circuit consisted of two parts. One was a Hall sensor array with 32 digital Hall sensors (BMM150, Bosch Sensortec), which read the magnetic flux density from the bioresorbable magnets on the implanted device and mapped the distribution of the magnetic field. The other was a Bluetooth microcontroller that got readings from the Hall sensor array and wirelessly transmitted the data to user interfaces through Bluetooth universal asynchronous receiver–transmitter (UART) communication.

#### Hall sensor array

The Hall sensor array had 32 digital Hall sensors, which were low in noise (lowest noise of ∼0.3 μT) and compact in size (1.5 mm in width and length, 0.6 mm in height). We arranged 32 sensors in a 14 mm × 14 mm square, with the distance between two sensors being 2.5 mm, which was realized by utilizing a 4-layer PCB and via-in-pad design to minimize the room for the tracks on the circuit board. The Hall sensors communicated with the microcontroller through inter-integrated circuit (I^2^C) interface. Because there were only two I^2^C address pins on each Hall sensor, corresponding to only four I^2^C addresses, we utilized an 8-channel I^2^C multiplexer (TCA9548A, Texas Instruments) to enable the multiplexing of I^2^C interface. Every 4 sensors with different I^2^C addresses (0×10, 0×11, 0×12, and 0×13) were connected in one I^2^C bus from one channel on the multiplexer, which helped communicating with 32 sensors with only one I^2^C bus from the microcontroller.

#### Bluetooth microcontroller circuit

The Bluetooth microcontroller circuit used a BLE system-on-chip (SoC, nRF52832, Nordic Semiconductor) to control data acquisition and wireless communication. Through the I²C protocol, the BLE SoC configured the registers of the Hall sensors and read magnetic-field measurements from both the sensor array and the reference sensor. The microcontroller then transmitted the data wirelessly to the user interface through a UART-over-Bluetooth Low Energy link. The circuit board also incorporated an outward-extending arm with a Hall sensor at its distal end to provide an environmental magnetic-field reference. This reference sensor matched the array sensors in both model and mounting orientation, allowing it to measure ambient magnetic-field variations synchronously. The BioBeacon dimensions and the required sensor-placement geometry determined the arm length.

#### Connection between Hall sensor array and Bluetooth microcontroller

Two interconnection schemes connected the Hall sensor array board to the Bluetooth microcontroller board. The first scheme used four flexible wires, each 1.5 cm long, to connect the two boards. During normal operation, folding the boards about the flexible interconnect created a compact stacked configuration and reduced the lateral footprint of the system to 2.9 cm × 1.9 cm. The second scheme used a 4-pin, 1.25-mm-pitch pin header and a mating socket header. This plug-and-play design enabled rapid connection and disconnection, facilitating board installation, removal, and replacement.

#### Wireless power receiver and battery-charging circuit

A compact receiver coil integrated with the MF camera captured the alternating magnetic field generated by the cage-mounted transmitter coil. The antenna terminals fed a bridge rectifier (BAS4002ARPP, Infineon Technologies), which converted the induced alternating voltage into a direct voltage. A 10-μF capacitor smoothed the rectified output, while a Zener diode provided overvoltage protection. A step-down converter (TPS82130, Texas Instruments) regulated the output to 5 V and supplied the onboard single-cell lithium-battery charging circuit. The rechargeable battery subsequently powered the MF-camera electronics, enabling continuous operation during wireless energy transfer.

#### Data-acquisition mode and sampling-rate configuration

The MF camera firmware operated all 32 Hall sensors and one environmental reference Hall sensor in forced mode. At the beginning of each data frame, the BLE SoC sequentially issued a forced-measurement command to all sensors. After the reference sensor asserted its data-ready signal, the microcontroller read the tri-axial magnetic-field data from all sensors, compensated for the environmental magnetic field, and transmitted the complete frame through BLE. Each Hall sensor automatically returned to sleep mode after completing one measurement and remained idle until the next trigger. The firmware independently adjusted the internal measurement repetition counts for the X/Y and Z axes, with maximum implemented settings of 200 and 250 repetitions, respectively. Increasing the repetition counts enhanced internal averaging, reduced output noise, and improved measurement precision, but also increased the conversion time and energy consumption. The frame-trigger interval independently controlled the system sampling rate according to:

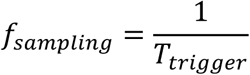

A trigger interval of 33.3 ms corresponded to a sampling rate of 30 Hz. To complete every measurement before the next trigger, the firmware used lower repetition counts at higher frame rates and higher repetition counts with longer trigger intervals for high-precision measurements. This configuration allowed the MF camera to balance sampling rate, measurement noise, and power consumption according to different application requirements. In addition, the user interface transmitted control commands through the BLE link to switch remotely among preset sampling modes during operation. Upon receiving a command, the microcontroller updated the frame-trigger period and the corresponding X/Y- and Z-axis measurement repetition counts, thereby dynamically balancing sampling rate, measurement noise, and power consumption according to application requirements.

#### Power analysis

A source meter (Keithley 2450) characterized the power consumption of the MF camera. The source meter operated in source-voltage/measure-current mode, supplied a constant voltage of 3.7 V to the MF camera, and simultaneously recorded the operating current. The test program varied the internal X/Y- and Z-axis measurement repetition counts of the Hall sensors to evaluate power consumption under different measurement-precision settings. The instantaneous power followed

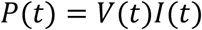

where *P*(*t*) denotes the instantaneous power at time t, *V*(*t*) denotes the supply voltage and *I*(*t*) denotes the operating current. Numerical integration of the power–time curve for one complete magnetic-field frame in Origin 2026 (OriginLab) yielded the per-frame acquisition and transmission energy according to

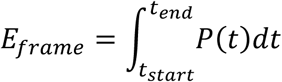

Each complete frame contained tri-axial magnetic-field measurements from 32 array sensors and one environmental reference sensor. A power monitoring chip (INA226, Texas Instruments) recorded the voltage and current profiles during long-term operation. Every 33 s, the data-logging program collected a burst of four consecutive measurements at 1 s intervals. Throughout the wireless-power test, the wireless charging system continuously supplied the MF camera while the camera performed high-precision magnetic-field acquisition and wireless data transmission. All Hall sensors used 160 X/Y-axis repetitions and 200 Z-axis repetitions. The battery-powered control followed the same acquisition protocol without wireless charging. The resulting voltage–time profiles enabled comparison of the long-term operating stability under battery-powered and wirelessly powered conditions.

### Collection of datasets for deep learning

In order to collect magnetic field readings from the Hall sensor array under different pressures, we used a pressure controller (PC1, FluidicLab). A CO₂ laser (VLS3.60DT Platform-SYS, Universal Laser Systems Inc.) cut a 4-mm hole on a circular-shaped poly(methyl methacrylate) (PMMA) plate, followed by sticking the plate on the back of a syringe and sealing the edge with a water-soluble wax (Crystalbond 555, TED PELLA Inc.). Adhering and sealing the implantable device on the hole with the wax, followed by connecting the syringe head to the pressure controller with a silicone tube completed the structure for simulating the change of pressure.

In order to collect magnetic field readings from the Hall sensor array in different relative positions and orientations, we used a 6-degree-of-freedom robotic arm (UR3e, Universal Robots). The MF camera was mounted on the robotic arm, while the BioBeacon was fixed on an optical table. The robotic arm moved the MF camera through predefined positions and orientations relative to the stationary BioBeacon. The robotic arm provided three translational and three rotational degrees of freedom across the required workspace, with a nominal pose repeatability of ±30 μm. The pressure and pose sampling schemes, acquisition protocol, and training, validation and test dataset configurations are detailed in Supplementary Note 8.

### Machine-learning models for pressure and pose prediction

Each sample used the magnetic-field readings from 32 tri-axial Hall sensors as a 96-feature input vector and the simultaneously recorded pressure as a scalar regression target. Data preprocessing excluded the three channels from the environmental reference sensor. StandardScaler from the scikit-learn Python library calculated the mean and standard deviation of each feature from the training set and applied the same transformation to the validation data.

Bottom row of Fig. 3j,k used a fully connected multilayer perceptron (MLP) for pressure prediction. The network contained four hidden layers with 128, 256, 256, and 128 neurons, respectively. A rectified linear unit (ReLU) followed each hidden layer, and a final linear layer generated one pressure estimate. Training used the mean squared error (MSE) loss, the Adam optimizer, and a learning rate of 1 × 10^-4^. The training program compared batch sizes of 512, 1024, 1536, and 2048 and evaluated model checkpoints throughout a maximum of 1000 epochs. The results in bottom row of Fig. 3j,k used the checkpoint at epoch 700 with a batch size of 2048. Python’s pickle module serialized the selected complete MLP model to a local file for subsequent pressure prediction. All other pressure-prediction experiments used a residual multilayer perceptron (ResMLP). An input layer first projected the 96-dimensional feature vector into a 256-dimensional hidden space, and three residual blocks then extracted higher-level features. Each residual block contained a 256-dimensional fully connected layer, a ReLU activation, a dropout layer with a rate of 0.1, and a second 256-dimensional fully connected layer. A shortcut connection added the block input directly to its output. A final linear layer mapped the 256-dimensional representation to one pressure estimate. We compared the same set of batch sizes for ResMLP as for the MLP. ResMLP was trained to minimize the MSE loss using the AdamW optimizer, with an initial learning rate of 2 × 10^-5^ and a weight-decay coefficient of 1 × 10^-4^. A cosine-annealing warm-restart scheduler restarted the learning-rate cycle every 100 epochs and used a minimum learning rate of 1 × 10^-7^. Mixed-precision computation accelerated training for a maximum of 8000 epochs. Early stopping terminated training when the validation MSE showed no improvement for 600 consecutive epochs. PyTorch stored the model parameters that achieved the lowest validation MSE. MAE, MSE, and *R*^2^ jointly quantified the pressure-prediction performance of both networks. Five-dimensional pose prediction used a separately trained ResMLP with the same hidden architecture and a five-output linear layer, with the checkpoint selected according to the lowest validation MAE on standardized pose targets.

#### Data transmission, processing, and real-time pressure decoding on the computer

#### Bluetooth communication

The built-in Bluetooth adapter of the computer established a direct BLE connection with the BLE SoC (nRF52832) in the wearable circuit without an external Bluetooth receiver. A Python program used the Bleak library for BLE communication to discover and connect to the target device and then identified the Nordic UART Service characteristics through their UUIDs. The program subscribed to the TX characteristic to receive sensor data in real time and used the RX characteristic to transmit control commands to the BLE SoC. An automatic reconnection routine continuously attempted to restore communication after an unexpected disconnection.

#### Data packet processing

The nRF52832 packed each data frame into a 198-byte payload and transmitted it to the computer via BLE. A byte buffer reconstructed the incoming Bluetooth data stream into 198-byte packets. The program decoded each packet as 99 little-endian, 16-bit signed integers. The first 96 values contained the tri-axial magnetic-field readings from the 32 Hall sensors, and the final three values contained the tri-axial readings from the environmental reference sensor. The nRF52832 firmware compensated the array signals for the environmental MF before transmission; therefore, the computer directly used the first 96 channels as pressure-decoding features.

#### Real-time pressure decoding

After device-specific calibration, the computer program loaded the corresponding feature scaler and pretrained pressure-regression model. For each frame, the program standardized the 96 magnetic-field features using the same parameters as the training process and fed the resulting feature vector into the corresponding MLP or ResMLP model. The model then produced a calibrated pressure estimate in kPa in real time. The current program implements the ResMLP model and can load another model file that matches a specific device or experiment.

#### User interface and data logging

A graphical interface developed with PyQt5 GUI toolkit and the PyQtGraph plotting library displayed the decoded pressure numerically, continuously updated the pressure–time trace, and visualized the tri-axial magnetic-field maps and field-magnitude map. The interface used a 10-ms software refresh interval. The program also recorded the timestamps, 96 array channels, three reference-sensor channels, and model outputs in a CSV file. Through the same BLE connection, the computer transmitted control commands to the nRF52832 and remotely switched among different sampling modes.

### Animal studies

#### Animals and ethical approval

All procedures associated with animal studies followed the ethical guidelines of the Institutional Animal Center Peking University. The Institutional Animal Care and Use Committee (IACUC) at Peking University (Protocol No. FT-HanMD-1) approved the protocol. Male rats (SD and Wistar, age at initiation of the treatment: at least 9 weeks, but no more than 15 weeks, purchased from Beijing Vital River Laboratory Animal Technology Co., Ltd.) were habituated for up to 3 days before surgery. Vaporized isoflurane induced and maintained anaesthesia throughout all surgical procedures. The postoperative regimen included penicillin sodium for infection prophylaxis and meloxicam for analgesia.

#### Implantation of BioBeacon and ICP validation

We removed the fur from the dorsal surface of the head, disinfected the surgical field, and secured the rat in a stereotaxic frame. A midline scalp incision exposed the skull, and a skull drill produced a circular cranial opening ∼5.5 mm in diameter over the parietal region without breaching the dura. Sterile saline filled the cranial opening and established a fluid environment for pressure transmission and BioBeacon testing. A biocompatible adhesive (Dyad Flow, Kerr) secured the BioBeacon to the margin of the cranial opening and sealed the interface between the BioBeacon and the skull. A thin thermoplastic polyurethane (TPU) tube connected the sealed space between the BioBeacon and the cranial opening to a commercial pressure transducer. The tube transmitted fluid pressure from this sealed space to the commercial transducer, allowing the BioBeacon and the commercial transducer to monitor pressure changes within the same local environment. Placement of the MF camera above the BioBeacon approached its sensing area near the implant. The MF camera recorded the dynamic magnetic signal from the BioBeacon at a sampling rate of 15 Hz, and a PowerLab data-acquisition system (16/35, ADInstruments) simultaneously acquired the reference pressure signal from the commercial transducer at a sampling rate of 1 kHz. A 2.25-mmHg offset was added to the measurements from commercial transducer to account for the height-dependent hydrostatic offset.

#### Spectral analysis

For spectral comparison, the predicted pressure signal and the measurements from the commercial pressure transducer were independently mean-centred and multiplied by a Hann window (w(n)). The fast Fourier transform (FFT) of each windowed signal yielded X(f), from which the one-sided amplitude spectrum for positive frequencies below the Nyquist frequency was calculated as *A*(*f*) = 2|*X*(*f*)|/ ∑*_ss_ w*(*n*). Spectral amplitudes were reported in mmHg. Following the comparison, inspection of the implant and surgical site confirmed the integrity of the implantation. The subsequent experimental design determined whether the reference tubing remained in place or left the cranial opening. Surgical-site cleaning and scalp suturing completed the procedure.

#### Freely moving measurements and tail-lifting tests

For measurements in awake and freely moving rats, nonmagnetic bone screws provided stable anchoring points in the skull around the BioBeacon implantation site. A custom aluminium-alloy support frame sat adjacent to the bone screws, and biocompatible adhesive bonded the frame to the screws. Additional screws secured the MF camera to the support frame, confirming stable attachment and continuous BioBeacon signal acquisition during animal movement. After each rat had fully recovered from anaesthesia, we placed it in the cage and continuously recorded the BioBeacon signal while the animal walked, turned, and changed posture freely. During the tail-lifting test, an operator briefly lifted the rat by its tail, thereby inducing a pronounced whole-body postural change while the head-mounted MF camera continuously monitored the BioBeacon at a sampling rate of 10 Hz. The normalized motion intensity was calculated from the tri-axial measurements of the reference sensor. At each time point (t), the initial three-axis field vector *B*(*t*_0_) was subtracted from the field vector (*B*(*t*)), and the Euclidean norm of the resulting vector difference was calculated as *M*(*tt*) = ‖*B*(*t*) − *B*(*t*_0_)‖. The resulting time series was normalized for visualization. The same point-by-point calculation was applied independently to the tri-axial measurements of each MF-camera sensor to obtain the displayed channel-wise Δ*B*. These baseline-referenced quantities were used for visualization in the plots, whereas the machine-learning model used the original geomagnetic-compensated multichannel MF measurements as inputs.

#### Transient middle cerebral artery occlusion and reperfusion models

The intraluminal filament method^44^ established a transient middle cerebral artery occlusion (t-MCAO) model. We first implanted the BioBeacon and installed the nonmagnetic bone screws, aluminium-alloy support frame, and MF camera as described above. An ∼2-cm midline cervical incision exposed the common, external and internal carotid arteries (CCA, ECA and ICA). Microsurgical dissection separated these arteries from the surrounding fascia and connective tissues while preserving the adjacent nerves. Electrocautery divided the occipital and superior thyroid arteries, a permanent ligature closed the distal ECA, and a loose slipknot encircled its proximal segment. With the CCA and ICA temporarily clamped, a filament (A5-204005, Cinontech) entered the ECA through a small arteriotomy. After release of the ICA clip, the filament advanced through the ICA until its marker reached the predefined reference position and its tip occluded the origin of the middle cerebral artery. Tightening of the slipknot secured the filament, and release of the CCA clip restored carotid flow. After 1 h of occlusion, the CCA was temporarily clamped and the filament partially withdrawn. The ICA was then clamped before complete filament removal. Permanent ligation of the proximal ECA prevented bleeding, and sequential release of the ICA and CCA clips initiated reperfusion. Layered sutures closed the cervical incision after haemostasis was confirmed.

#### Assessment of successful t-MCAO induction

After each rat fully recovered from anaesthesia, we assessed neurological function at predefined time points. We first lifted the rat gently by the tail, allowed all limbs to leave the supporting surface, and examined bilateral forelimb extension. Healthy rats normally extended both forelimbs symmetrically, whereas rats with successful t-MCAO commonly showed flexion, adduction, or incomplete extension of the forelimb contralateral to the occluded hemisphere. We then placed the rat on a flat open surface and observed spontaneous locomotion, body inclination, and circling behaviour. Persistent circling or falling toward the paretic side contralateral to the occluded hemisphere indicated a lateralized neurological deficit. After the final signal-acquisition session, 2,3,5-triphenyltetrazolium chloride (TTC) staining provided histological confirmation of cerebral infarction. Immediate removal of the whole brain after euthanasia and omission of fixatives before staining preserved tissue enzymatic activity. We chilled the fresh brain at −20 °C for 15 min to slightly firm the tissue and then prepared consecutive coronal sections using a blade. Incubation in freshly prepared 2% TTC solution in phosphate-buffered saline proceeded at 37 °C for 20 min in the dark, while gentle inversion every 5 min promoted uniform staining. 4% paraformaldehyde preserved the sections for subsequent storage. Dehydrogenases in viable brain tissue reduced TTC to red formazan and produced a brick-red appearance, whereas infarcted tissue lacked sufficient enzymatic activity and remained pale or white.

#### Data acquisition and motion quantification in t-MCAO and sham-operated rats

During filament insertion, the occlusion period, and filament withdrawal, the MF camera continuously acquired data from the t-MCAO rat at 4 Hz, with the same sampling rate used for the sham-operated rat. To reduce battery consumption during the 120-h monitoring period, the MF camera recorded five consecutive magnetic-field measurements per acquisition group. Samples were acquired at 0.5-s intervals, with group starts 121.5 s apart, in both rats. For motion analysis, the tri-axial magnetic-field readings within each acquisition group were averaged to obtain a vector *B_k_*, using the reference sensor. Motion-related magnetic variation between consecutive groups (relative motion intensity) was calculated as (100‖*B_k_*− *B_k_*_−1_‖/[(‖*B_k_*‖ + ‖*B_k_*_−1_‖)/ 2]). Changes across gaps exceeding 300 s were excluded from the summary statistics. Medians and interquartile ranges calculated in 1.5-h bins characterized the temporal profiles. For the day–night comparison, valid changes were assigned according to the timestamp of the current group, and their arithmetic means were calculated separately for 08:00–20:00 and 20:00–08:00.

#### In vivo degradation test of FeNWs

A 16 wt% solution of Pluronic F127 in physiological saline served as the dispersion medium. Thorough mixing dispersed 5 mg of FeNWs uniformly in 1 mL of the F127 solution and formed a homogeneous FeNW/F127 suspension. Storage at 4 °C maintained the suspension in a liquid and injectable state. Equal division of the 1-mL suspension among three rats delivered ∼333 μL of suspension and 1.67 mg of FeNWs to the subcutaneous cervical region of each rat. The experimental protocol euthanized two of the corresponding rats on day 1 and day 279 after injection and collected tissue from the injection sites to evaluate the long-term *in vivo* degradation of FeNWs.

## Supporting information

Supplementary Video 1-Magnetic-field-induced alignment of iron nanowires

Supplementary Video 2-In vitro demonstration of constant pressure sensing under multi-axis motion

Supplementary Video 3-In vitro demonstration of sinusoidal pressure sensing under multi-axis motion

Supplementary Video 4-Wireless intracranial pressure monitoring in a freely moving rat

Supplementary Video 5-Behavioural deficits following transient middle cerebral artery

Supplementary Video 6-Different respiratory states in a rat and corresponding ICP waveforms

## Data availability

All data supporting the findings of this study are included in the paper and its Supplementary Information. Source data are provided with this paper.

## Code availability

The code used in this study, including the firmware for MF-camera data acquisition and transmission, the Python programs for BLE data reception and real-time pressure decoding, and the scripts for data preprocessing and ResMLP model training, is publicly available at GitHub (https://github.com/niezhongyinzy/code_for_BioBeacon).

## Acknowledgements

We acknowledge the financial support from the Scientific Research Innovation Capability Support Project for Young Faculty (ZYGXQNJSKYCXNLZCXM-H1), the Beijing Natural Science Foundation (JQ26045), the National Natural Science Foundation of China (62471007), and the Basic Research Program of Jiangsu (BM2024001).

## Author contributions

These authors contributed equally: Zhongyi Nie, Xiang Lin, Ji Wan, Jie Xu Z.N. and Mengdi Han conceived the idea and designed the research. Z.N., D.W., Minki Hong, Yaozheng Wang and Yiran Wang synthesized the materials and fabricated the BioBeacon implants. Z.N., K.W., S.Z., Y.H. and C.Z. developed the MF-camera hardware and data-acquisition software. Z.N. and J.W. conducted theoretical analyses and numerical simulations. Z.N., K.W. and J.W. developed the decoding algorithms and analysed the data. Z.N., X.L., R.Z. and C.Y. prepared the figures. Z.N. and X.L. performed in vitro characterization and biocompatibility assessments. Z.N., X.L., J.X., W.M. and Yan Wang conducted the animal experiments. Z.N., X.L., N.S. and Mengdi Han wrote and revised the manuscript. Y.S., H.H., J.K. and Mengdi Han supervised the research.

