## Supplementary information for "Bioresorbable permanent-magnet beacons for tracking internal biomechanics during free movement"

### Supplementary Notes

#### Supplementary Note 1. Tissue-transparent magnetic fields facilitate motion-tolerant sensing

*1.1 Unified measurement model*

For a transcutaneous sensing system, the measured signal generally depends on the target physiological variable, the relative pose between the implant and the external reader, and the properties of the intervening medium. A unified measurement model can be written as

$$\mathbf{y}_k = \mathbf{f}_k(x, \mathbf{q}, \boldsymbol{\eta}_k). \quad \text{(S1)}$$

Here, $x$ is the target physiological variable, $\mathbf{q}$ describes the relative position and orientation between the implant and the external reader, and $\boldsymbol{\eta}_k$ represents the intrinsic properties of the intervening tissues and media relevant to readout modality $k$. The relative pose $\mathbf{q}$ also determines the propagation path, incidence direction, and geometric alignment between the implant and the reader.

*1.2 Pose dependent transmission and coupling in alternative readout modalities*

For optical, acoustic, and resonant or inductive readouts, the measurement can be approximately expressed as

$$\mathbf{y}_{\text{alt}} = A_{\text{alt}}(\mathbf{q}, \boldsymbol{\eta}_{\text{alt}})\, \mathbf{s}_{\text{alt}}(x). \quad \text{(S2)}$$

Here, $\mathbf{s}_{\text{alt}}(x)$ represents the signal generated by the physiological variable, and $A_{\text{alt}}$ represents the effective transmission or coupling between the implant and the external reader. Changes in relative position or orientation can alter the propagation path, incidence angle, alignment, or coupling efficiency. For example, motion can change the optical or acoustic path through heterogeneous tissues, as well as the mutual inductance and alignment of resonant coils. Consequently, motion-induced changes in $A_{\text{alt}}$ may become mixed with changes caused by $x$. Without additional information about pose, propagation, or coupling, these two contributions can be difficult to distinguish.

*1.3 Tissue transparent quasi static magnetic readout*

Biological tissues and fluids have relative magnetic permeabilities close to 1 and produce negligible attenuation or distortion of quasi-static magnetic fields under the operating conditions considered here. Therefore, the dependence of the magnetic forward model on the properties of the intervening tissues can be neglected:

$$\mathbf{f}_{\mathrm{MF}}(x, \mathbf{q}, \boldsymbol{\eta}_{\mathrm{MF}}) \approx \mathbf{h}(x, \mathbf{q}). \tag{S3}$$

The corresponding measurement can thus be written as

$$\mathbf{y}_{\mathrm{MF}} \approx \mathbf{h}(x, \mathbf{q}) + \mathbf{B}_{\mathrm{geo}}, \tag{S4}$$

where $\mathbf{h}$ is the MF generated by the implant and $\mathbf{B}_{\mathrm{geo}}$ denotes the geomagnetic field. Unlike a tissue-dependent transmission or coupling factor, this background does not modulate the magnetic signal during its passage through the body and can be compensated separately.

The negligible tissue-induced attenuation of quasi-static magnetic fields avoids motion-dependent transmission variations associated with changes in the intervening tissue path, thereby providing a physical basis for motion-tolerant transcutaneous sensing. The measured magnetic field (MF) distribution nevertheless remains sensitive to changes in the relative position and orientation between BioBeacon and the MF camera. These pose-induced variations are separated from the target biomechanical signals through multi-magnet encoding and data-driven reconstruction, as described in Supplementary Notes 6 and 7.

#### Supplementary Note 2. Empirical scaling of pressure-induced membrane displacement

To establish a quantitative basis for tailoring pressure-sensing BioBeacon devices to different pressure ranges and anatomical constraints, the dependence of the pressure-induced central membrane displacement $x$ on the applied pressure $P$, aperture radius $r$, initial membrane thickness $t_0$, and membrane modulus $E$ is examined. Because membrane deformation increases the membrane surface area and reduces its effective thickness, a geometric correction is first introduced before evaluating the scaling relationship.

The deformed membrane is approximated as a spherical cap with a base radius $r$, corresponding to the aperture radius, and a height $x$. Its initial projected area, $A_0$, and deformed surface area, $A$, are

$$A_0 = \pi r^2, \qquad A = \pi(r^2 + x^2). \tag{S5}$$

Assuming an approximately conserved membrane volume and a uniform effective thickness gives the corrected thickness

$$t_{\mathrm{corr}} = t_0 \frac{A_0}{A} = \frac{t_0 r^2}{r^2 + x^2}. \qquad \text{(S6)}$$

Here, $t_{\mathrm{corr}}$ represents an area-averaged geometric estimate of the membrane thickness after deformation rather than a direct measurement of the local thickness.

For a reduced model containing $x$, $P$, $E$, $r$ and $t_{\mathrm{corr}}$, dimensional consistency requires $x/r$ to depend only on $P/E$ and $t_{\mathrm{corr}}/r$. A separable power-law approximation of this dependence can be written as

$$\frac{x}{r} \approx C \left(\frac{P}{E}\right)^{\alpha} \left(\frac{r}{t_{\mathrm{corr}}}\right)^{\beta}, \qquad \text{(S7)}$$

where $C$ is dimensionless and $\alpha$ and $\beta$ are empirical exponents. Rearranging equation (S7) gives the corresponding dependence of $x$ on the four parameters:

$$x \approx C \frac{P^{\alpha} r^{1+\beta}}{E^{\alpha} t_{\mathrm{corr}}^{\beta}}. \qquad \text{(S8)}$$

Four representative single-variable sweeps are analysed to evaluate the individual parameter dependencies (fixed conditions are given in Supplementary Fig. 13a–d). The displacement shows a linear relationship with $r^{1.75}$ for nominal aperture radii of 1.5–3.0 mm ($R^2$ = 0.9974; Supplementary Fig. 13a), with $t_{\mathrm{corr}}^{-0.75}$ for initial thicknesses of 33.71–77.82 μm ($R^2$ = 0.9707; Supplementary Fig. 13b), with $P^{0.75}$ for pressures increasing from 0 to 8 kPa in 0.5-kPa increments ($R^2$ = 0.9967; Supplementary Fig. 13c), and with $E^{-0.75}$ for moduli of 0.547–1.754 MPa ($R^2$ = 0.9897; Supplementary Fig. 13d). Varying the modulus requires the preparation of separate membrane samples. Despite this independent fabrication, their initial thicknesses remain closely matched (46.46–48.74 μm; inter-sample range <5% of the mean), whereas $E$ varies by approximately 3.2-fold. The modulus is therefore treated as the primary swept variable, and the small remaining differences in $t_0$ are accounted for in the pooled analysis by using the specimen-specific $t_0$ values to calculate $t_{\mathrm{corr}}$.

These single-variable sweeps support an empirical description with $\alpha = \beta = 0.75$ over the investigated parameter ranges. Equation (S8) therefore becomes

$$x \approx CS, \qquad S = \frac{P^{0.75} r^{1.75}}{E^{0.75} t_{\mathrm{corr}}^{0.75}}, \tag{S9}$$

where $S$ is the scaling coordinate. For calculating $S$, $P$ and $E$ are expressed in kPa, whereas $r$ and $t_{\mathrm{corr}}$ are expressed in μm, giving $S$ in μm.

The combined scaling analysis includes 95 measurements from 15 experimental series spanning $P = 0$–$8$ kPa, $r = 1.50$–$3.00$ mm, $t_0 = 33.71$–$77.82$ μm and $E = 0.547$–$1.754$ MPa. The pooled measurements approximately collapse onto a common linear trend, $x = 0.742S + 159.3$ μm ($n = 95$, $R^2 = 0.8681$; Supplementary Fig. 13e).

#### Supplementary Note 3. Physical origin of the enhanced coercivity and remanence of Fe nanowires

Bulk Fe generally contains multiple magnetic domains separated by domain walls. Under an external magnetic field, the movement of these domain walls provides a relatively low-energy pathway for magnetization reversal, resulting in low coercivity and remanence. When Fe is confined into nanowires with diameters below the critical size for stable domain-wall formation, the reduction in magnetostatic energy can no longer compensate for the energy required to form a domain wall. The nanowires therefore approach a single-domain or quasi-single-domain state, in which domain-wall-mediated reversal is suppressed. For ideal Fe nanowires with an aspect ratio of ~10, the theoretical single-domain critical diameter is about 140 nm. The 30- and 60-nm-diameter FeNWs investigated here are both below this characteristic size. Their elongated geometry also produces strong shape anisotropy, with a smaller demagnetizing factor along the nanowire axis than perpendicular to it. The nanowire axis consequently becomes the magnetic easy axis, stabilizing the axial magnetization and contributing to the enhanced remanence.

Within the single-domain size range, magnetization reversal is primarily governed by two idealized mechanisms: curling and coherent rotation. When the diameter lies between the coherent-rotation threshold and the single-domain critical size, the nanowire diameter remains sufficiently large to allow the magnetization to vary spatially without a prohibitive exchange-energy cost. During reversal, the magnetic moments rotate nonuniformly around the nanowire axis, forming a vortex-like curling configuration across the cross-section. This configuration reduces the transverse magnetization and hence the magnetostatic energy, but increases the exchange energy because neighbouring magnetic moments are no longer parallel. In the idealized model, when the diameter decreases below the coherent-rotation threshold, the magnetization must vary over a shorter distance during curling, rapidly increasing its exchange-energy cost. Curling therefore becomes energetically unfavourable, and the magnetic moments remain approximately parallel and rotate collectively through coherent rotation. The Stoner-Wohlfarth model describes this ideal coherent-rotation limit.

These two mechanisms represent idealized limits of magnetization reversal. In real nanowires, finite length produces enhanced and nonuniform demagnetizing fields near the wire ends, allowing magnetization reversal to nucleate locally in these regions. Microstructural inhomogeneities and misalignment between the applied field and the nanowire axis further promote deviations from the ideal reversal behaviour. These effects can produce mixed reversal pathways involving both curling-like and quasi-coherent rotation. Nevertheless, decreasing the nanowire diameter increases the exchange-energy penalty for spatially nonuniform magnetization, thereby progressively suppressing curling-dominated reversal and increasing the coercivity toward the coherent-rotation limit. The enhanced stability of the axial magnetization can consequently contribute to higher remanence and hysteresis-loop squareness.

Consistent with this size-dependent behaviour, the 30-nm FeNWs exhibit a coercivity of ~750 Oe and a remanent magnetization of ~12 emu/g, compared with ~500 Oe and ~7 emu/g, respectively, for the 60-nm FeNWs (Supplementary Fig. 22). The higher coercivity of the 30-nm FeNWs is consistent with the stronger suppression of nonuniform magnetization reversal at reduced diameters, while their higher remanence is also consistent with the enhanced stability of the axial magnetization.

**Supplementary Note 4. Comparison with macroscopic magnetic structures made from bioresorbable magnetic materials**

For an implanted magnetic beacon, the remanence ($M_r$) of the macroscopic magnet determines the field generated without external excitation, whereas its coercivity ($H_c$)

determines the resistance of this magnetization to reversal. Bioresorbable magnetic materials, including iron, iron oxides, iron-substituted hydroxyapatite and molecular magnets, have been processed into macroscopic structures, mostly by dispersion in polymer or inorganic matrices. Their coercivities range from negligible to moderate (up to ~250 Oe in polymer-based structures), but their remanence remains below ~10 emu/g in all cases. An $\varepsilon$-$Fe_2O_3$/silica monolith reaches an $H_c$ above 18000 Oe but an $M_r$ of only ~0.8 emu/g.

By combining the shape anisotropy of single-domain FeNWs with field-assisted alignment of their easy axes (Supplementary Note 3 and Fig. 2f,g), the FeNW/PCL nanocomposite achieves an $M_r$ of ~30 emu/g, more than three times the highest reported value and nearly 40 times that of the $\varepsilon$-$Fe_2O_3$/silica monolith, together with an $H_c$ of ~400 Oe. This coercivity exceeds the geomagnetic field (~0.5 Oe) by nearly three orders of magnitude, sufficient to keep the magnetization stable against ambient fields. Even the unaligned composite ($M_r$: ~15 emu/g) exceeds these structures in remanence. Values for representative structures[1–10], normalized to their total mass, are summarized in Supplementary Table 2. When $M_r$ or $H_c$ was not reported directly, it was read from the published hysteresis loops or estimated from the other parameter and the reported magnetic susceptibility.

**Supplementary Note 5. Geomagnetic compensation with the reference sensor**

To compensate for geomagnetic variations during untethered operation, the reference sensor is positioned at an adjustable distance from the main array, allowing it to track geomagnetic variations with negligible contribution from BioBeacon (Fig. 3b). Distance-dependent measurements show that the MF generated by BioBeacon decreases to less than 2 μT (comparable to the noise of the Hall sensors) at a lateral separation of 20 mm and a vertical separation of 8 mm (Supplementary Fig. 27). The reference sensor is therefore positioned laterally 23 mm and vertically 9 mm from the Hall sensor array. Reference subtraction removes the common-mode variations observed in the raw and reference signals (Fig. 3c, top and middle), resulting in a compensated waveform that reveals the simulated physiological signal (Fig. 3c, bottom).

#### Supplementary Note 6. Physical basis of motion-tolerant pressure sensing based on dual-magnet design

*6.1 Fixed-magnet referencing for pressure–motion separation*

Magnetic pressure sensing requires distinguishing membrane deformation from rigid-body motion between the implant and the MF camera. In a single-magnet configuration, both processes change the distance and orientation between the magnet and the MF camera. Consequently, pressure and rigid-body translation can produce similar magnetic responses (Supplementary Fig. 28a).

Introducing a second, mechanically fixed magnetic element provides an internal spatial reference. In the dual-magnet configuration, pressure displaces only the membrane-mounted magnet relative to the fixed magnet, whereas rigid-body motion collectively translates and rotates both magnets without changing their internal relative configuration. These two processes therefore produce different spatial superpositions of the magnetic fields generated by the two elements (Supplementary Fig. 28b).

Supplementary Fig. 28 uses two cylindrical magnets as a simplified conceptual model and shows only axial rigid-body translation. In the actual BioBeacon-P/PA, the fixed reference magnet is annular (Supplementary Fig. 8); this difference in geometry does not change the principle. The formulation below extends it to scenarios of three-dimensional translations and rotations.

*6.2 Pose-dependent forward model of the dual-magnet system*

Here, the target variable in equation (S1) is pressure, $x = P$. Let A and B denote the fixed and movable magnets, respectively. Pressure produces a membrane displacement $\delta(P)$ that translates B along the implant axis $\mathbf{e}_z$, while A remains fixed within the implant. Let $\mathbf{p}_A$ and $\mathbf{p}_B$ denote the nominal magnet positions within the implant. The implant pose $\mathbf{q}$ relative to the MF camera is described by a rotation $\mathbf{R}$ and a translation $\mathbf{t}$. The magnet positions in the camera frame (superscript C) are then

$$\mathbf{p}_A^{\mathrm{C}} = \mathbf{t} + \mathbf{R}\mathbf{p}_A, \qquad \mathbf{p}_B^{\mathrm{C}} = \mathbf{t} + \mathbf{R}[\mathbf{p}_B + \delta(P)\mathbf{e}_z]. \tag{S10}$$

Equation (S10) separates internal deformation from rigid-body motion: $\mathbf{R}$ and $\mathbf{t}$ act collectively on both magnets, whereas pressure changes only the position of B relative to A. The magnetic moments of both magnets rotate with $\mathbf{R}$, assuming that membrane deformation does not appreciably rotate the moment of B.

Stacking the outputs of the $N = 32$ triaxial sensors gives the 96-dimensional array form of equation (S4), $\mathbf{y} \approx \mathbf{h}(P, \mathbf{q}) + \mathbf{B}_{\text{geo}}$, where $\mathbf{h}$ is the field generated by the two magnets and $\mathbf{B}_{\text{geo}}$ is the geomagnetic field. The reference sensor mainly measures $\mathbf{B}_{\text{geo}}$. Subtracting its reading from every array sensor removes $\mathbf{B}_{\text{geo}}$ and yields the compensated measurement $\tilde{\mathbf{y}} \approx \mathbf{h}(P, \mathbf{q})$; the approximation applies when the implant-generated field at the reference sensor is negligible, as experimentally established in Supplementary Fig. 27.

*6.3 Separation of pressure and rigid-body motion*

Linearizing $\tilde{\mathbf{y}}$ around a nominal state ($P_0$, $\mathbf{q}_0$) gives

$$\Delta\tilde{\mathbf{y}} \approx \mathbf{j}_P \Delta P + \mathbf{J}_q \Delta\mathbf{q}, \qquad \text{(S11)}$$

where $\mathbf{j}_P = \partial\mathbf{h}/\partial P$ is the array-level signature of a small pressure change and each column of $\mathbf{J}_q = \partial\mathbf{h}/\partial\mathbf{q}$ is the signature of a small translation or rotation. Pressure is locally distinguishable from pose variation when $\mathbf{j}_P$ contains a component outside the space spanned by the columns of $\mathbf{J}_q$:

$$\mathbf{j}_P \notin \text{col}(\mathbf{J}_q). \qquad \text{(S12)}$$

The physical origin of this condition can be illustrated by axial displacement. Let $\mathbf{J}_A$ and $\mathbf{J}_B$ denote the derivatives of the array measurement with respect to the camera-frame positions $\mathbf{p}_A^{\text{C}}$ and $\mathbf{p}_B^{\text{C}}$. From equation (S10), pressure moves only B along $\mathbf{R}\mathbf{e}_z$, so $\mathbf{j}_P \propto \mathbf{J}_B\mathbf{R}\mathbf{e}_z$. In contrast, axial rigid-body translation moves only B in a single-magnet device but both magnets in the dual-magnet device:

$$\mathbf{j}_{t_z}^{\text{single}} = \mathbf{J}_B\mathbf{R}\mathbf{e}_z, \qquad \mathbf{j}_{t_z}^{\text{dual}} = (\mathbf{J}_A + \mathbf{J}_B)\mathbf{R}\mathbf{e}_z. \qquad \text{(S13)}$$

In the single-magnet case, $\mathbf{j}_P \parallel \mathbf{j}_{t_z}^{\text{single}}$, so a pressure-induced displacement may be indistinguishable from axial rigid-body translation, consistent with Supplementary Fig. 28a. In the dual-magnet case, the contribution $\mathbf{J}_A$ is present during rigid-body motion but absent during pressure-induced deformation. This contribution breaks the proportionality, giving $\mathbf{j}_P \nparallel \mathbf{j}_{t_z}^{\mathrm{dual}}$. The same principle extends to lateral translations, rotations and more general multi-magnet architectures.

*6.4 Spatial MF mapping for separation of pressure and motion*

A point measurement reduces the magnetic response to a few scalar values and can lose the spatial features that distinguish pressure from motion. In contrast, an array of $N$ triaxial sensors provides $3N$ simultaneous measurements of the MF distribution. For one pressure variable and a six-degree-of-freedom pose, $3N \geq 7$ is a necessary, although not sufficient, dimensional condition for joint local estimation. The 32-sensor MF camera provides 96 measurements and satisfies it with substantial redundancy. The spatially distributed measurements retain field gradients and multiaxial spatial relationships that would be lost in a single-point measurement.

**Supplementary Note 7. Advantages of data-driven reconstruction for motion-tolerant MF sensing**

The spatial magnetic-field distribution measured by the MF camera depends simultaneously on the target biomechanical variable, the relative pose between BioBeacon and the MF camera, and the properties of the fabricated device and MF camera. This measurement can be generally expressed as

$$\mathbf{y} = \mathbf{h}(x, \mathbf{q}; \boldsymbol{\theta}) + \mathbf{B}_{\mathrm{geo}}, \qquad \text{(S14)}$$

where $x$ and $\mathbf{q}$ are defined as in equation (S1), and $\boldsymbol{\theta}$ represents device-specific parameters.

An analytical reconstruction would require an explicit and sufficiently accurate description of the forward mapping $\mathbf{h}$. Small deviations in magnetic moment, magnet alignment, structural dimensions, material properties, or channel responses could propagate into the reconstructed biomechanical value.

Data-driven reconstruction avoids explicitly estimating each of these intermediate parameters. Calibration data acquired over different biomechanical states and relative poses are used to directly learn the inverse mapping

$$\hat{x} = \mathcal{G}_{\varphi}(\mathbf{y} - \mathbf{B}_{\mathrm{geo}}), \tag{S15}$$

where $\mathbf{B}_{\mathrm{geo}}$ is removed using the reference sensor (Supplementary Note 5). Here, $\mathcal{G}_{\varphi}$ denotes the trained reconstruction model. Because this mapping is learned from experimentally measured MF distributions, it directly incorporates the nonlinear response and device-specific variations of the fabricated system.

#### Supplementary Note 8. Sampling strategy and configuration of pose–pressure datasets

##### *8.1 Dataset design and pressure sampling*

In order to collect a comprehensive dataset for deep learning that decouples pressure changes from disturbances caused by changes in the relative position and orientation between the Hall sensor array and the implanted device, the training set should include the readings of magnetic field from the Hall sensor array when pressure and relative pose ($x$, $y$, $z$, $\theta$, $\varphi$) differ.

We program the output pressure as 0, 0.3, 0.6, 0.9, 1.2, 2.4, 3.6, 4.8, 6.0 kPa during collection of the training set, using finer pressure increments within 0–1.2 kPa to support accurate decoding of small pressure changes, while maintaining training coverage up to 6 kPa to capture elevated ICP.

##### *8.2 Training dataset acquisition*

We set the movement range of the training set within a volume ($x$, $y$, and $z$ axes) of 7 mm × 7 mm × 3.5 mm with a step of 0.5 mm, resulting in 1800 3-D positions uniformly distributed in the volume. The vertical calibration starts at $z$ = ~3 mm ($z$ denotes the vertical separation between the lower surface of the MF camera and the upper surface of the BioBeacon), so the 3.5-mm vertical span corresponds to a $z$ range of ~3–6.5 mm. The 3-D positions are repeated after changing the pitch angle ($\varphi$) within ±3° with a step of 1.5°, which results in a 4-D position set of 9000 states. Then the 4-D position sets are repeated after changing the roll angle ($\theta$) within ±3° with a step of 1.5°, resulting in a 5-D position set of 45000 states. The rotational symmetry of the device about the yaw axis $\psi$ eliminates the need for yaw-angle sampling. For every motion and rotation, the robotic arm pauses moving every 1 s after every step of 0.5 mm. Every time the robotic arm stops, the digital output pin on the robotic arm generates a pulse signal that triggers data acquisition of the Hall sensor array. For all 9 pressures, 405000 different states are collected. For the training set with an extended vertical range, the vertical span is increased from 3.5 to 6.5 mm, yielding a translational volume of 7 mm × 7 mm × 6.5 mm (corresponding to $z$ = ~3–9.5 mm), with the translational step size, orientations and pressure levels staying the same.

*8.3 Validation and test dataset acquisition*

To evaluate interpolation at previously unseen states within the training domain, the first validation trajectory covers a translational volume of 6 mm × 6 mm × 2 mm with a step size of 2 mm along $x$ and $y$ axes and 1 mm along the $z$ axis. Both the pitch angle $\varphi$ and the roll angle $\theta$ vary within ±2° with an increment of 2°. A 0.25 mm offset shifts the $x$, $y$, and $z$ coordinates away from the training grid. Thus, the validation states remain entirely within the training range but do not coincide with complete training poses. The validation pressures are set to 0, 0.2, 0.4, 0.6, 0.8, 1.0, 1.8, 3.0, 4.2, 5.4 kPa. This design prevents identical pressure–pose combinations from appearing in both datasets and evaluates spatial, angular, and pressure interpolation.

To test performance over a larger pose range, the test trajectory expands the translational volume to 8 mm × 8 mm × 6 mm with a step size of 1 mm along all three axes. Both pitch $\varphi$ and roll $\theta$ vary within ±2° with an increment of 2°. A 0.25 mm offset shifts the $z$-coordinate planes away from the training grid. These offsets prevent any complete five-dimensional pose from matching a training state. This extended-range dataset includes poses both inside and outside the training domain and evaluates model generalization across an enlarged spatial range.

#### Supplementary Note 9. Effect of calibration range on the sensing workspace

Supplementary Fig. 32 compares the pressure-decoding error maps of models trained with the original and expanded vertical calibration ranges, evaluated on the same extended-range test dataset described in Supplementary Note 8. Fig. 3d shows the first five $z$ planes, spanning 4 mm, whereas Supplementary Fig. 32 presents all seven planes. Within the planes shown in Fig. 3d, decoding errors remain low inside the training-covered region (MAE, 0.034 kPa; $R^2$, 0.9992) but increase near the peripheral regions outside the calibrated range (MAE, 0.35 kPa; $R^2$, 0.91). With the original vertical training range of 3.5 mm ($z$ = ~3–6.5 mm), errors increase markedly in the final three planes ($z$ = 7.25–9.25 mm), which extend beyond the calibrated range (Supplementary Fig. 32a).

Extending the vertical training range from 3.5 to 6.5 mm (corresponding to an expanded $z$ range of ~3–9.5 mm), while maintaining the same lateral coverage, reduces errors at larger camera–implant separations (Supplementary Fig. 32b). For $z \geq 7.25$ mm and within the calibrated lateral region, the mean absolute error (MAE) decreases from 1.12 to 0.162 kPa, and $R^2$ increases from 0.2152 to 0.9829, enabling reliable readout at camera–implant distances of ~10 mm. Errors persist at the lateral edges, where calibration is not expanded. These results indicate that broader calibration can enlarge the reliable sensing workspace and extend the readout distance.

**Supplementary Note 10. Equivalent displacement estimation accuracy of BioBeacon-PA**

BioBeacon-PA encodes pressure through pressure-induced deformation of the flexible membrane, which changes the axial separation between the movable and fixed magnets and thereby alters the spatial MF distribution. To quantify the decoding performance in terms of this internal configuration change, an independent pressure–displacement calibration is performed under an optical microscope (Supplementary Fig. 36a–c). The axial displacement of the magnet-bearing membrane is manually measured at pressures from 0 to 6 kPa in three independent pressure-loading experiments. The mean displacement values are fitted using a third-order polynomial constrained to pass through the origin, yielding the empirical pressure–displacement calibration function (Supplementary Fig. 36d):

$$d(P) = 140.502P - 17.6199P^2 + 1.10514P^3, \qquad \text{(S16)}$$

where $P$ is the applied pressure in kPa and $d$ is the corresponding axial displacement in micrometres. Over the complete calibration range of 0–6 kPa, the measured displacement reaches 443 μm. Within the pressure range evaluated by the decoding model (0–5.4 kPa), the corresponding internal displacement extends from 0 to 419 μm.

For each validation sample $i$, the reference pressure $P_i$ and the predicted pressure $\hat{P}_i$ are independently converted into equivalent displacements using equation (S16), giving the error in equivalent displacement estimation

$$e_{d,i} = d(\hat{P}_i) - d(P_i). \tag{S17}$$

The displacement mean absolute error (MAE) and root-mean-square error (RMSE) are then calculated over all samples. This pointwise conversion preserves the nonlinear pressure–displacement response of BioBeacon-PA instead of approximating the displacement error using a constant sensitivity. Across all 4440 validation samples acquired at different relative positions and orientations between BioBeacon-PA and the MF camera, the equivalent displacement estimates show an MAE of 2.74 μm and an RMSE of 3.64 μm. These values correspond to 0.65% and 0.87%, respectively, of the 419-μm displacement range.

The signed and absolute errors in equivalent displacement estimation are further evaluated separately at each applied pressure (Supplementary Fig. 36e,f). The signed error distributions remain near zero across the tested pressure levels, with only small pressure-dependent offsets (-0.26 μm; Supplementary Fig. 36e). The pressure-specific displacement MAEs, indicated by the mean values of the absolute error distributions in Supplementary Fig. 36f, remain between 2.00 and 3.70 μm throughout the tested operating range, demonstrating micrometre-scale equivalent accuracy at different internal magnet separations. The moderately higher displacement errors at low pressures mainly reflect the greater local slope of the pressure–displacement calibration curve, which converts a given pressure error into a larger equivalent displacement error.

Conventional imaging modalities, including computed tomography (CT), magnetic resonance imaging (MRI) and ultrasonography, can visualize implant geometry but require dedicated imaging equipment and involve trade-offs among spatial resolution, temporal resolution and imaging depth that limit continuous deformation tracking in freely behaving subjects. Analytical magnetic localization avoids tomographic imaging but relies on explicit forward models and accurately calibrated magnetic moments, sensor geometries and device configurations. Such assumptions become restrictive for deformable multi-magnet implants, in which material inhomogeneity, fabrication deviations and MF superposition can bias the inferred internal displacement. By directly learning experimentally measured spatial MF patterns, the present approach combines micrometre-scale equivalent displacement estimation accuracy with continuous wireless readout and tolerance to variations in the external position and orientation of BioBeacon relative to the MF camera. External reader volumes and reported localization errors or imaging resolutions for representative magnetic-localization and medical-imaging systems are summarized in Supplementary Table 3.

**Supplementary Note 11. Mapping of strain, shear, and bending**

In the implantable BioBeacon-S configuration, a planar b-DCPU membrane carrying four discrete magnets converts tensile strain into rectangular expansion and shear into parallelogram distortion (Supplementary Fig. 10). In the ex vivo demonstration, four discrete magnets (diameter: 2 mm; pitch: 5 mm) are bonded directly to a rabbit stomach. This configuration uses the tissue itself as a deformable substrate, thereby minimizing constraints on its native deformation. Tensile expansion increases the inter-magnet distances, whereas shear changes the angles of the initially square layout. Gastric expansion transforms the initial square arrangement into an expanded and tilted quadrilateral and redistributes the magnetic features in the MF map (Supplementary Fig. 37a). A mapping-based analysis of the four magnetic features provides an estimate of the angular and area distortion.

BioBeacon-B (3 mm × 10 mm), a b-DCPU film containing aligned FeNWs, is imaged with the 256-sensor MF camera (pixel number: 16 × 16; pitch, 2.5 mm; area, 39 mm × 39 mm; Supplementary Figs. 6 and 7). Under through-thickness magnetization, bending reorients the local magnetic moments and shifts the smoothed boundary between positive and negative $B_z$ regions in accordance with the bent profile (Supplementary Fig. 37b). Under longitudinal magnetization, tensile deformation alters the positions and shapes of these field regions, which qualitatively reflect the extent and spatial distribution of the deformation (Supplementary Fig. 37c). Such encoding versatility expands the potential utility of BioBeacon for measuring deformation across diverse tissues and anatomical structures.

**Supplementary Note 12. Motion-intensity analysis and activity patterns after stroke**

Throughout the monitoring period, both rats remain unrestrained and are free to move within their cages. The relative motion intensity, calculated from changes in the reference-MF vector between successive acquisitions (see Methods), provides a continuous measure of their spontaneous activity (Fig. 5j). Over the 120-hour recording, the sham-operated rat exhibits mean motion intensities of 43.49% during the daytime (08:00–20:00; e.g., 13.24–25.24 h on the trace) and 63.04% at night (20:00–08:00; e.g., 1.24–13.24 h on the trace). The corresponding values in the t-MCAO rat are 73.00% (e.g., 17.30–29.30 h on the trace) and 78.35% (e.g., 5.30–17.30 h on the trace), respectively.

**Supplementary Note 13. System-level trade-offs in bioresorbable implantable monitoring**

Temporary implantable monitoring requires continuous measurements over a clinically relevant period and subsequent elimination of implanted materials. Active implants incorporate electronics for signal conditioning and wireless telemetry, but generally rely on non-bioresorbable electronics, encapsulation and power-management components. Passive bioresorbable implants simplify the implanted structure and place readout outside the body, but depend more on the transcutaneous interrogation link.

In passive wireless systems, motion can alter the position and orientation of the implant relative to the reader and can thereby produce signal changes unrelated to the target physiological variable. Tethers or constrained reader geometries can stabilize this link but restrict natural behaviour. Continuous monitoring during free behaviour therefore requires motion tolerance.

BioBeacon converts biomechanical deformation into changes in its internal magnetic configuration without implanted batteries, active electronics or telemetry. An external MF camera captures spatial magnetic responses for data-driven decoding. Negligible tissue-dependent attenuation limits transmission variations during motion, while spatial MF measurements distinguish internal deformation from changes in implant–camera position and orientation. These features support continuous quantitative monitoring during free behaviour with a fully bioresorbable implant.

The radar plots in Supplementary Fig. 47 compare BioBeacon with representative implantable monitoring systems across five dimensions: implant size, external reader size, continuous monitoring during free behaviour, implant bioresorbability and implant power requirements. Supplementary Table 4 provides the detailed scoring criteria.

### Supplementary Tables

**Supplementary Table 1. Detailed functions of different configurations of BioBeacon**

| Abbreviation | Full name | Function |
|---|---|---|
| BioBeacon-P | Bioresorbable permanent-magnet beacon for sealed-cavity pressure sensing | Measures absolute pressure using a sealed cavity as an internal reference, where pressure-induced membrane deformation displaces a central magnet axially relative to a fixed annular magnet and alters the spatial MF distribution. |
| BioBeacon-PA | Bioresorbable permanent-magnet beacon for atmosphere-referenced pressure sensing | Measures differential pressure across the membrane in a compact, cavity-free configuration by encoding pressure-induced deformation in the spatial MF distribution, providing atmosphere-referenced gauge-pressure sensing when one membrane surface is exposed to air. |
| BioBeacon-B | Bioresorbable permanent-magnet beacon for bending sensing | Demonstrates continuous-deformation encoding in a flexible FeNW–polyurethane film, where bending reorients the local magnetic moments and spatially nonuniform tensile deformation produces corresponding local variations in the spatial MF pattern. |
| BioBeacon-S | Bioresorbable permanent-magnet beacon for strain and shear sensing | Resolves in-plane deformation encoding using a four-magnet layout, where tensile strain changes the inter-magnet spacing and shear changes the included angles between the magnets, producing corresponding changes in the spatial MF distribution. |

**Supplementary Table 2. Remanence and coercivity of representative macroscopic magnetic structures made from bioresorbable magnetic materials**

| No. | Magnetic component | Matrix | $H_c$ (Oe) | $M_r$ (emu g$^{-1}$) |
|---|---|---|---|---|
| This work (aligned) | Fe nanowires (Ø ~60 nm) | PCL | ~400 | ~30 |
| This work (unaligned) | Fe nanowires (Ø ~60 nm) | PCL | ~400 | ~15 |
| 1 | $Fe_3O_4$ NPs (~10 nm)[1] | Gelatin | ~0 | ~0 |
| 2 | $Fe_3O_4$ NPs (~50 nm)[2] | PLLA | 118 | 9.1 |
| 3 | $Fe_3O_4$ particles (<5 μm)[3] | PCL | 114 | 2.17 |
| 4 | γ-$Fe_2O_3$ NPs (~10 nm)[4] | PHEMA | 0.52 | 0.0036 |
| 5 | α-$Fe_2O_3$ NPs (~87 nm)[5] | Chitosan | ~160 | 0.00425 |
| 6 | ε-$Fe_2O_3$ NPs (14–34 nm)[6] | $SiO_2$ | 18170 | ~0.82 |
| 7 | Fe particles (~40 μm)[7] | PLA | <10 | ≤0.2 |
| 8 | Fe (sintered)[8] | — | ~1.3 | 0.066 |
| 9 | Fe-doped HA NPs (~13 nm)[9] | PCL | ~15 | ~0.25 |
| 10 | Metal-Free Molecular Magnets[10] | PNIPAM | 253 | ~0.09 |

**Supplementary Table 3. External reader volume and reported localization error or imaging resolution of representative magnetic localization and medical imaging systems**

| No. | Device | Size of device ($cm^3$) | Error or resolution (μm) |
|---|---|---|---|
| ★ | This work (MF camera) | ~4.5 | 2.74 |
| 1 | Intermediary patch[11] | ~30 | ~170 |
| 2 | Magnetomicrometry[12] | ~50 | ~229 |
| 3 | Myokinetic array[13] | ~100 | ~180 |
| 4 | SKYSCAN 1276[14] | ~1067144 | ~6 |
| 5 | Revvity Quantum GX[15] | ~2625700 | ~5 |
| 6 | SOMATOM Force[16] | ~4230883.5 | ~240 |
| 7 | 80-mT portable MRI[17] | ~114250 | ~1300 |
| 8 | MAGNETOM Vida[18] | ~10906548 | ~400 |
| 9 | Bruker BioSpec Maxwell 7[19] | ~5359296 | ~94 |
| 10 | ACUSON Sequoia + HLX[20] | ~688080 | ~100 |
| 11 | Butterfly iQ+[21] | ~319 | ~1600 |
| 12 | USoP[22] | ~30 | ~229 |

**Supplementary Table 4. Scoring instruction for the radar plots**

| Score | Implant size ($mm^3$) | External reader size ($cm^3$) | Continuous monitoring in freely behaving subjects | Implant bioresorbability | Implant power requirement |
|---|---|---|---|---|---|
| 1 | ≥1000 | ≥5000 | Monitoring requires a percutaneous implant-to-external connection or direct wired readout | The implanted system is non-bioresorbable | Active implanted components powered by an implanted battery or transcutaneous wiring |
| 2 | 100–<1000 | 100–<5000 | Wireless sensing is demonstrated, but measurements are intermittent, session-based or acquired only at discrete time points | Only part of the implanted system is bioresorbable, with essential permanent components remaining | Active implanted components without an implanted battery; energy is supplied by wireless power transfer |
| 3 | 20–<100 | 5–<100 | Continuous wireless time-series recording is demonstrated, but subject movement, posture or reader–implant geometry remains constrained | The main sensing structure is bioresorbable, but minor non-bioresorbable implanted components remain | Passive implant without active electronics, but external RF, acoustic or optical excitation is required |
| 4 | <20 | <5 | Continuous untethered monitoring is demonstrated in freely behaving subjects without fixed reader–implant alignment | All implanted components required for operation are bioresorbable | Fully passive implant requiring no external energy delivery to the implant |

### Supplementary Figures

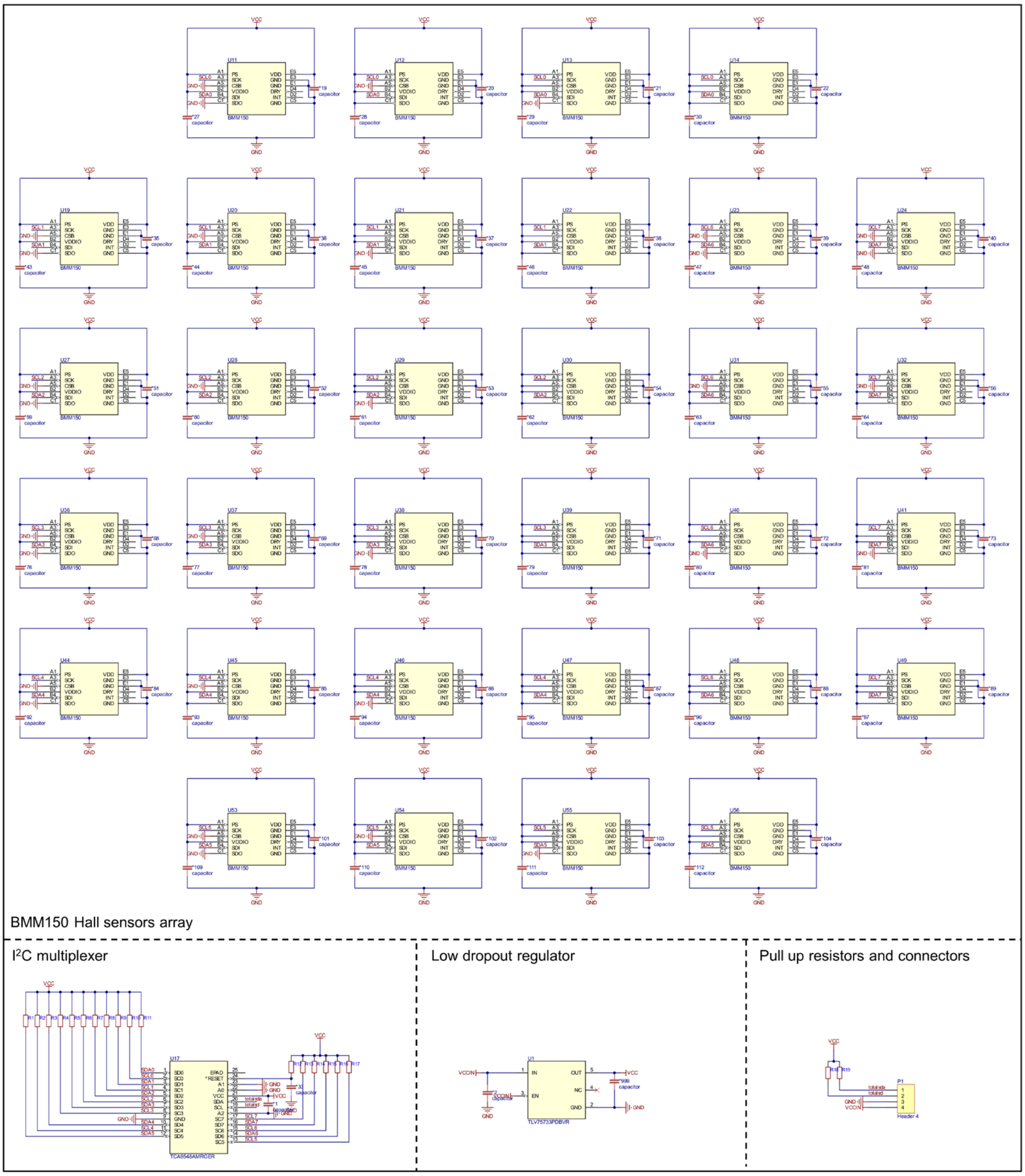

**Supplementary Fig. 1 | Circuit schematic of the Hall-sensor array board.** The circuit comprises 32 BMM150 magnetic sensors, an I$^2$C multiplexer, a low-dropout regulator, pull-up resistors and connectors.

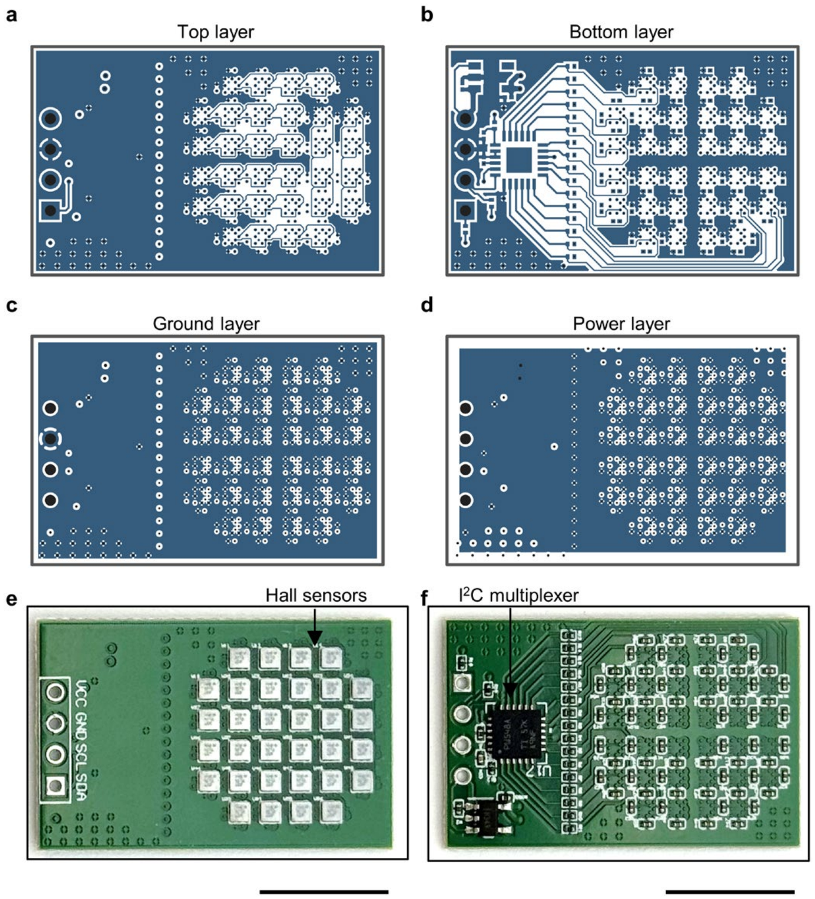

**Supplementary Fig. 2 | PCB layout of the Hall-sensor array board. a–d**, PCB design layers. Black circles indicate drilled holes, dark grey outlines indicate board boundaries, and blue regions indicate copper layers. **e**,**f**, Photographs of the front and back sides of the assembled Hall-sensor array board. Scale bars, 1 cm.

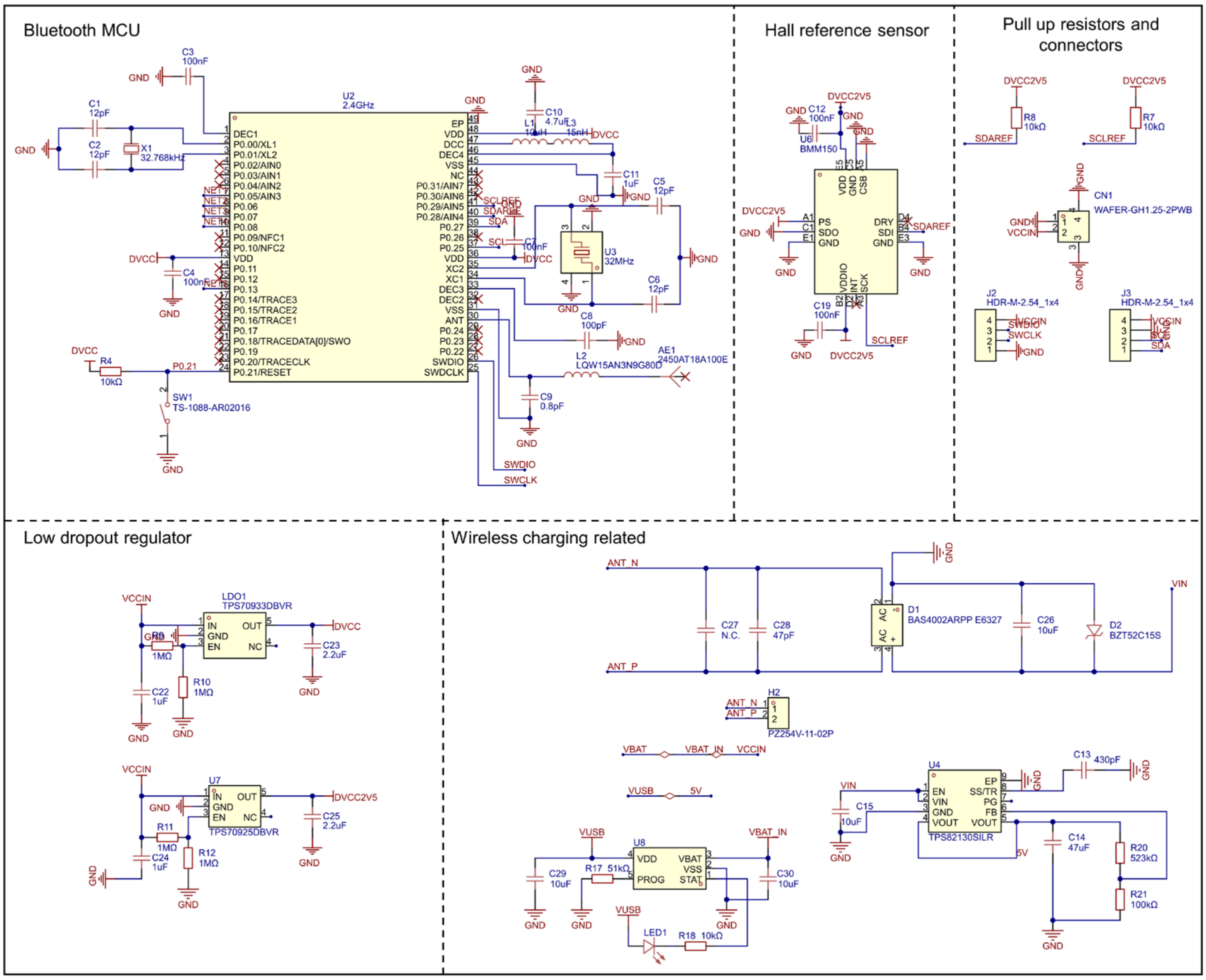

**Supplementary Fig. 3 | Circuit schematic of the BLE control board.** The circuit comprises a Bluetooth microcontroller, reference Hall sensor, voltage regulators, connectors and wireless-power management components.

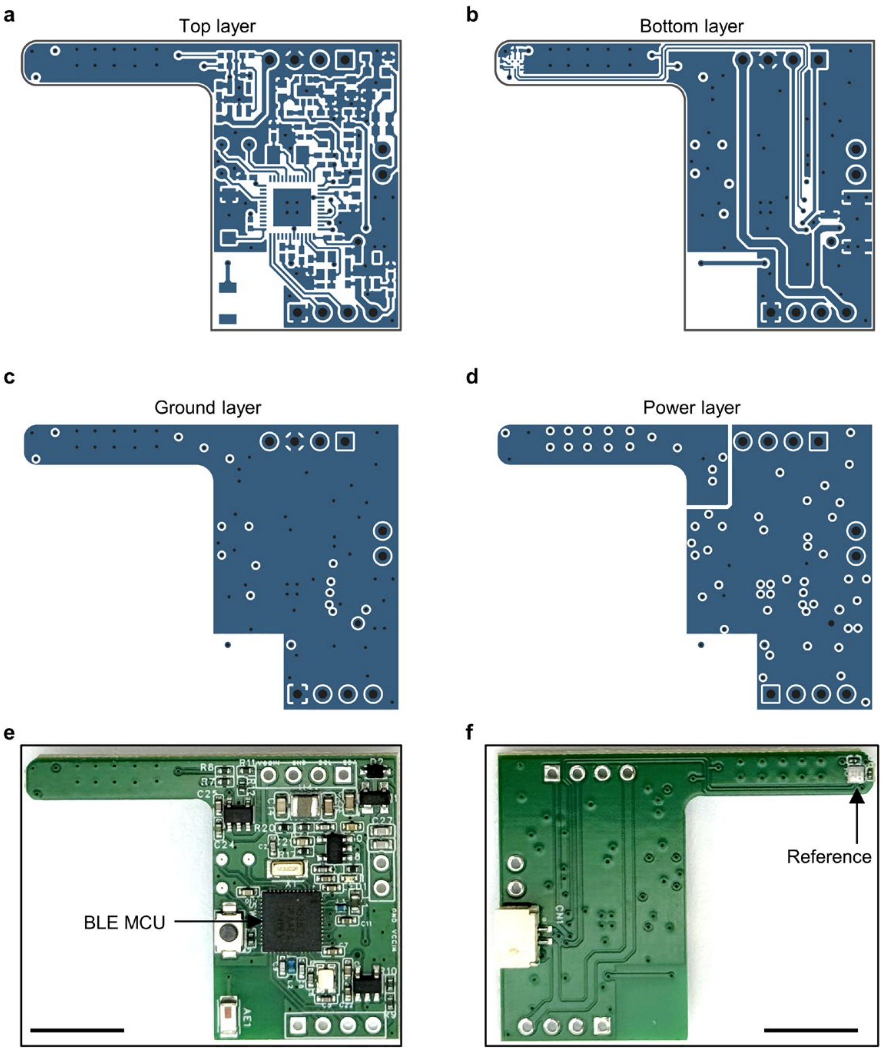

**Supplementary Fig. 4 | PCB layout of the BLE control board. a–d**, PCB design layers. Black circles indicate drilled holes, dark grey outlines indicate board boundaries, and blue regions indicate copper layers. **e**,**f**, Photographs of the front and back sides of the assembled BLE control board. Scale bars, 1 cm.

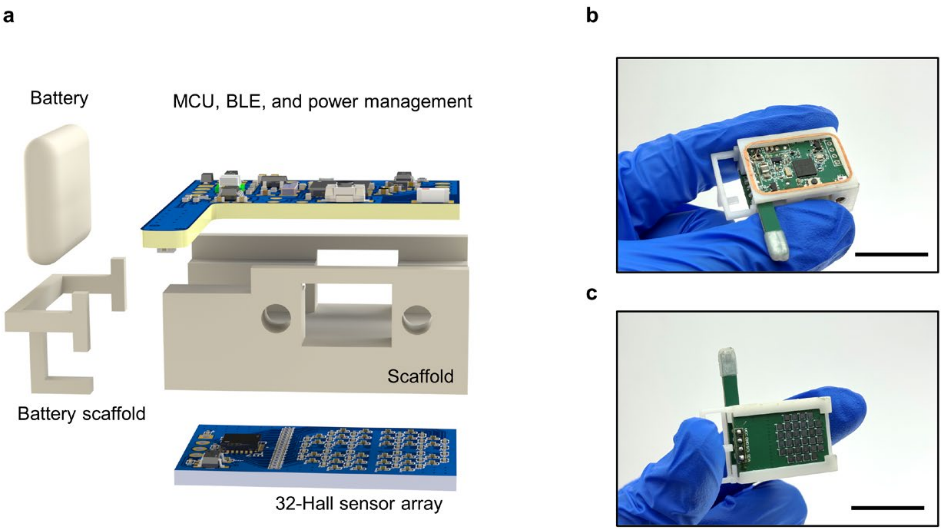

**Supplementary Fig. 5 | Assembly of the wearable magnetic-field camera. a**, Exploded view showing the battery, battery scaffold, control board, supporting scaffold and 32-Hall-sensor array. **b**,**c**, Photographs of the assembled camera. Scale bars, 2 cm.

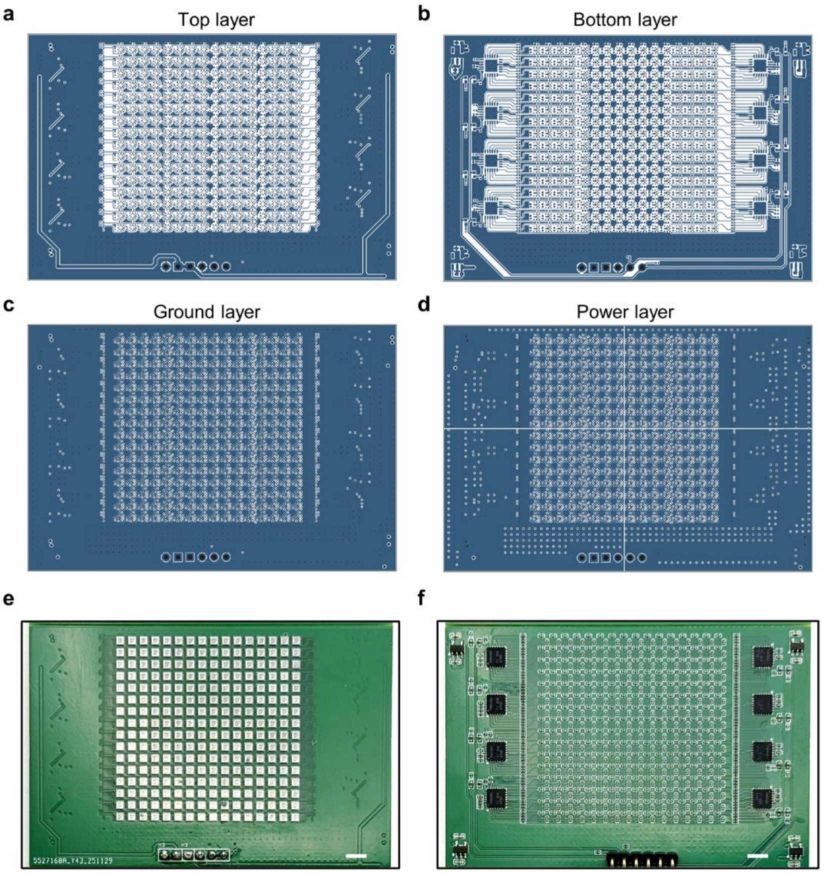

**Supplementary Fig. 6 | PCB layout of the 256-Hall-sensor array. a–d**, PCB design layers. Black circles indicate drilled holes, dark grey outlines indicate board boundaries, and blue regions indicate copper layers. **e**,**f**, Photographs of the front and back sides of the assembled 256-Hall-sensor array board. Scale bars, 4 mm.

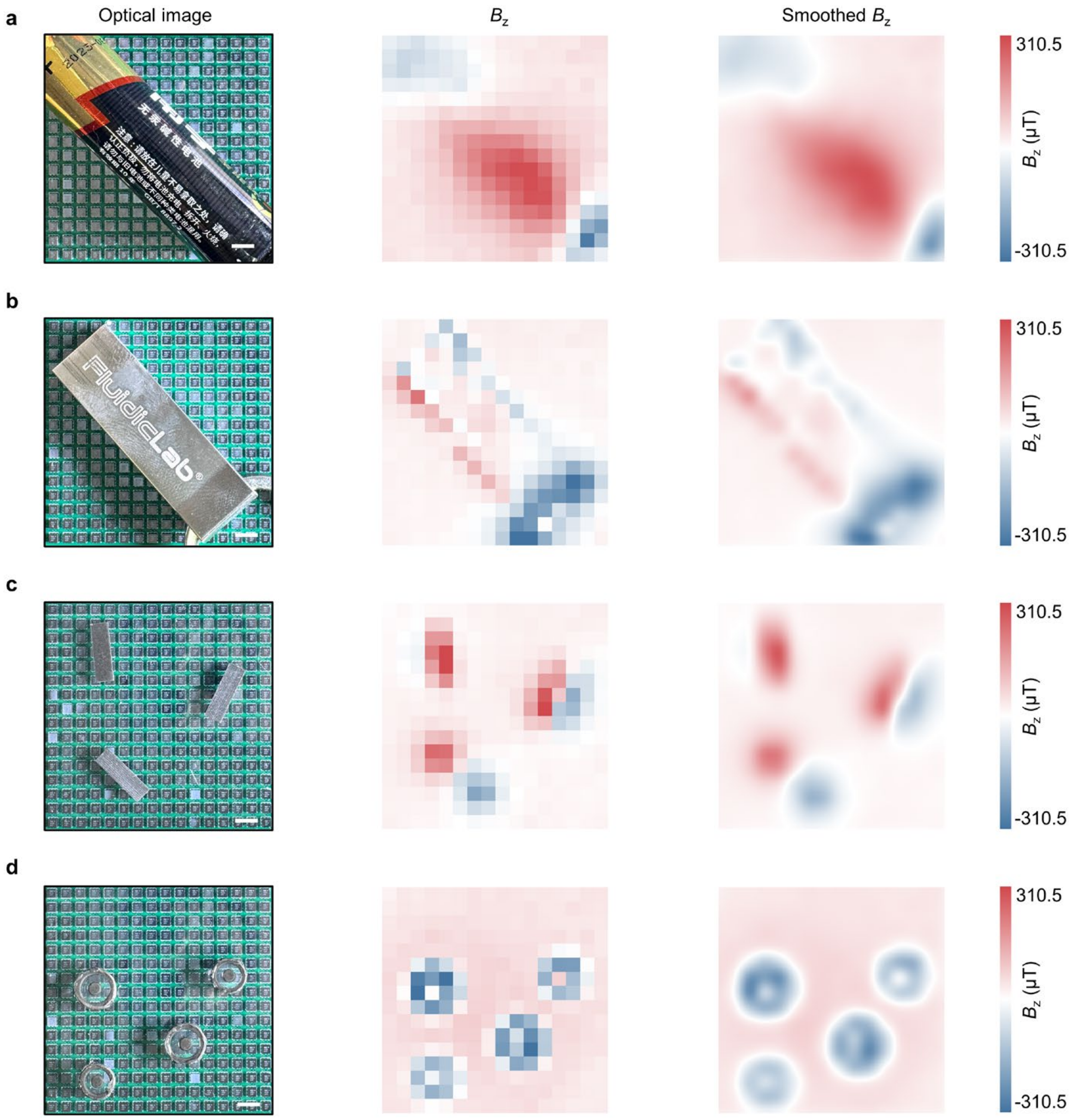

**Supplementary Fig. 7 | Magnetic-field mapping using the 256-sensor camera. a–d**, Optical images and corresponding measured and smoothed $B_z$ maps of four representative magnetic objects. Scale bars, 4 mm.

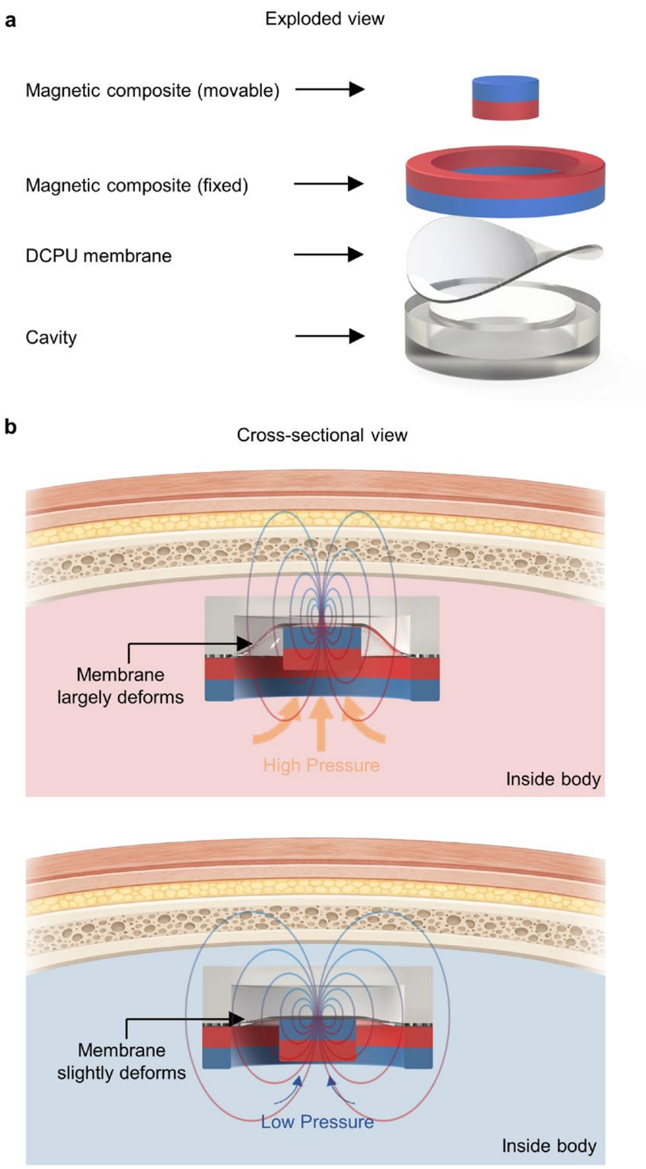

**Supplementary Fig. 8 | Structure and pressure-sensing mechanism of BioBeacon-P. a**, Exploded view of the movable magnet, fixed annular magnet, b-DCPU membrane and cavity. **b**, Cross-sectional views showing pressure-induced membrane deformation and relative magnet displacement. Different biomechanical signals induce different magnetic field (MF) distributions, as illustrated by the MF lines. MF lines are shown schematically for illustration.

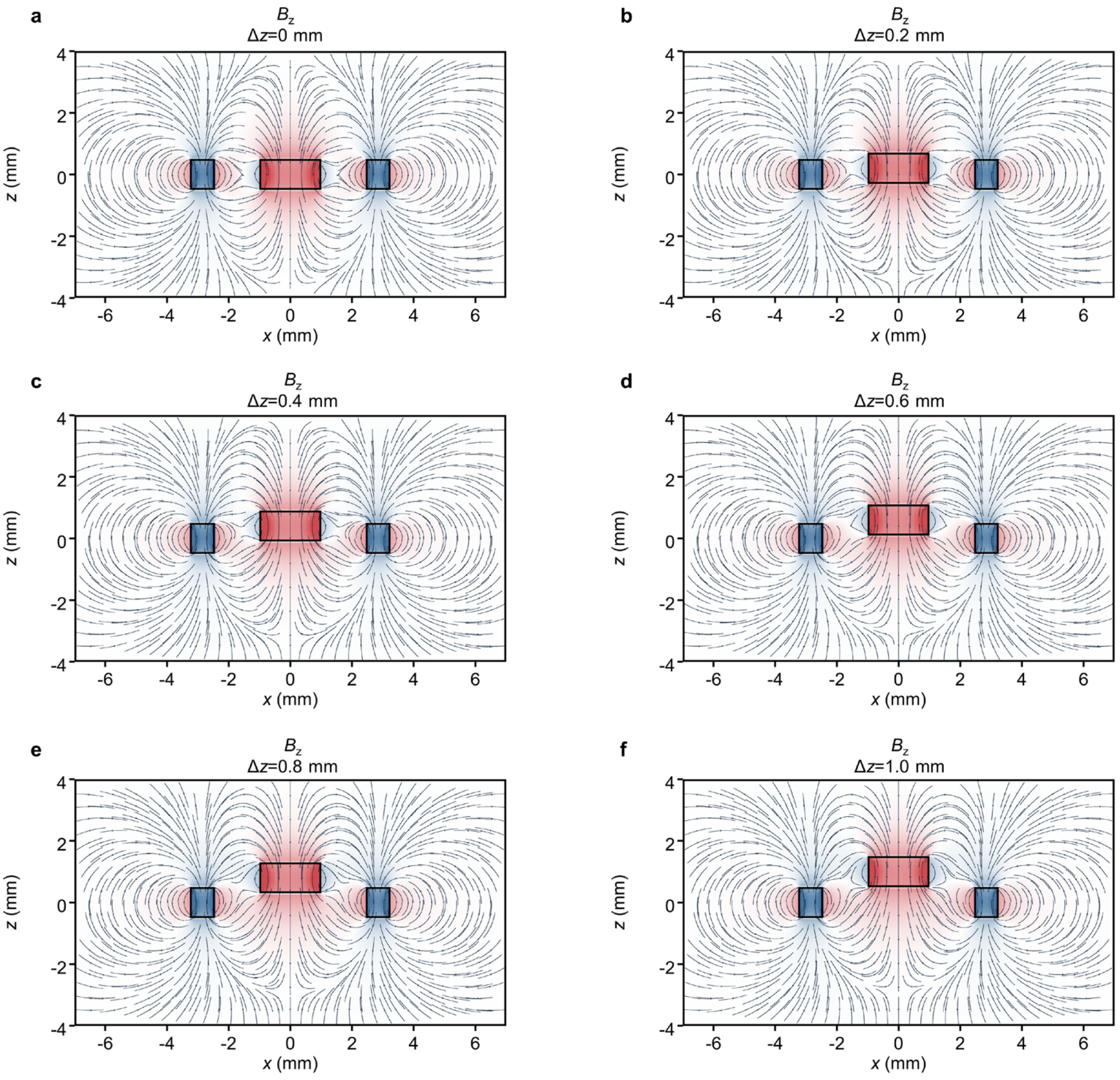

**Supplementary Fig. 9 | Simulated magnetic-field distributions of BioBeacon-P. a–f**, Finite-element-analysis-derived $B_z$ distributions for axial displacements of the movable magnet from 0 to 1.0 mm.

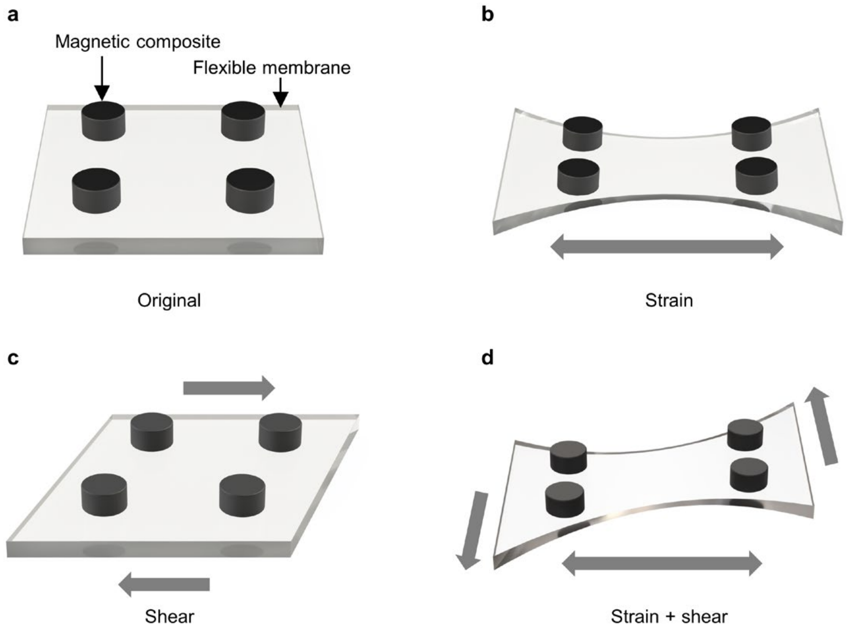

**Supplementary Fig. 10 | Deformation modes of BioBeacon-S. a–d**, Schematics of the original configuration and its deformation under tensile strain, shear and combined strain and shear.

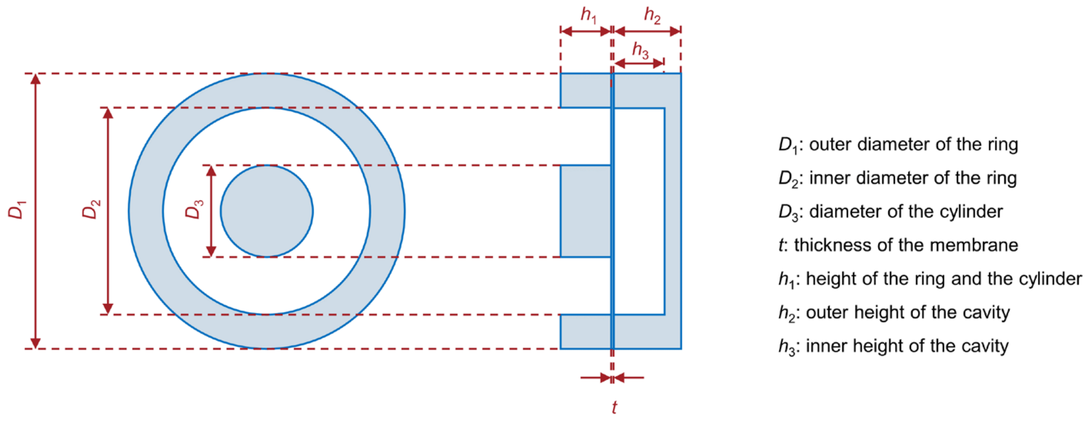

**Supplementary Fig. 11 | Geometric parameters of BioBeacon-P.** The diagram defines the diameters, membrane thickness and cavity dimensions used to describe the device.

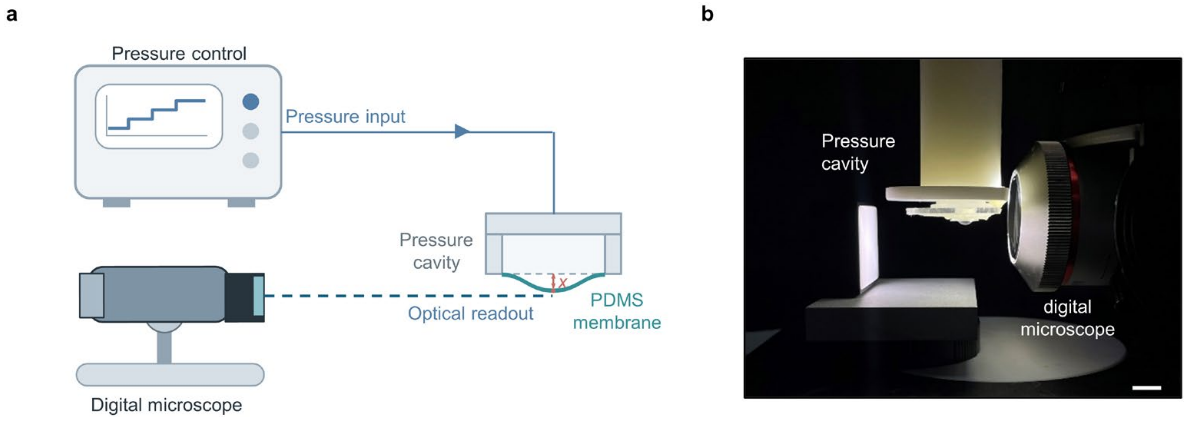

**Supplementary Fig. 12 | Experimental setup for membrane-scaling measurements. a**, Schematic of pressure application and optical measurement of central membrane displacement. **b**, Photograph of the experimental setup. Scale bar, 1 cm.

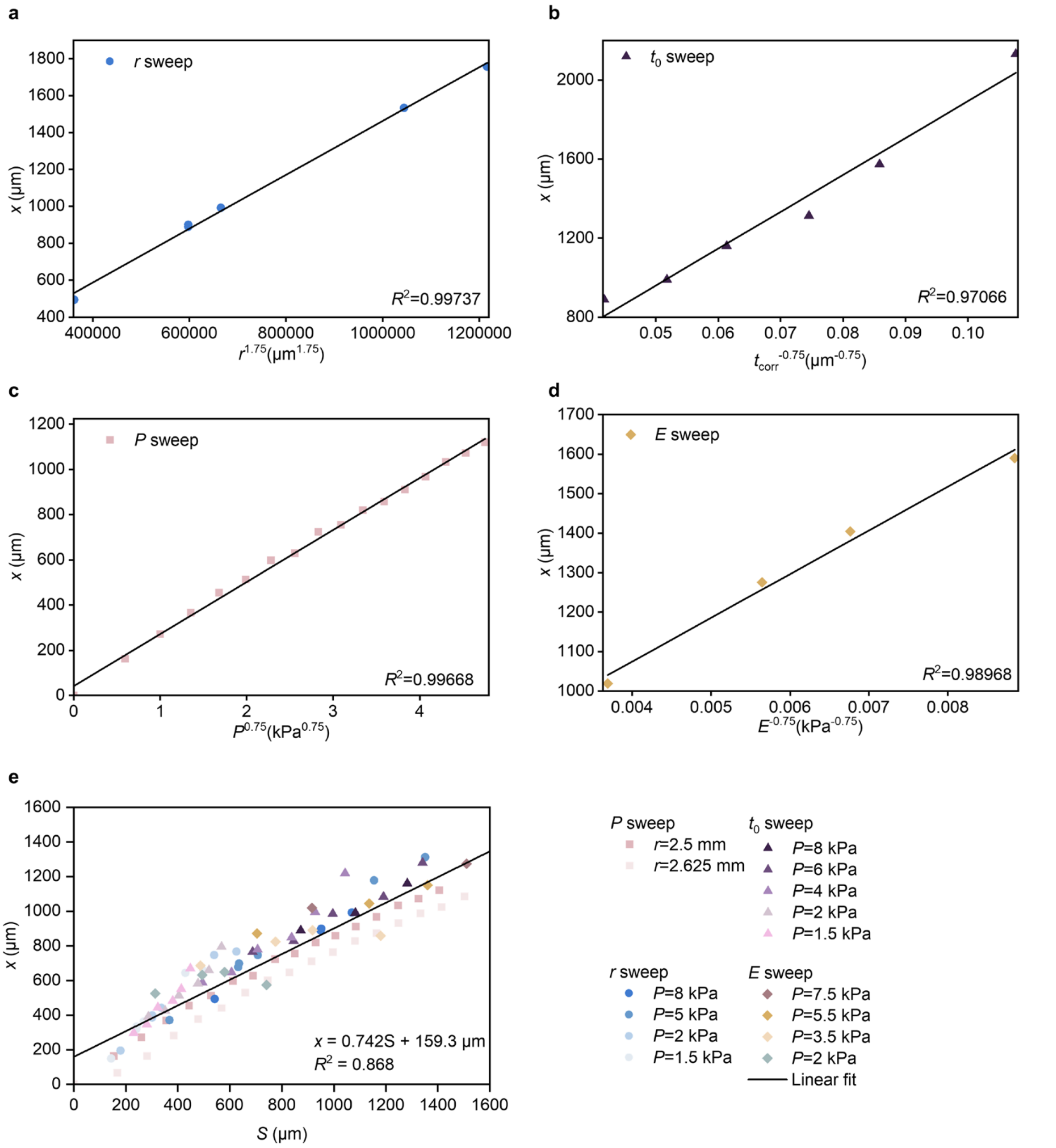

**Supplementary Fig. 13 | Empirical scaling of pressure-induced membrane displacement.** **a**, Central membrane displacement, $x$, plotted against $r^{1.75}$ for the aperture-radius sweep at $P$ = 8 kPa, $E$ = 1.178 MPa and $t_0$ = 43.92 μm. The aperture radius ranges from 1.50 to 3.00 mm, including two separate samples at 2.00 mm. **b**, Central membrane displacement plotted against $t_{corr}^{-0.75}$ for the initial-thickness sweep at $P$ = 8 kPa, $r$ = 2.5 mm and $E$ = 1.178 MPa. The initial membrane thickness ranges from 33.71 to 77.82 μm, where $t_{corr}$ denotes the geometrically corrected membrane thickness. **c**, Central membrane displacement plotted against $P^{0.75}$ for the pressure sweep from 0 to 8 kPa at $r$ = 2.5 mm, $E$ = 1.178 MPa and $t_0$ = 43.92 μm. **d**, Central membrane displacement plotted against $E^{-0.75}$ for the modulus sweep at $P$ = 7.5 kPa and $r$ = 2.5

mm. The membrane modulus ranges from 0.547 to 1.754 MPa, with closely matched initial membrane thicknesses of 46.46–48.74 μm. **e**, Pooled relationship between *x* and the scaling coordinate for 95 measurements from 15 experimental series. For calculating *S*, *P* and *E* are expressed in kPa, whereas *r* and $t_{corr}$ are expressed in μm, giving *S* in μm. Squares, circles, triangles and diamonds denote the pressure, aperture-radius, initial-thickness and modulus sweeps, respectively; colours distinguish the individual experimental series (conditions indicated in the legend). Black lines indicate linear fits with free intercepts; the corresponding $R^2$ values are shown in the panels.

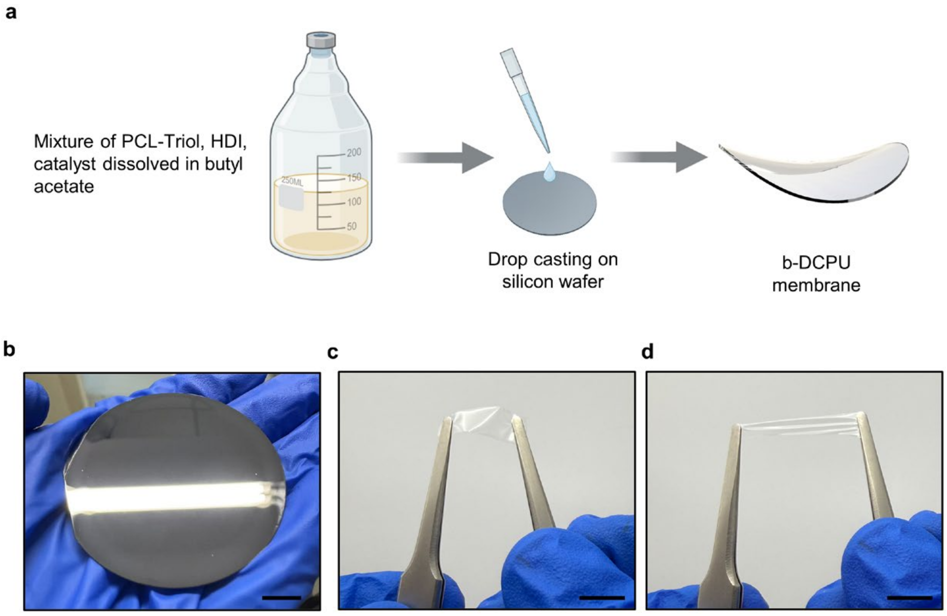

**Supplementary Fig. 14 | Fabrication of the b-DCPU membrane. a**, Schematic of precursor preparation and drop casting. **b**–**d**, Photographs of the resulting membrane on a 2-inch silicon wafer and stretchable free-standing membrane. Scale bars, 1 cm.

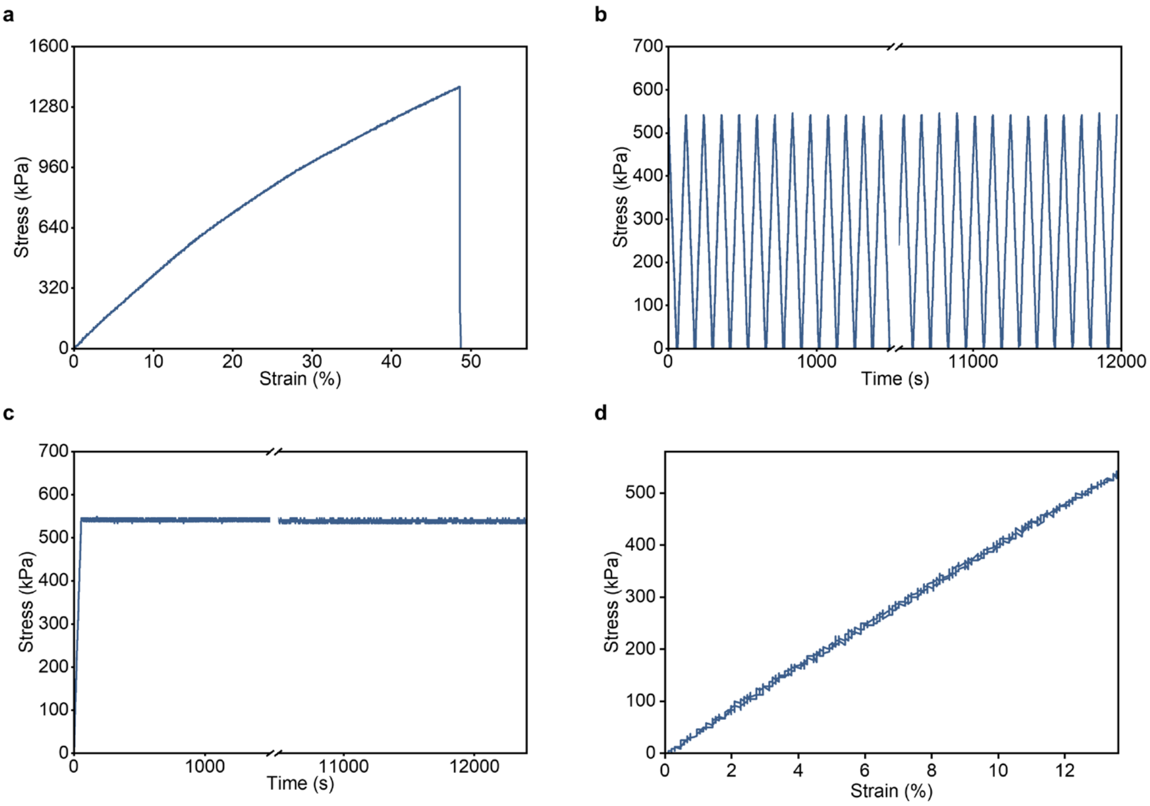

**Supplementary Fig. 15 | Mechanical properties of b-DCPU. a**, Uniaxial stress–strain curve showing fracture at a strain of ~48%. **b**, Stress response during repeated stretching and release over 12000 s, showing stable cyclic performance. **c**, Stress stability under sustained tensile deformation over 12000 s. **d**, Loading–unloading stress–strain curve showing negligible mechanical hysteresis.

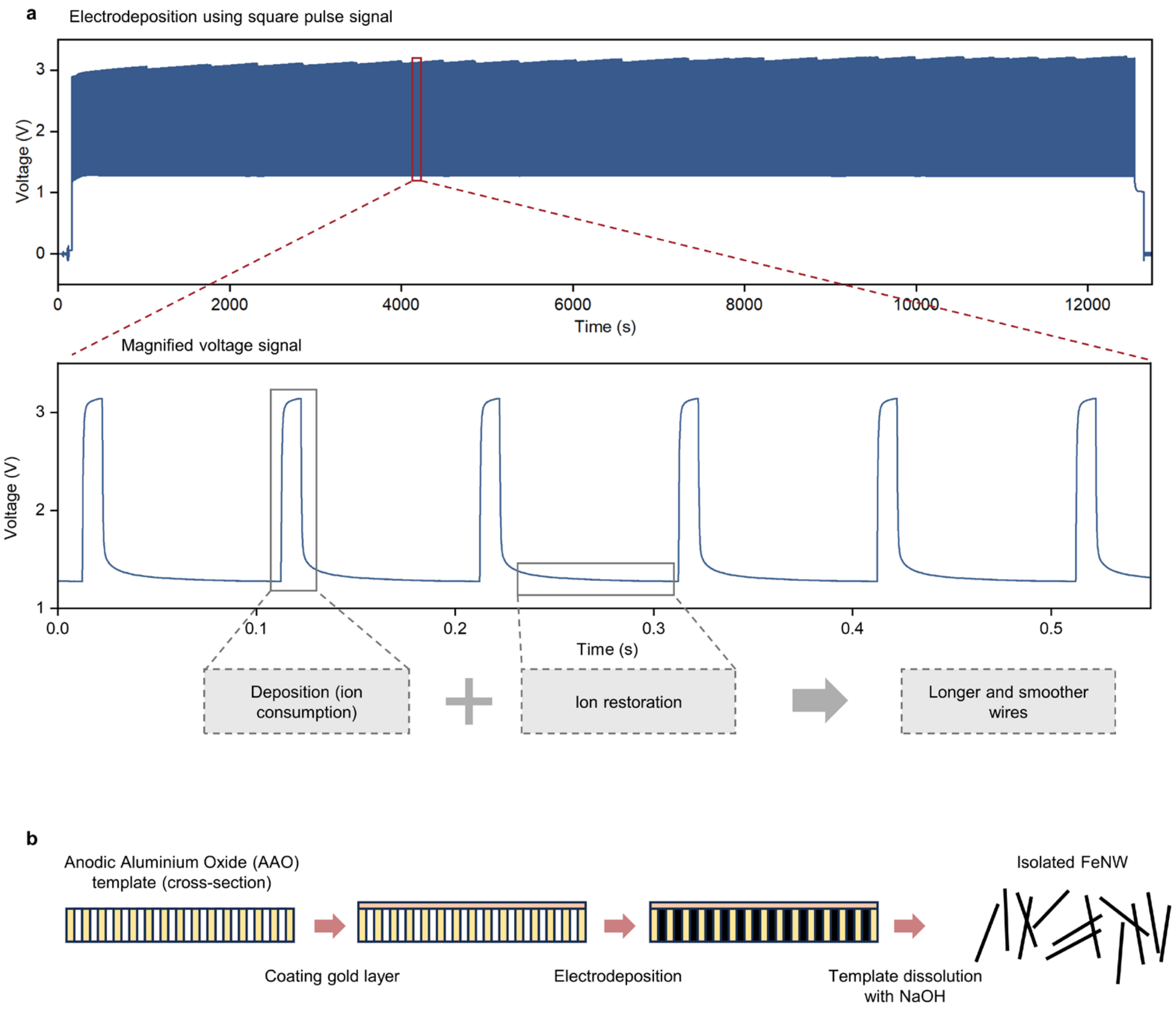

**Supplementary Fig. 16 | Electrodeposition of iron nanowires. a**, Deposition pulse sequence and magnified waveform. **b**, Fabrication process comprising Au coating of the anodic aluminium oxide template, electrodeposition and template dissolution.

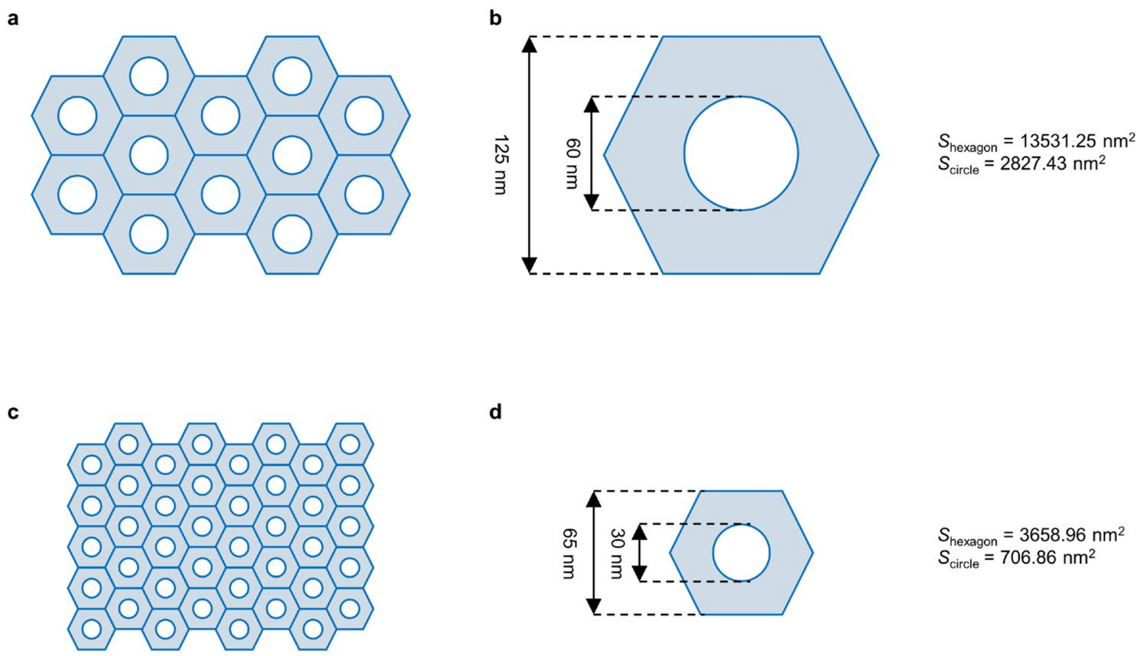

**Supplementary Fig. 17 | Pore geometry of anodic aluminium oxide templates. a–d**, Schematics and unit-cell dimensions of templates with nominal pore diameters of 60 and 30 nm.

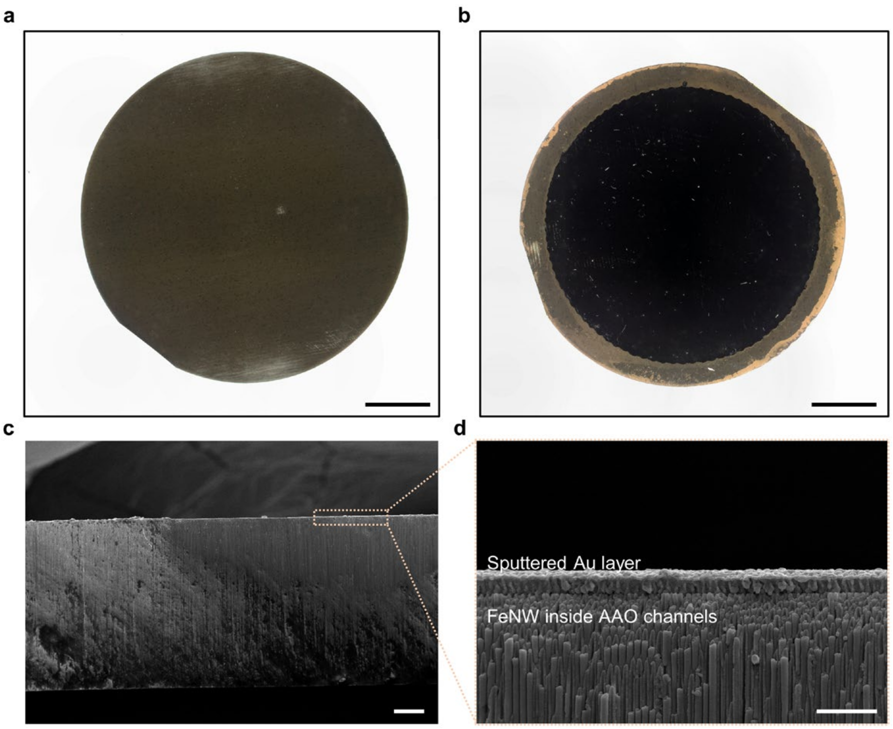

**Supplementary Fig. 18 | Anodic aluminium oxide templates after electrodeposition. a**,**b**, Optical images of the electrodeposited templates. **c**,**d**, Cross-sectional scanning electron micrographs showing the sputtered Au layer and FeNWs inside the template channels. Scale bars, 5 mm (**a**,**b**), 10 µm (**c**) and 1 µm (**d**).

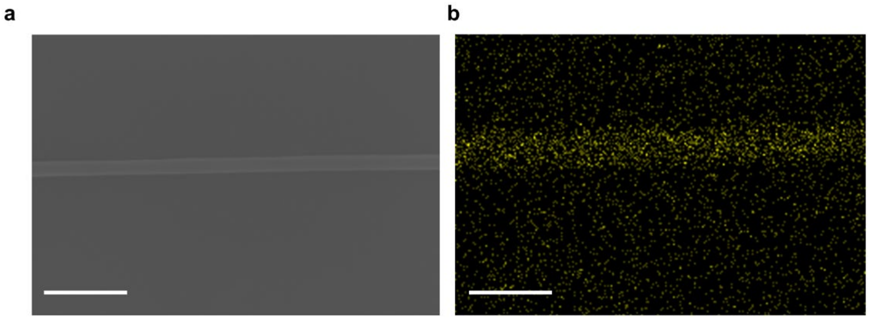

**Supplementary Fig. 19 | Morphology and elemental composition of isolated FeNWs. a**, Scanning electron micrograph of an isolated FeNW. **b**, Corresponding elemental map of Fe. Scale bars, 0.5 μm.

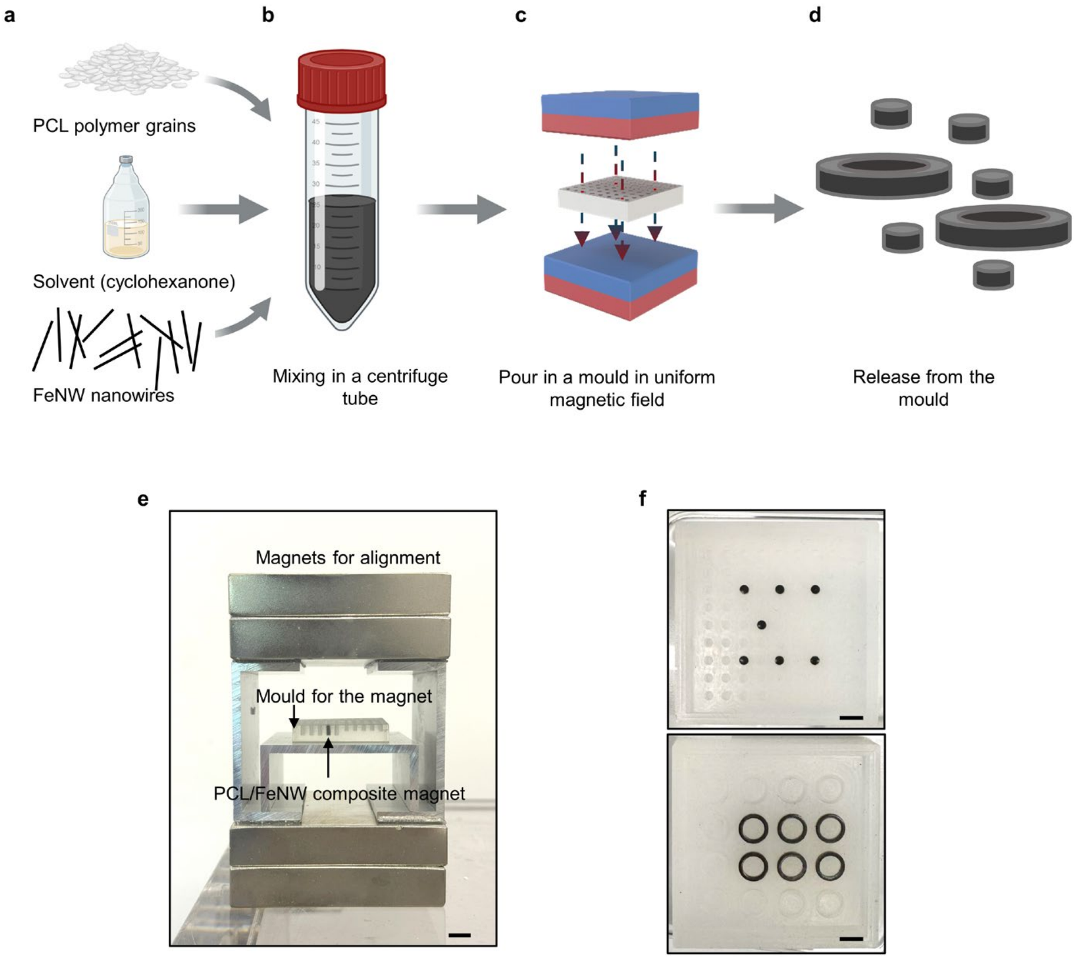

**Supplementary Fig. 20 | Fabrication of FeNW/PCL composite magnets. a–d**, FeNWs and PCL are mixed in solvent, cast into a mould, aligned in a uniform external magnetic field during solvent evaporation and released after solidification. **e**, Photograph of the mould positioned between two plate magnets for magnetic-field-assisted alignment. **f**, Representative cylindrical and annular FeNW/PCL composite magnets. Scale bars, 10 mm (**e**) and 5 mm (**f**). Illustrations in (**a**,**b**) were created with BioRender.com (https://biorender.com).

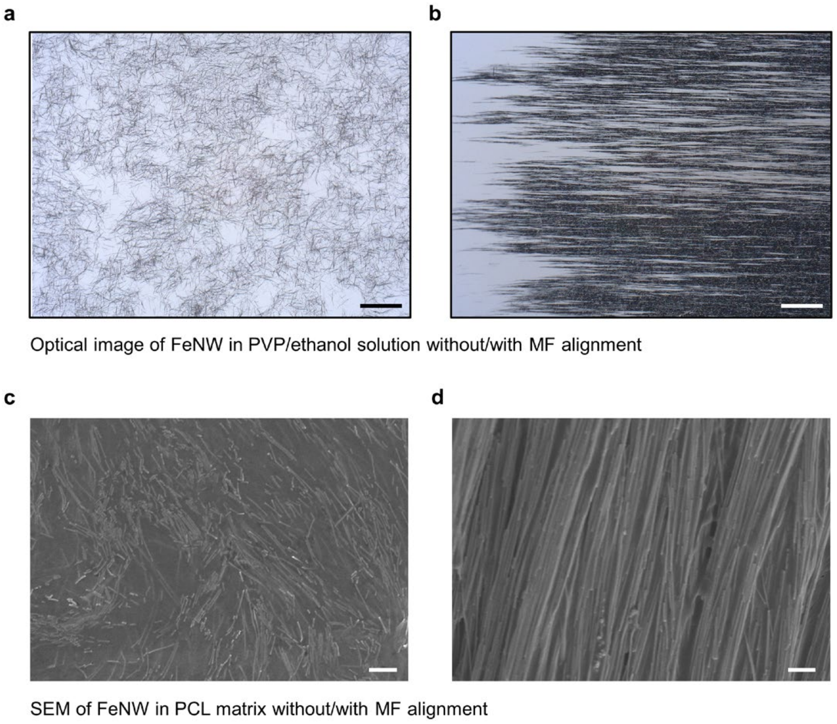

**Supplementary Fig. 21 | Magnetic-field-induced alignment of FeNWs. a**,**b**, Optical images of FeNWs dispersed in PVP/ethanol without and with an alignment field. **c**,**d**, Scanning electron micrographs of FeNWs in PCL prepared without and with field-assisted alignment. Scale bars, 100 μm (**a**,**b**) and 2 μm (**c**,**d**).

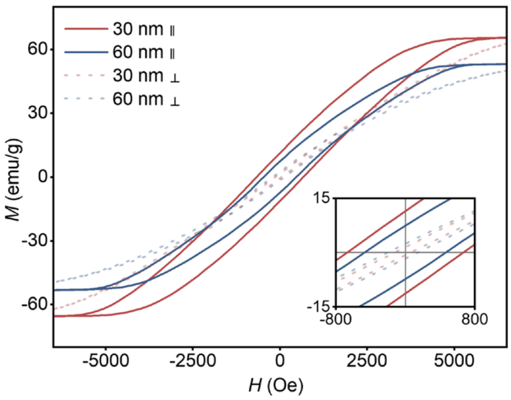

**Supplementary Fig. 22 | Diameter-dependent magnetic properties of FeNWs.** Hysteresis loops measured parallel and perpendicular to the nanowire axis for FeNWs with diameters of 30 and 60 nm. The inset shows the magnified low-field region. The 30-nm FeNWs exhibit higher coercivity and remanence than the 60-nm FeNWs along the nanowire axis.

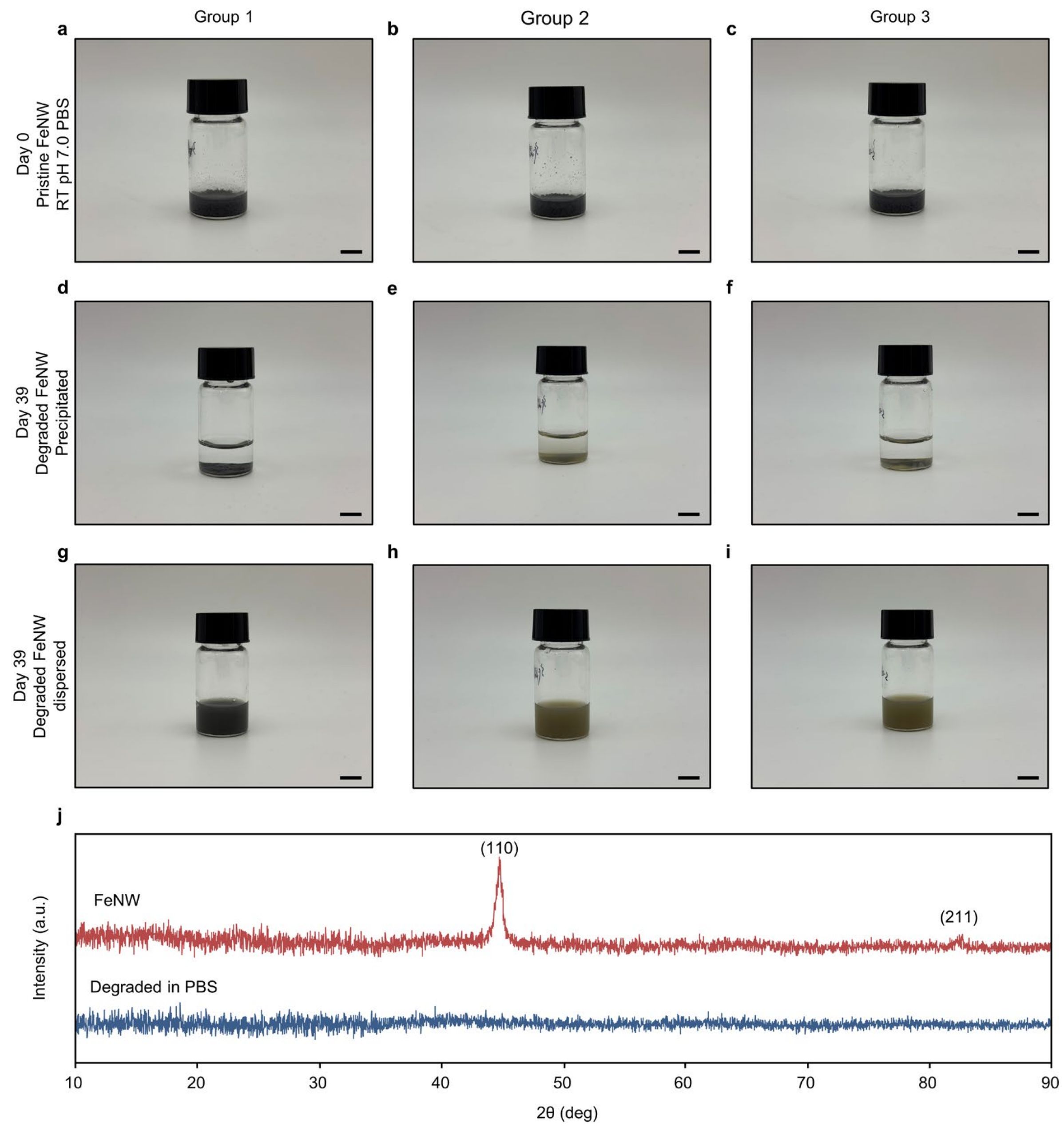

**Supplementary Fig. 23 | Degradation of FeNWs in phosphate-buffered saline. a–c**, Pristine FeNWs (black) from three independent samples on day 0. **d–f**, Precipitated degradation products (yellow-brown) on day 39. **g–i**, The same products after redispersion. **j**, X-ray diffraction patterns of pristine and degraded FeNWs. Scale bars, 1 cm.

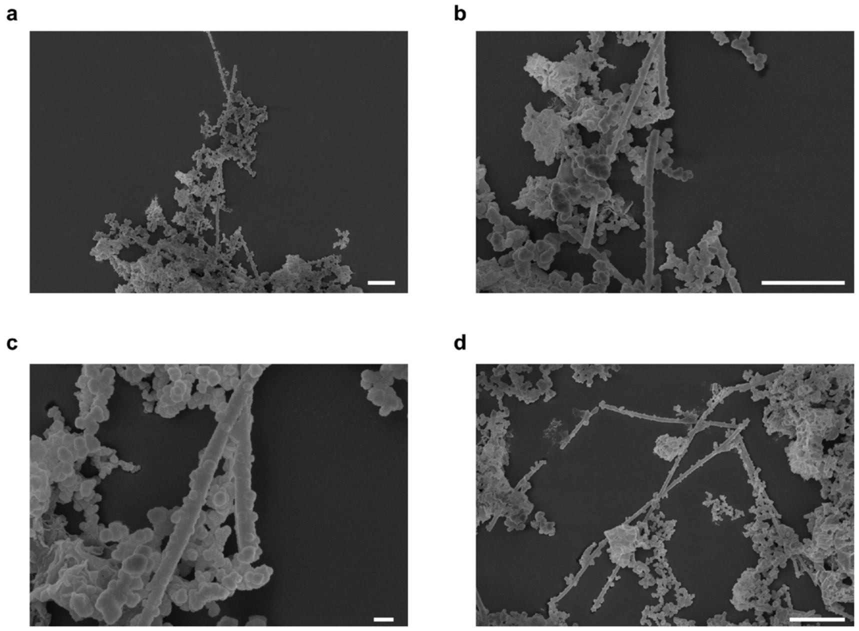

**Supplementary Fig. 24 | Morphology of degraded FeNWs.** Scanning electron micrographs showing degradation products and residual nanowire structures after immersion in phosphate-buffered saline. Scale bars, 2 µm (**a**,**b**,**d**) and 200 nm (**c**).

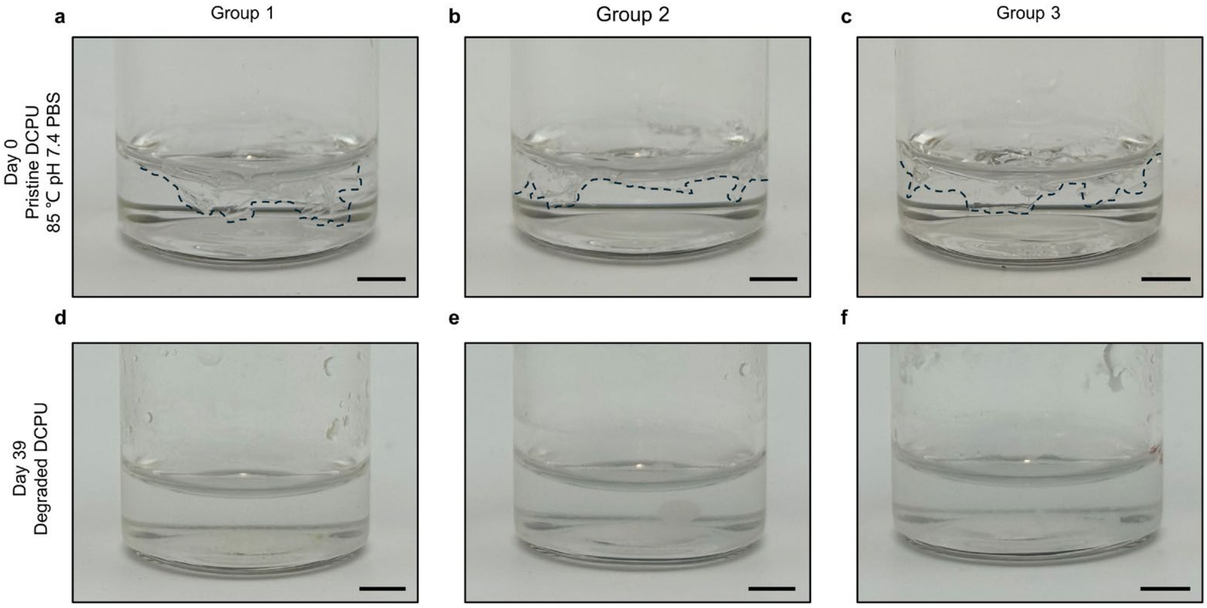

**Supplementary Fig. 25 | Degradation of b-DCPU in phosphate-buffered saline. a**–**c**, Pristine b-DCPU films from three independent groups on day 0, with clearly distinguishable film boundaries indicated by dashed outlines. **d**–**f**, Corresponding samples after 39 days at 85 ℃, with no discernible film boundaries or intact film shapes remaining, indicating substantial degradation. Scale bars, 5 mm.

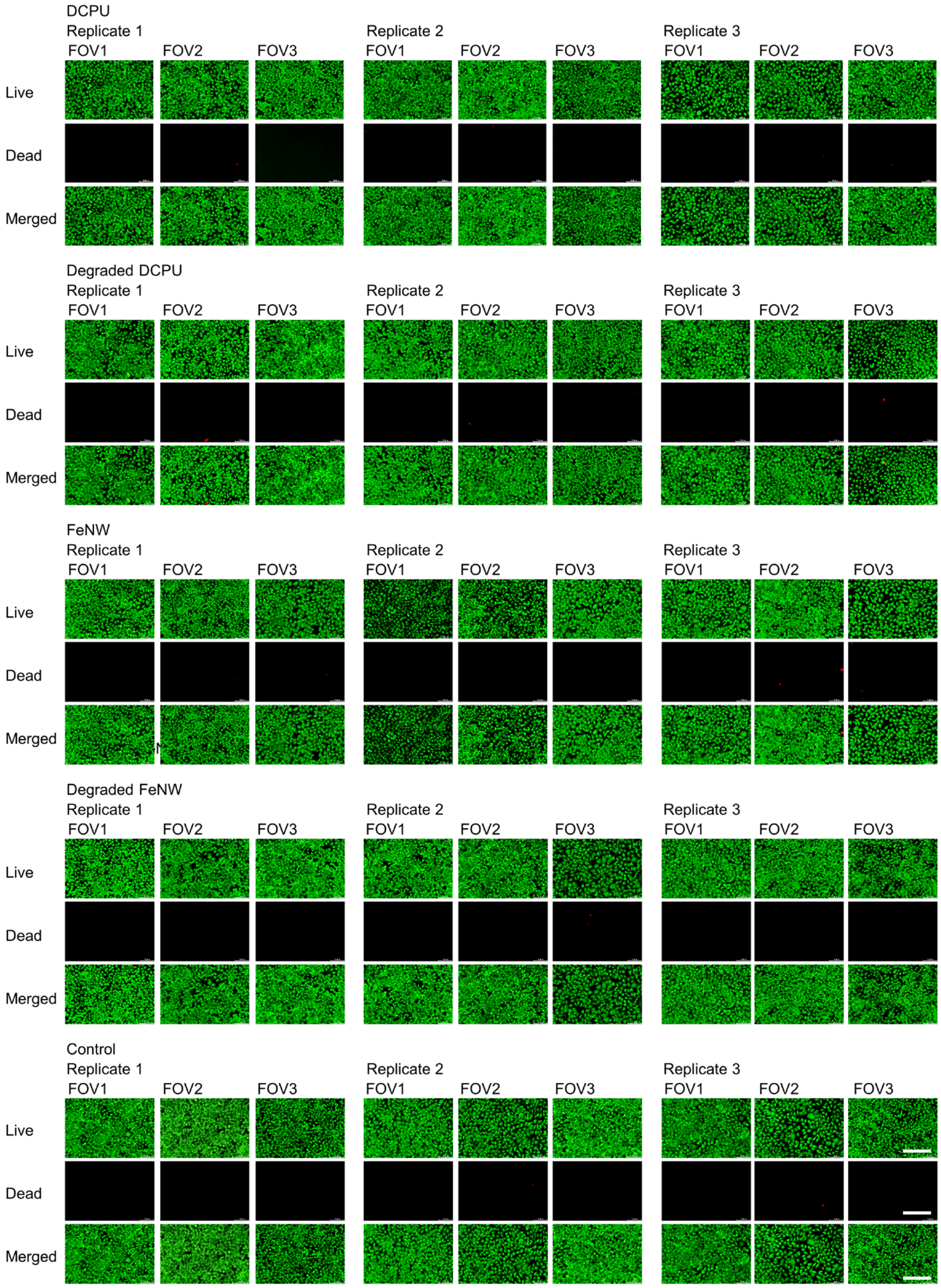

561

562 **Supplementary Fig. 26 | Live/dead staining after exposure to bioresorbable materials.**
563 Representative fluorescence images of cells exposed to b-DCPU, degraded b-DCPU, FeNWs
564 and degraded FeNWs, together with untreated controls. Three replicates and three fields of
565 view are shown for each condition. Scale bars, 200 μm.

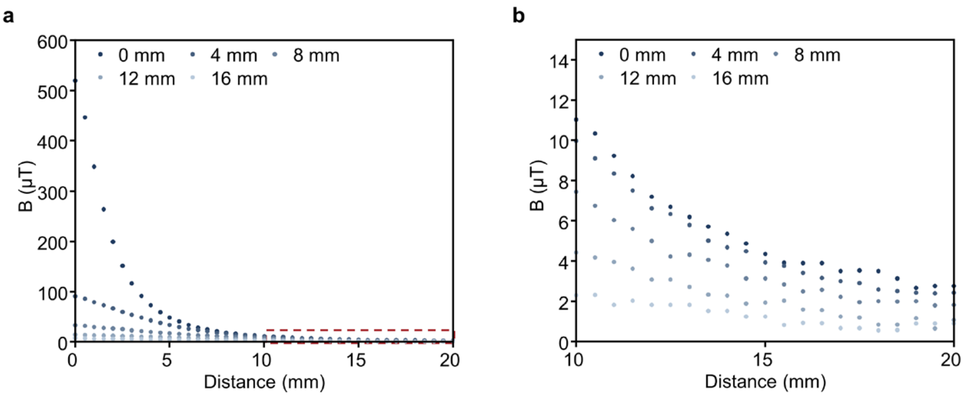

**Supplementary Fig. 27 | Distance dependence of the BioBeacon-generated magnetic field.** **a**, MF magnitude as a function of lateral distance at vertical separations of 0 to 16 mm. **b**, Magnified view of the region indicated by the dashed box in **a**, at lateral distances of 10 to 20 mm.

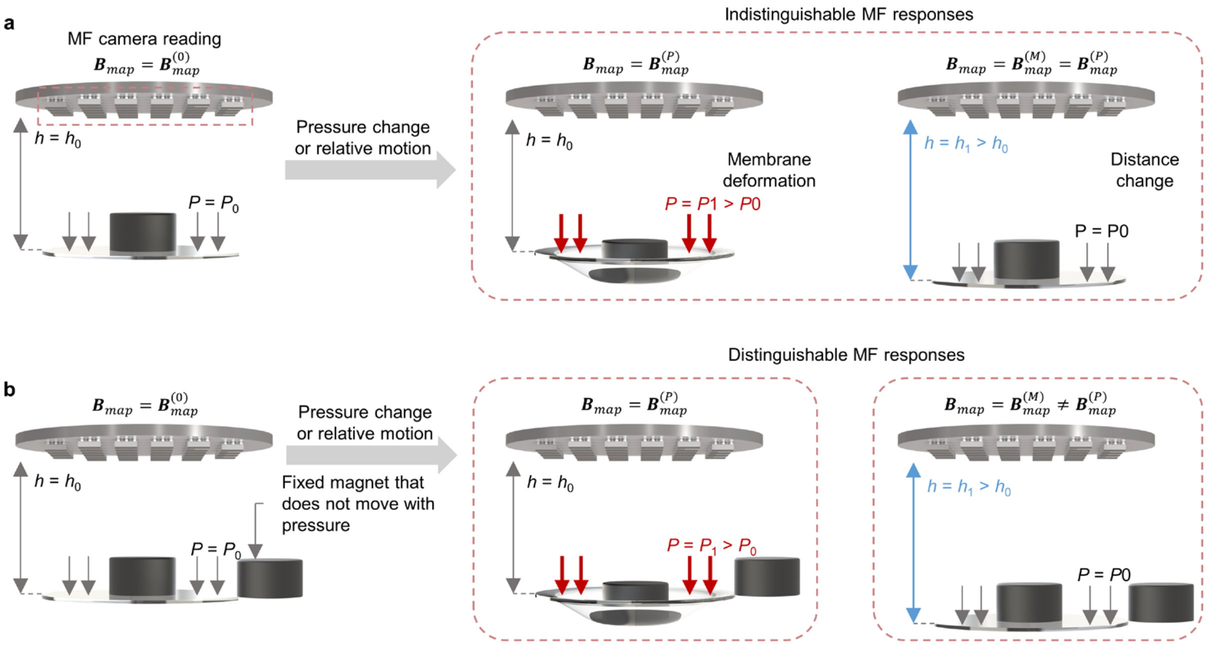

**Supplementary Fig. 28 | Physical basis of the dual-magnet sensing design. a**, In a single-magnet configuration, pressure-induced displacement and rigid-body motion can produce indistinguishable magnetic-field responses. **b**, A fixed second magnet provides an internal reference, enabling the two processes to produce distinguishable spatial magnetic-field patterns.

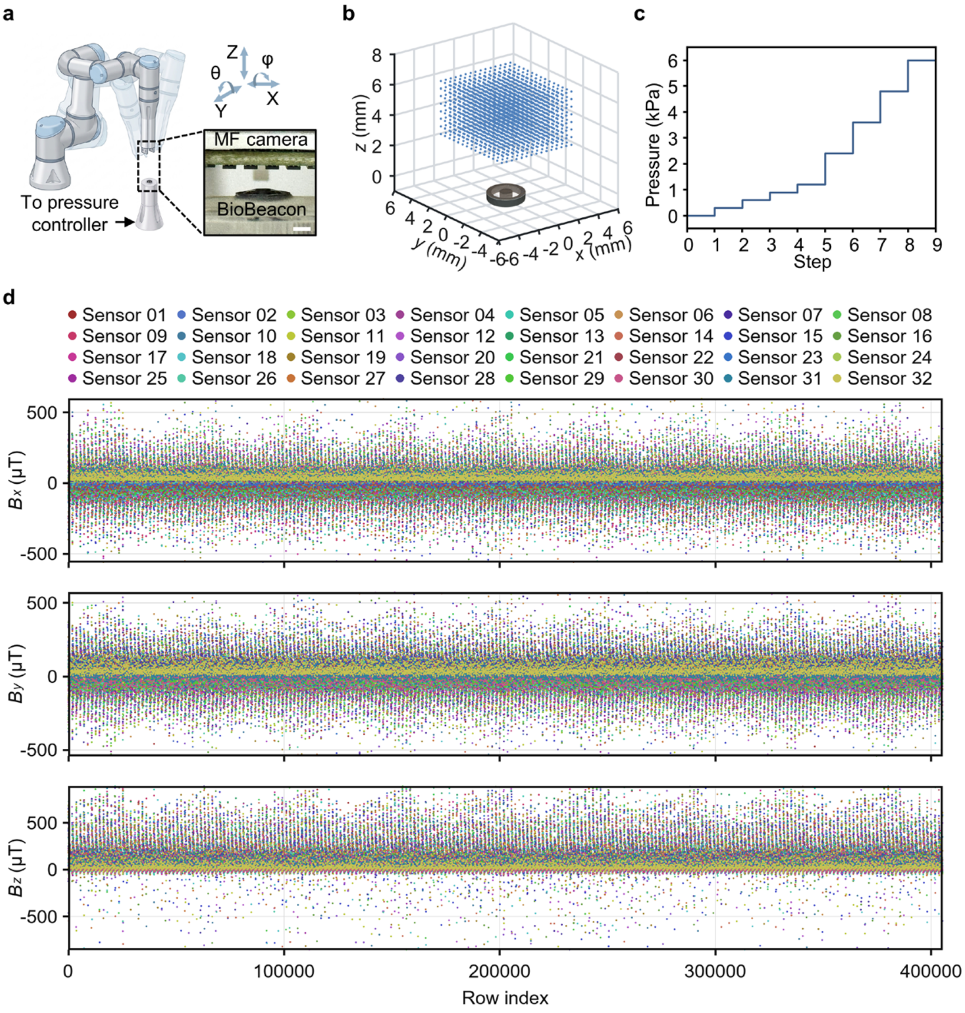

**Supplementary Fig. 29 | Collection of the pose–pressure calibration dataset. a**, Setup for pose–pressure calibration; inset, positions of BioBeacon and camera. **b**, Three-dimensional translational sampling grid around BioBeacon-PA. **c**, Applied pressure sequence. **d**, *x*-, *y*- and *z*-axis magnetic-field readings from the 32 Hall sensors during data collection. Scale bar, 2 mm (**a**, inset)

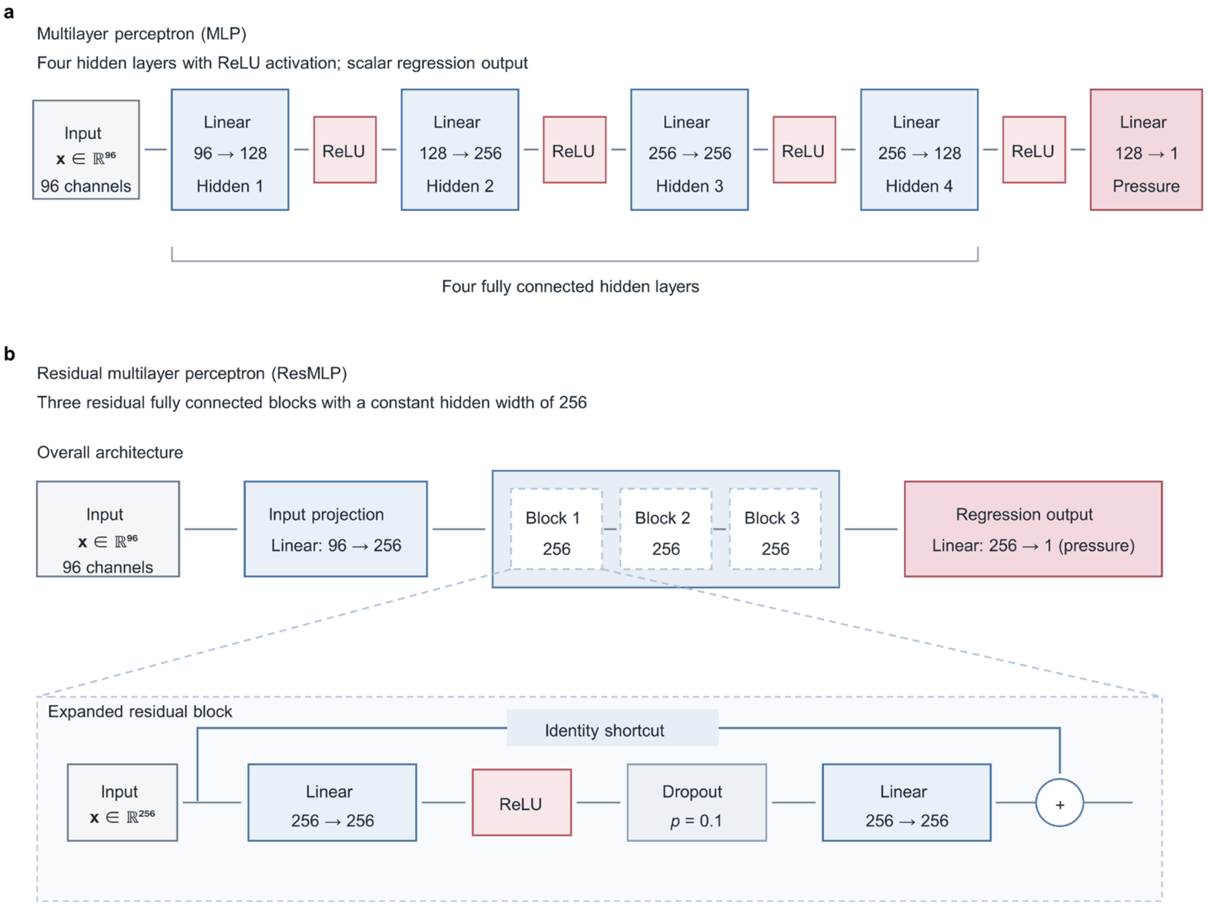

**Supplementary Fig. 30 | Neural-network architectures for pressure decoding. a**, Multilayer perceptron with four fully connected hidden layers. **b**, Residual multilayer perceptron comprising an input projection, three residual blocks and a scalar regression output.

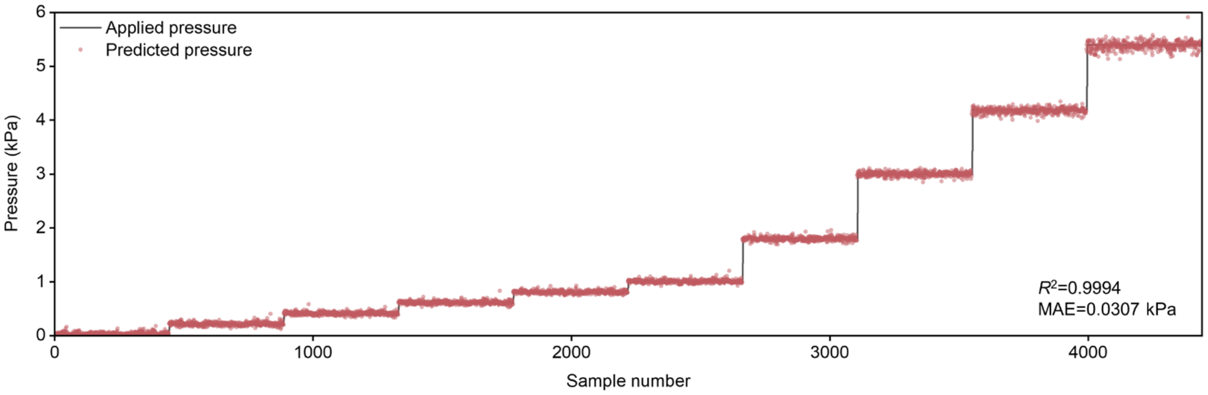

**Supplementary Fig. 31 | Pressure decoding on the independent validation dataset.** Predicted pressures closely follow the applied pressures across the tested range ($R^2$ = 0.9994, MAE = 0.031 kPa).

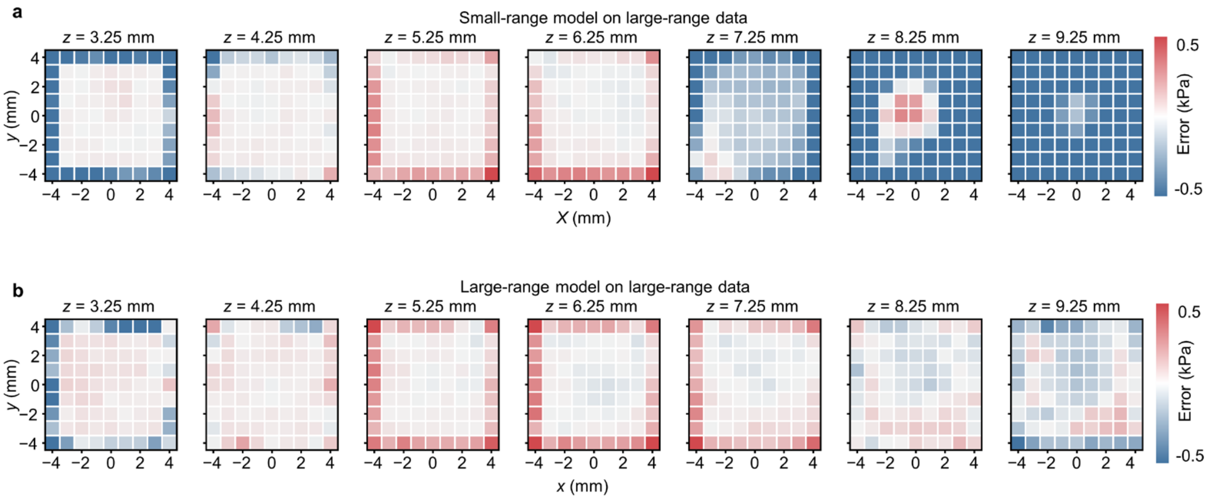

**Supplementary Fig. 32 | Effect of the vertical calibration range on pressure decoding. a**, Pressure-decoding error maps across seven vertical planes for the small-range model evaluated on the extended-range dataset. Errors increase markedly in the final three planes ($z = 7.25$–$9.25$ mm), which extend beyond the vertical calibration range. **b**, Corresponding error maps for the large-range model, showing substantially reduced errors in these three planes after expansion of the vertical calibration range.

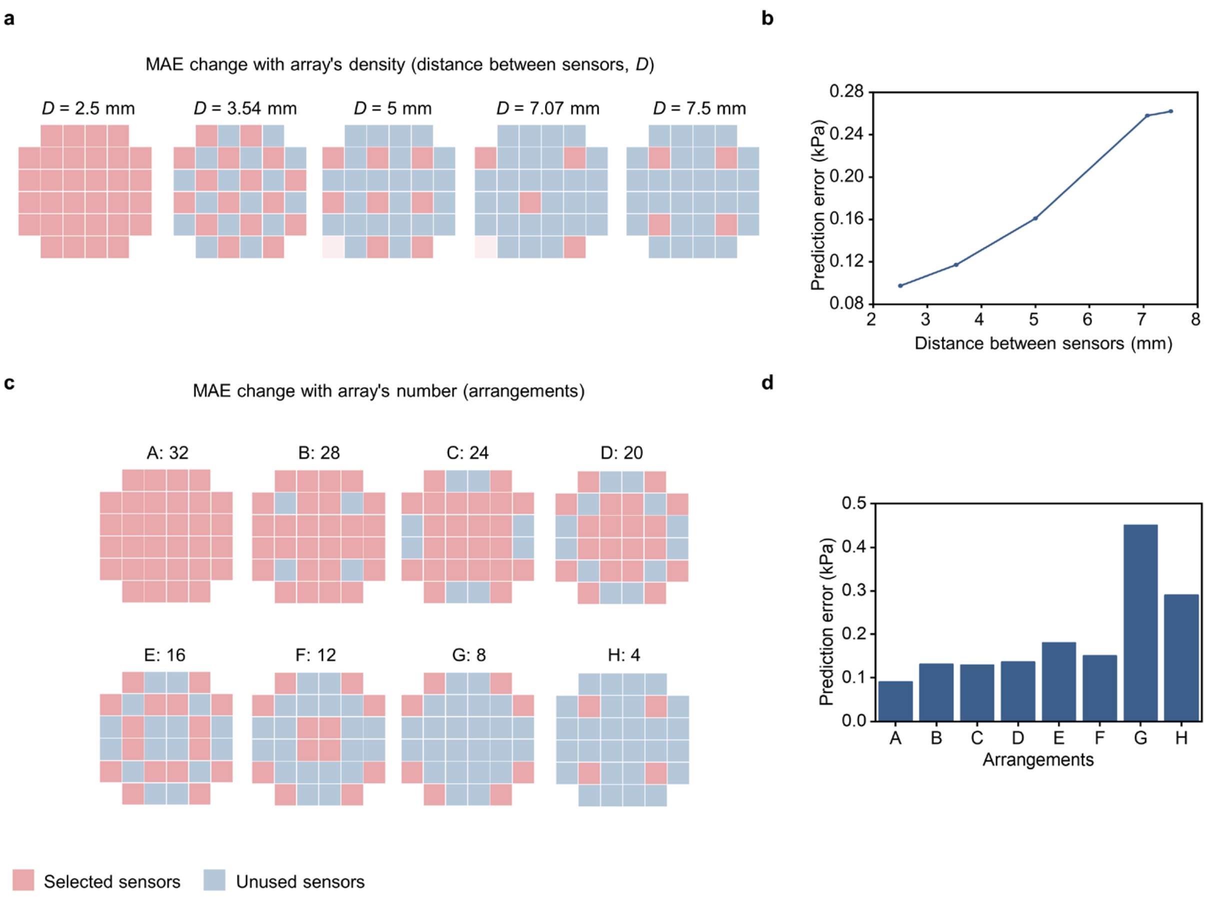

**Supplementary Fig. 33 | Effects of Hall-sensor spacing and number on decoding accuracy.** **a**, Array configurations with different sensor spacings. **b**, Prediction error as a function of sensor spacing. **c**, Array configurations containing 4–32 sensors. **d**, Prediction errors for the corresponding configurations. Increasing the sensor spacing or excessively reducing the sensor number generally increases the decoding error.

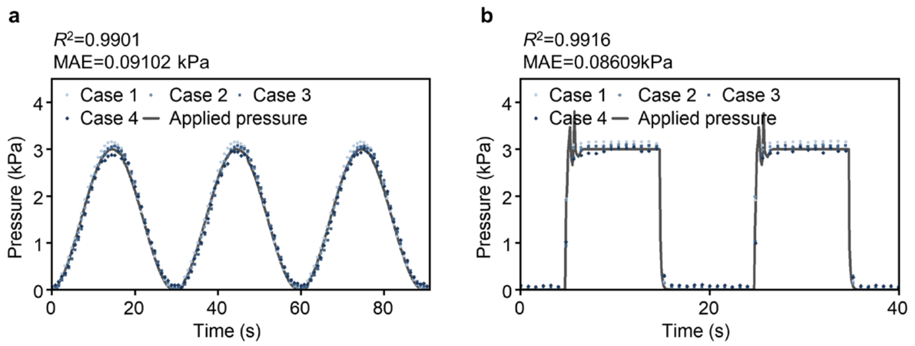

**Supplementary Fig. 34 | Pressure decoding at different relative camera–implant poses.** **a**,**b**, Predicted pressures for four representative poses during sinusoidal (**a**) and square-wave (**b**) pressure inputs. The predictions closely track the applied pressures across all four poses; overall $R^2$ and MAE values are indicated for each pressure waveform.

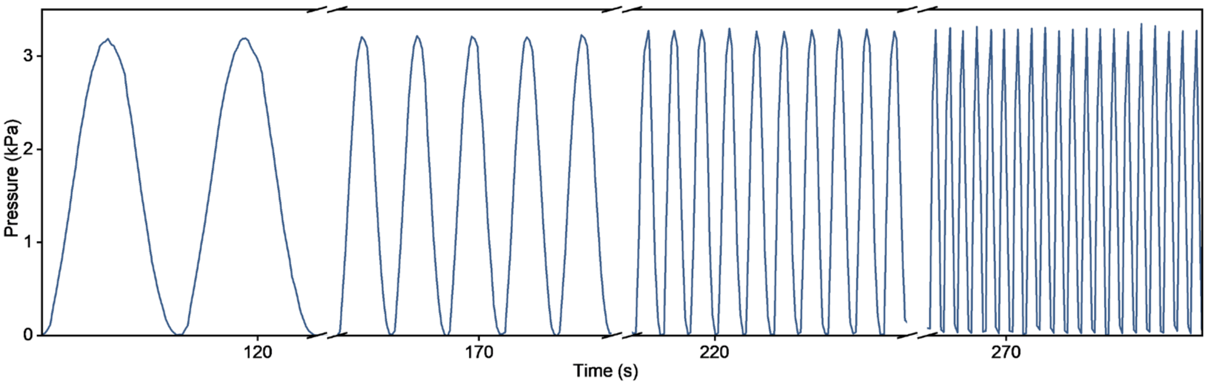

**Supplementary Fig. 35 | Dynamic pressure sensing at different input frequencies.** Decoded pressure traces under sinusoidal inputs with progressively increasing frequencies. The system preserves the complete pressure waveform over the tested frequency range of 0.2–2 Hz without evident waveform distortion.

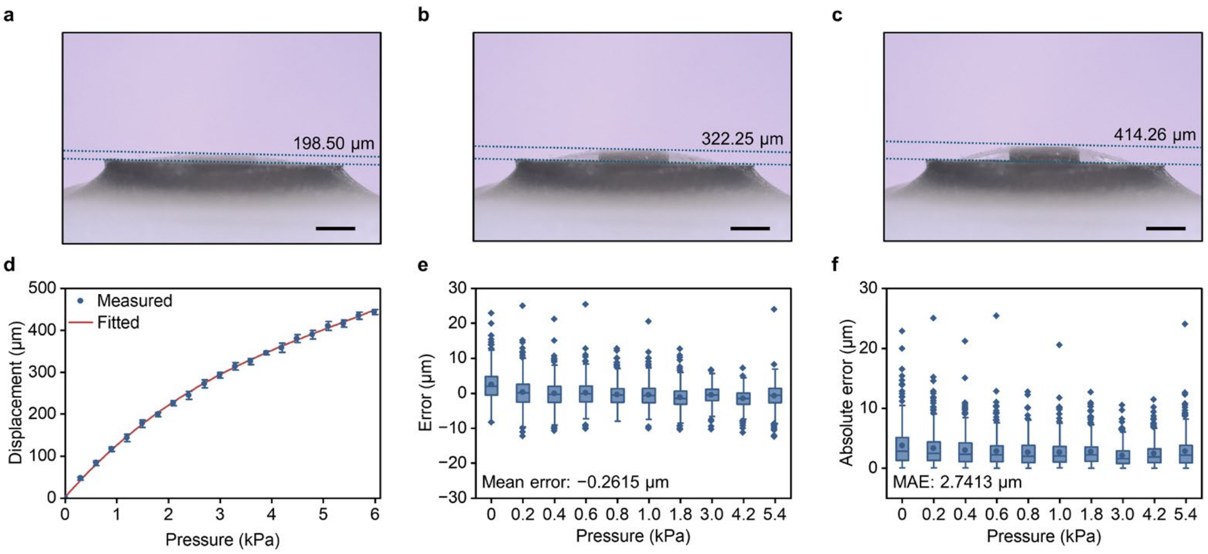

**Supplementary Fig. 36 | Equivalent displacement estimation accuracy of BioBeacon-PA.** **a**–**c**, Representative optical micrographs used to measure the axial displacement of the magnet-bearing membrane at three applied pressure levels. **d**, Measured pressure–displacement relationship and polynomial fit. Points show the mean, and error bars represent ±s.d. from three independent pressure-loading experiments on the same device ($n$ = 3 experiments). **e**, Signed errors in equivalent displacement estimation at different pressure levels ($n$ = 444 validation samples per pressure level), with the mean error indicated. **f**, Pressure-specific absolute errors in equivalent displacement estimation ($n$ = 444 validation samples per pressure level), with the overall mean absolute error (MAE) indicated. For (**e**) and (**f**), boxes span the 25th–75th percentiles, centre lines indicate medians, circles indicate means, whiskers extend to the most extreme values within 1.5 times the interquartile range, and diamonds indicate outliers. Scale bars, 1 mm.

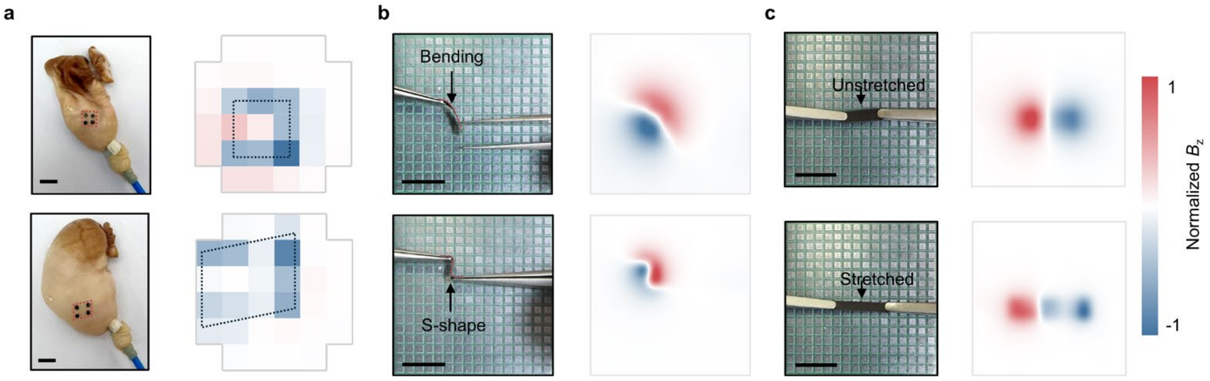

**Supplementary Fig. 37 | Deformation-induced magnetic-field changes of BioBeacon-S and BioBeacon-B. a**, Photographs and normalized $B_z$ maps of BioBeacon-S on an ex vivo rabbit stomach before (top) and after (bottom) inflation-induced strain and shear. **b**, Photographs and smoothed normalized $B_z$ maps of BioBeacon-B bent into curved (top) and S-shaped (bottom) configurations. **c**, Photographs and smoothed normalized $B_z$ maps of longitudinally magnetized BioBeacon-B in the unstretched (top) and stretched (bottom) states. Scale bars, 10 mm.

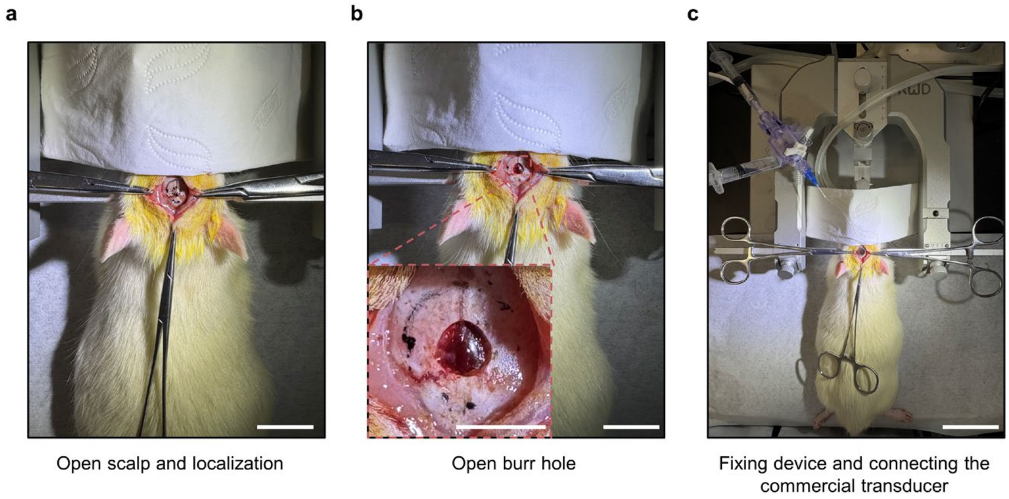

**Supplementary Fig. 38 | Craniotomy and connection to the commercial pressure transducer.** **a**, Scalp opening and surgical-site localization. **b**, Cranial opening and magnified view of the burr hole. **c**, Fixation of the animal and connection of the pressure-transmission line to the wired transducer. Scale bars, 2 cm (**a** and main image in **b**), 5 mm (inset in **b**) and 5 cm (**c**).

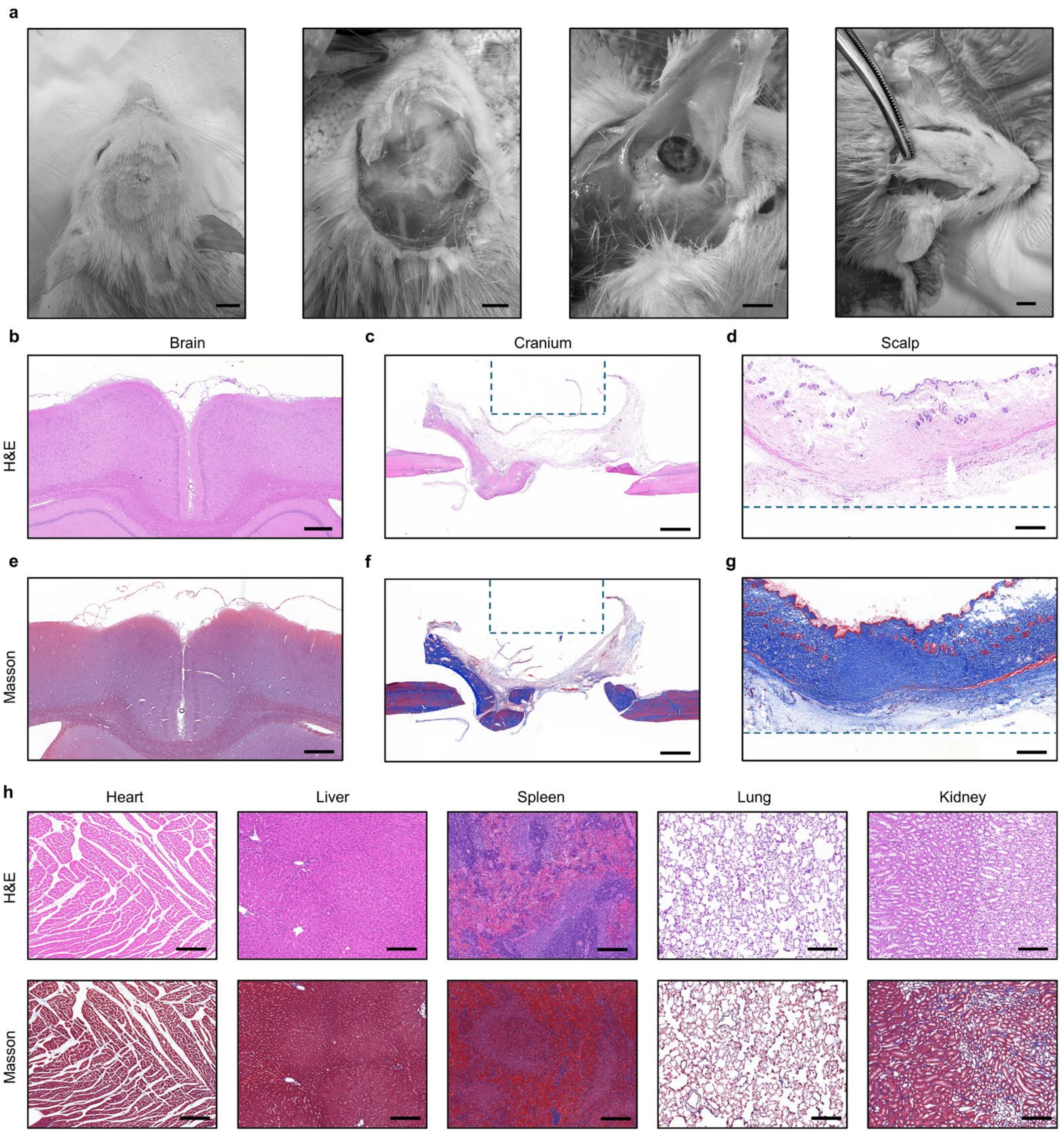

**Supplementary Fig. 39 | Histological evaluation after BioBeacon implantation. a**, Gross photographs of the tissue dissection process. **b–g**, Haematoxylin and eosin and Masson's trichrome staining of the brain, cranium and scalp. Blue dashed lines in (**c**,**d**,**f**,**g**) indicate the original tissue–BioBeacon interfaces. **h**, Corresponding staining of the heart, liver, spleen, lung and kidney. Scale bars, 5 mm (**a**), 500 µm (**b–g**) and 200 µm (**h**).

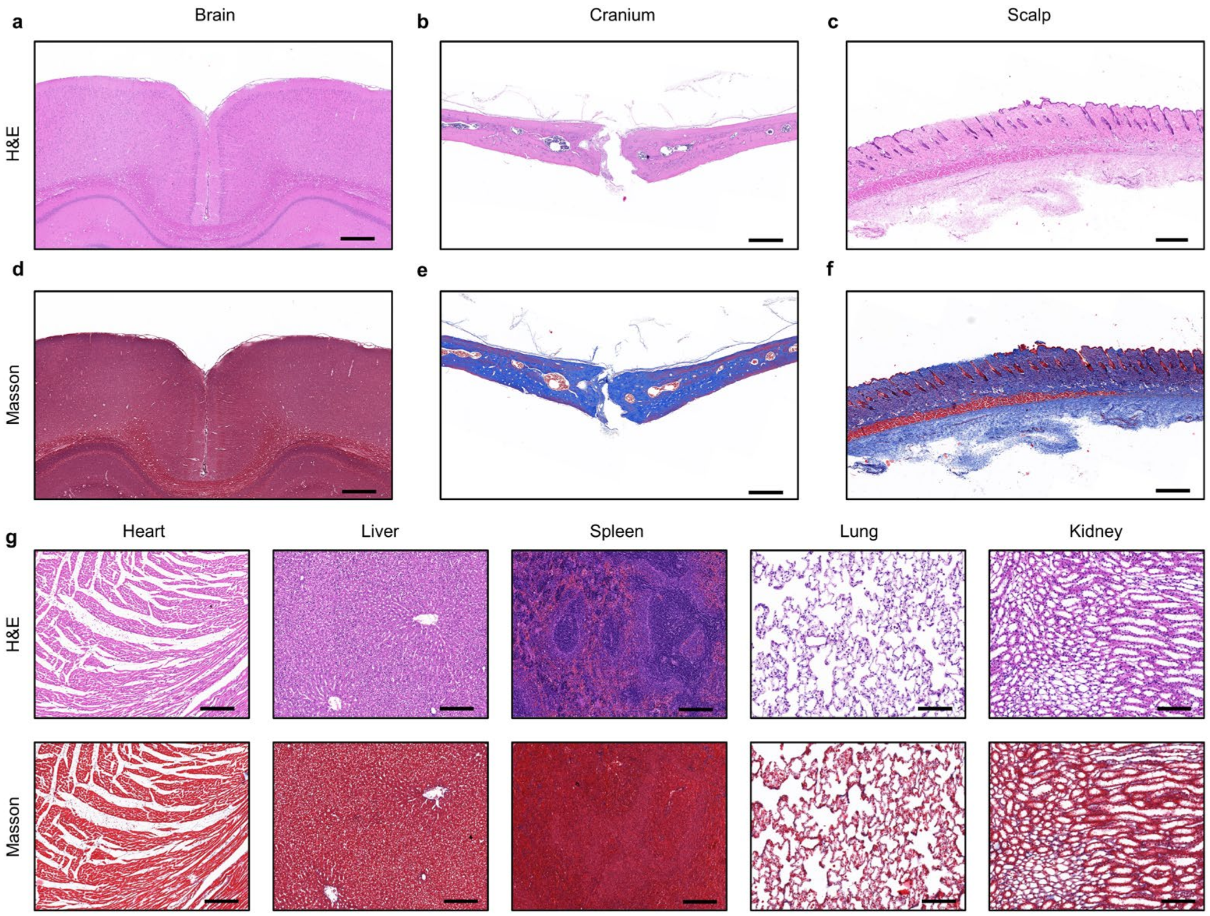

**Supplementary Fig. 40 | Histological evaluation of control animals without BioBeacon implantation.** **a**–**f**, Haematoxylin and eosin and Masson's trichrome staining of the brain, cranium and scalp. **g**, Corresponding staining of the heart, liver, spleen, lung and kidney. Scale bars, 500 μm (**a**–**f**) and 200 μm (**g**).

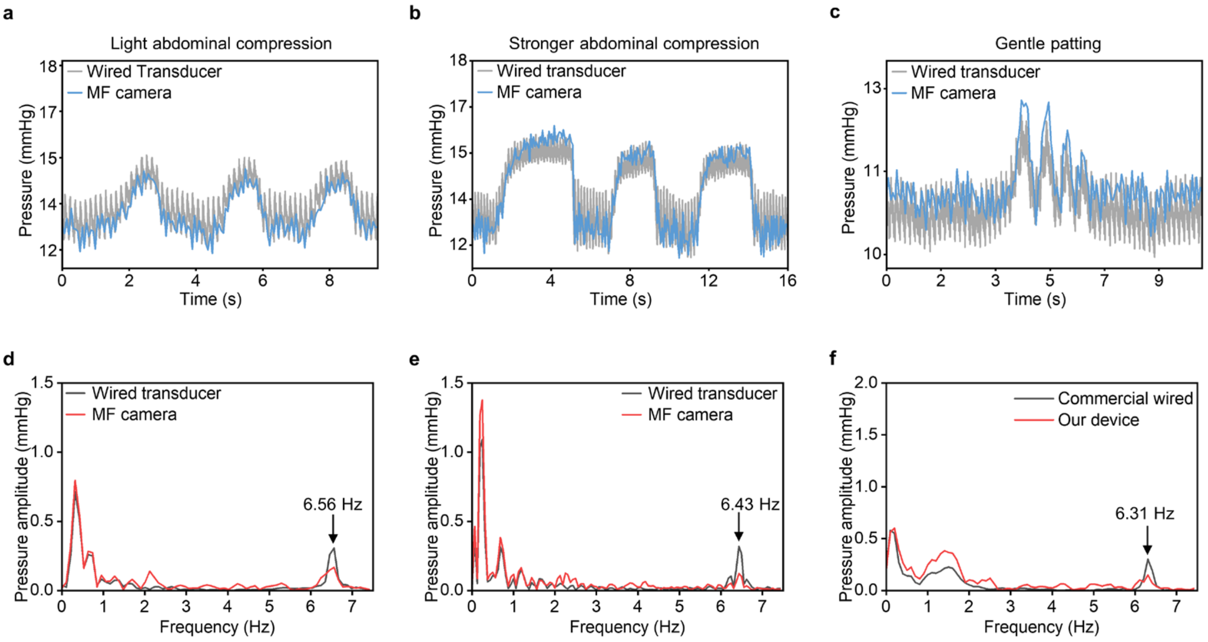

**Supplementary Fig. 41 | Comparison of decoded and commercial-reference intracranial pressure signals. a–c**, Pressure waveforms during light abdominal compression, stronger abdominal compression and gentle tapping. **d–f**, Corresponding amplitude spectra after mean removal and Hann windowing. The decoded and reference signals show matching cardiac-frequency peaks at 6.56, 6.43 and 6.31 Hz, respectively.

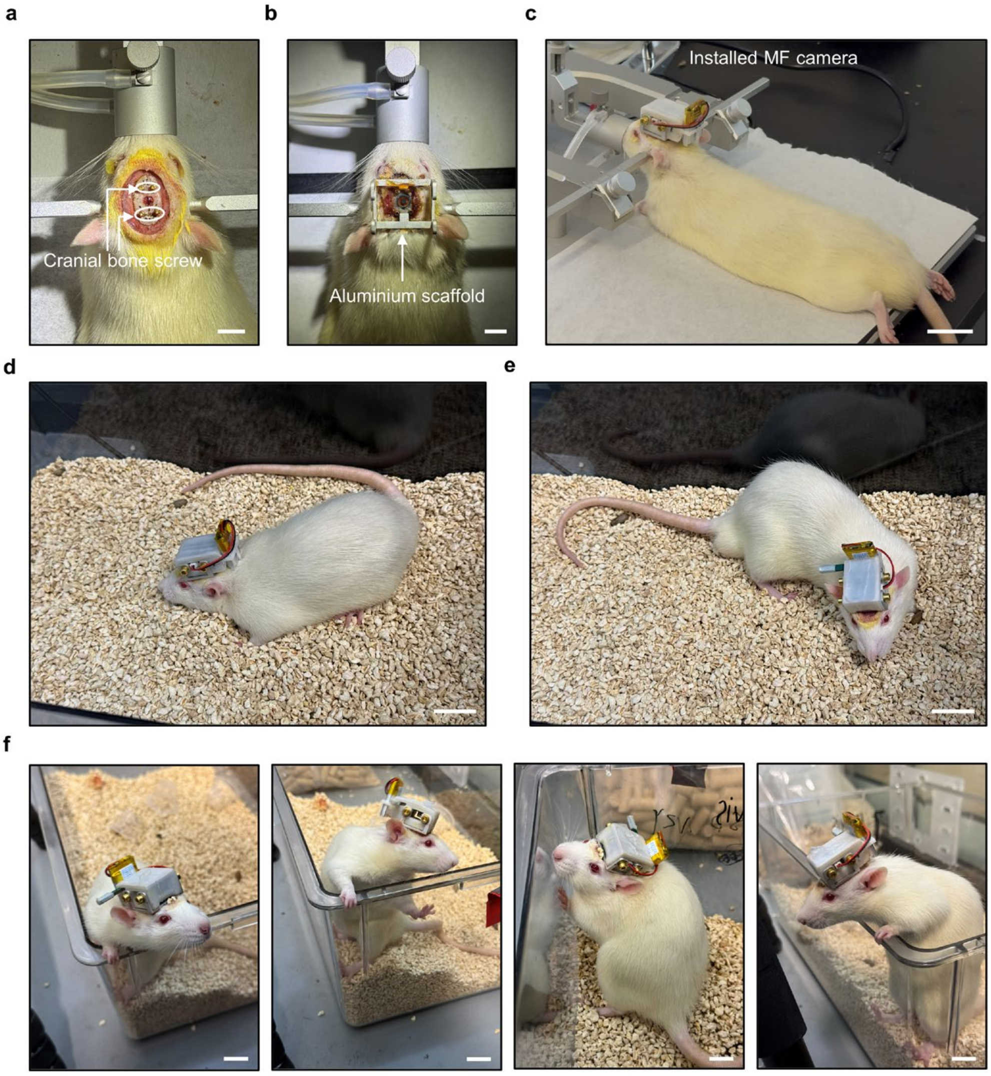

**Supplementary Fig. 42 | Installation of the wearable MF camera and free movement of the rat. a**–**c**, Cranial fixation, supporting scaffold and installed MF camera. **d**–**f**, Representative photographs of the rat moving freely while wearing the camera. Scale bars, 1 cm (**a**,**b**,**f**) and 3 cm (**c**–**e**).

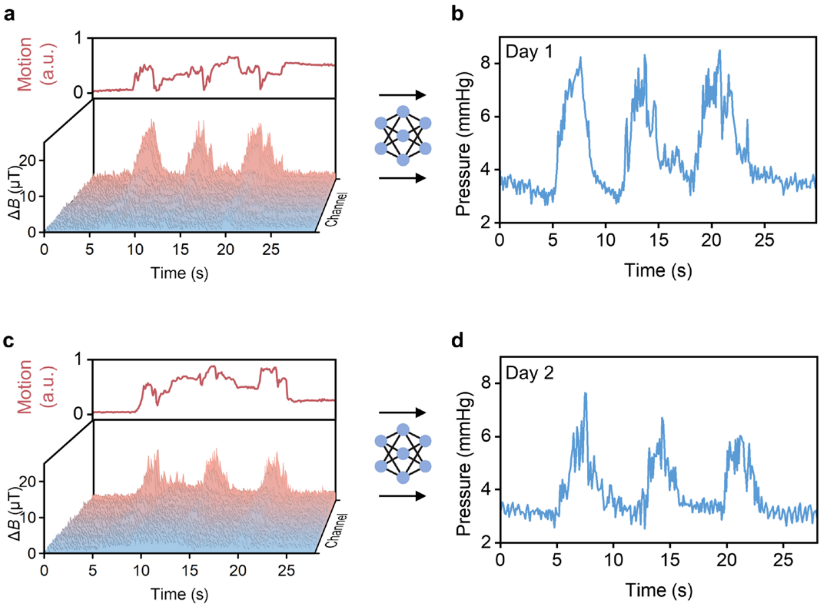

**Supplementary Fig. 43 | Intracranial pressure monitoring during tail lifting on postoperative day 1 and day 2. a**,**b**, Recordings during tail lifting on postoperative day 1, showing normalized motion intensity derived from the reference sensor (upper trace) and geomagnetically compensated multichannel MF signals (lower plot) (**a**), and corresponding decoded ICP showing elevations during tail lifting (**b**). **c**,**d**, Same as (**a**,**b**) but recorded on postoperative day 2.

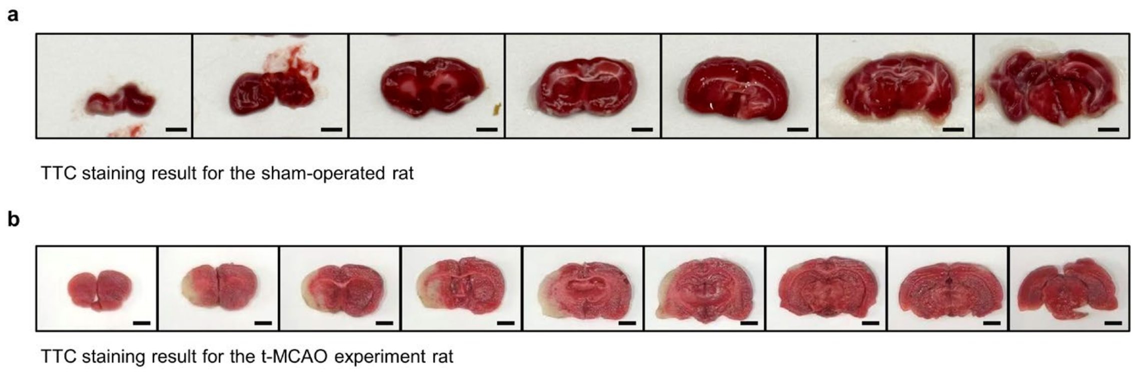

**Supplementary Fig. 44 | TTC staining after transient middle cerebral artery occlusion. a**, Brain sections from the sham-operated rat. **b**, Brain sections from the t-MCAO rat showing pale infarcted regions. Scale bars, 2.5 mm.

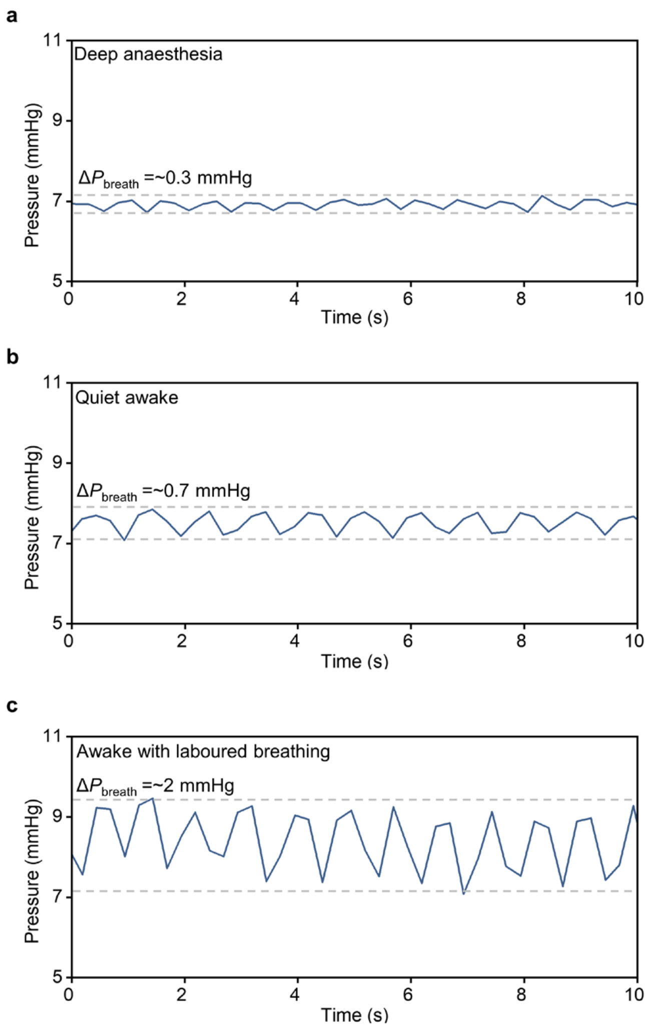

**Supplementary Fig. 45 | Respiratory-related intracranial pressure fluctuations. a–c**, Representative pressure traces under deep anaesthesia (**a**), during quiet wakefulness (**b**) and during laboured breathing (**c**), with fluctuation amplitudes of ~0.3, ~0.7 and ~2 mmHg, respectively.

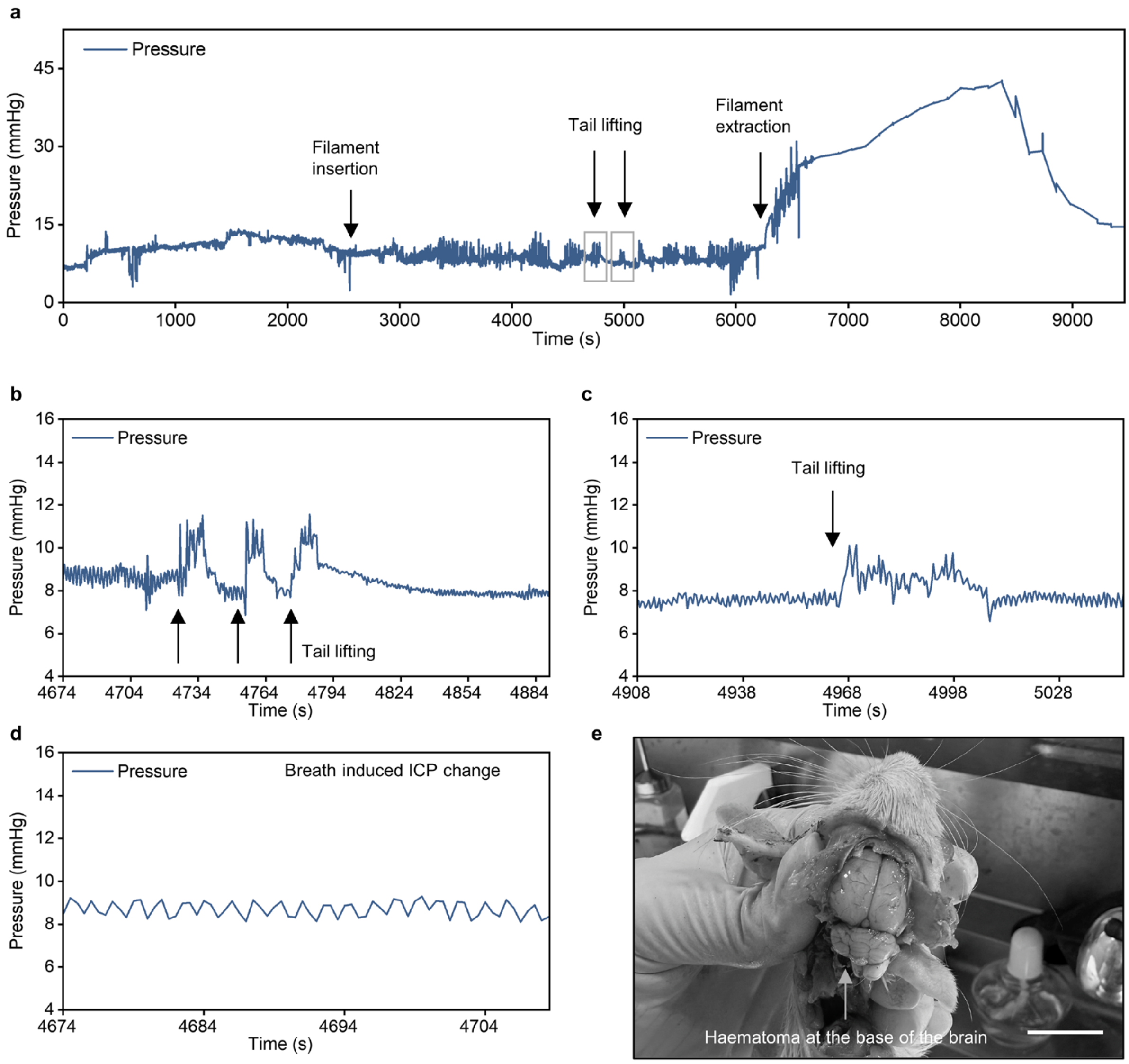

**Supplementary Fig. 46 | Intracranial pressure changes during an aborted t-MCAO procedure. a**, Pressure trace during filament insertion, tail lifting and filament extraction. **b**,**c**, Enlarged pressure responses during tail lifting. **d**, Respiratory-related pressure fluctuations. **e**, Post-mortem observation of a haematoma at the base of the brain. Scale bar, 15 mm.

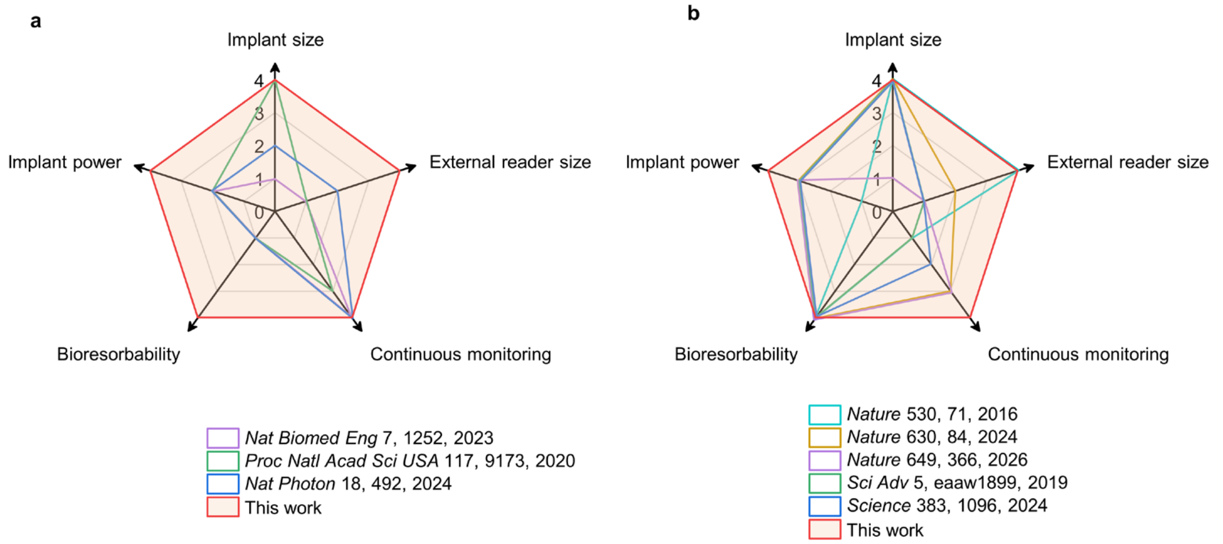

**Supplementary Fig. 47 | System-level comparison with representative implantable monitoring technologies.** **a**, Comparison with representative continuous implantable monitoring systems. **b**, Comparison with representative bioresorbable implantable sensors. The five dimensions are implant size, external reader size, continuous monitoring in freely behaving subjects, implant bioresorbability and implant power requirement. Scoring criteria are provided in Supplementary Table 4.

#### Captions for Supplementary Videos 1–6

**Supplementary Video 1 | Magnetic-field-induced alignment of iron nanowires.**
This video shows dispersed iron nanowires (FeNWs) in a polyvinylpyrrolidone (PVP)/ethanol solution rotating and aligning upon application of an external magnetic field.

**Supplementary Video 2 | *In vitro* demonstration of constant pressure sensing under multi-axis motion.**
This video demonstrates pressure decoding first under stepwise pressure changes with the MF camera stationary, and then as the MF camera translates and rotates relative to BioBeacon, with decoded pressures precisely tracking the applied constant pressure during motion.

**Supplementary Video 3 | *In vitro* demonstration of sinusoidal pressure sensing under multi-axis motion.**
This video demonstrates pressure decoding first under stepwise pressure changes with the MF camera stationary, and then as the MF camera translates and rotates relative to BioBeacon, with decoded pressures precisely tracking the applied sinusoidal pressure during motion.

**Supplementary Video 4 | Wireless intracranial pressure monitoring in a freely moving rat.**
This video shows wireless intracranial pressure (ICP) monitoring during spontaneous movement and tail lifting in an awake rat wearing the MF camera.

**Supplementary Video 5 | Behavioural deficits following transient middle cerebral artery occlusion.**
This video shows spontaneous circling during free locomotion in a rat following transient middle cerebral artery occlusion (t-MCAO), illustrating stroke-associated motor deficits.

**Supplementary Video 6 | Different respiratory states in a rat and corresponding ICP waveforms.**

This video shows breathing patterns in a rat during quiet wakefulness and during laboured breathing. The corresponding ICP recordings are shown in Supplementary Fig. 45.
